# The wheat powdery mildew resistance gene *Pm4* also confers resistance to wheat blast

**DOI:** 10.1101/2023.09.26.559489

**Authors:** Tom O’Hara, Andrew Steed, Rachel Goddard, Kumar Gaurav, Sanu Arora, Jesús Quiroz-Chávez, Ricardo Ramírez-González, Roshani Badgami, David Gilbert, Javier Sánchez-Martín, Luzie Wingen, Cong Feng, Mei Jiang, Shifeng Cheng, Susanne Dreisigacker, Beat Keller, Brande B.H. Wulff, Cristóbal Uauy, Paul Nicholson

## Abstract

Wheat blast, caused by the fungus *Magnaporthe oryzae*, threatens global cereal production since its emergence in Brazil in 1985 and recently spread to Bangladesh and Zambia. Here we demonstrate that the *AVR-Rmg8* effector, common in wheat-infecting isolates, is recognised by the gene *Pm4*, previously shown to confer resistance to specific races of *Blumeria graminis* f.sp. *tritici*, the cause of powdery mildew of wheat. We show that *Pm4* alleles differ in their recognition of different *AVR-Rmg8* alleles, and some confer resistance only in seedling leaves but not spikes making it important to select for those alleles that function in both tissues. This study has identified a gene recognising an important virulence factor present in wheat blast isolates in Bangladesh and Zambia and represents an important first step towards developing durably resistant wheat cultivars for these regions.

## Main Text

Wheat blast was first identified in Brazil in 1985 (Igarashi et al. 1986) and then spread to neighbouring countries in South America before appearing in Bangladesh in 2016 (Malaker et al. 2016) and Zambia in 2018 (Tembo et al. 2020). Wheat blast is caused by the fungus *Magnaporthe oryzae* (syn. *Pyricularia oryzae*) pathotype *triticum* (*MoT*) and is considered to pose a threat to major wheat (*Triticum aestivum*) producers including India and China, where conditions are conducive to this disease. Pathotypes of *M. oryzae* exhibit high levels of host specificity and relatively few genes in wheat have been reported to be effective against *MoT* pathotypes (Singh et al. 2021) (**Supplementary Table 1**). These include *Rmg7, Rmg8* and the 2NS translocation originating from *Aegilops ventricosa* that has been introgressed onto the short arm of wheat chromosome 2A (Singh et al. 2021). Genome-wide association studies (GWASs) of field-based resistance to wheat blast revealed that the 2NS resistance was the only potent and robust resistance in international field trials (Juliana et al. 2019; Juliana et al. 2020; He et al. 2022). Unfortunately, some *MoT* isolates are virulent on wheat varieties carrying the 2NS translocation and this resistance is not effective in all genetic backgrounds (Cruz et al. 2016; Téllez et al. 2019) making it important to identify additional sources of resistance. *Rmg7* was identified in a tetraploid wheat accession (*Triticum dicoccum*, KU120) (Tagle, Chuma, and Tosa 2015) and mapped to chromosome 2A, while *Rmg8* was identified in hexaploid wheat line S-615 and mapped to chromosome 2B (Anh et al. 2015). While *Rmg8* remains effective at higher temperatures, *Rmg7* loses its ability to confer resistance (Anh et al. 2018). It was later demonstrated that *Rmg7* and *Rmg8* both recognise the same effector *AVR-Rmg8* (Anh et al. 2018). Despite mapping these loci, no resistance gene effective against *MoT* has been cloned to date.

It has recently been reported that wheat blast isolates from Bangladesh and Zambia are part of a clonal lineage termed B71 (Latorre et al. 2023). It was shown that members of this lineage all carry *AVR-Rmg8* making identification of the corresponding resistance gene(s) an important target. We previously identified two wheat resistance genes, *Rwt3* and *Rwt4*, that acted as host specificity barriers against non-*MoT* pathotypes using isolates carrying specific effectors to screen a panel of wheat accessions (Arora et al. 2023). In this study we used *k*-mer based association of a large, whole genome shotgun-sequenced wheat diversity panel to identify the wheat gene recognising and providing resistance to isolates of *MoT* carrying *AVR-Rmg8*.

### Resistance to *AVR-Rmg8* maps to a 5.3 Mbp region on chromosome 2A of wheat cultivar SY-Mattis

We selected two isolates to identify resistance to *AVR-Rmg8*. The first isolate (Py 15.1.018) carries the *eI* allele of *AVR-Rmg8* and is virulent against cultivar Jagger and the CBFusarium ENT014 wheat line. (**Supplementary Fig. 1**), despite them both carrying the 2NS resistance (Walkowiak et al. 2020; Ferreira et al. 2020). The second isolate (NO6047+*AVR-Rmg8*), (henceforth referred to as NO6047+*AVR8*) is a derivative of isolate NO6047 that was transformed with allele *eI* of *AVR-Rmg8* under control of the *PWL2* promoter (Jensen and Saunders 2023) (**Supplementary Table 2**). NO6047 contains an alternative allele of the *AVR-Rmg8* effector, designated *eII’’’*, which is not recognised by *Rmg8* (Latorre et al. 2023). This makes NO6047 an ideal isolate to host the effector *Avr-Rmg8*. Isolate NO6047 is virulent on wheat line S-615 which carries *Rmg8* whereas isolate NO6047+*AVR8* is not virulent because of host recognition of *AVR-Rmg8* (Jensen and Saunders 2023). The resistance of any wheat accessions showing resistance to NO6047+*AVR-Rmg8* but not to NO6047 can be assumed to be due to recognition of the *AVR8-Rmg8* effector.

**Fig. 1:**
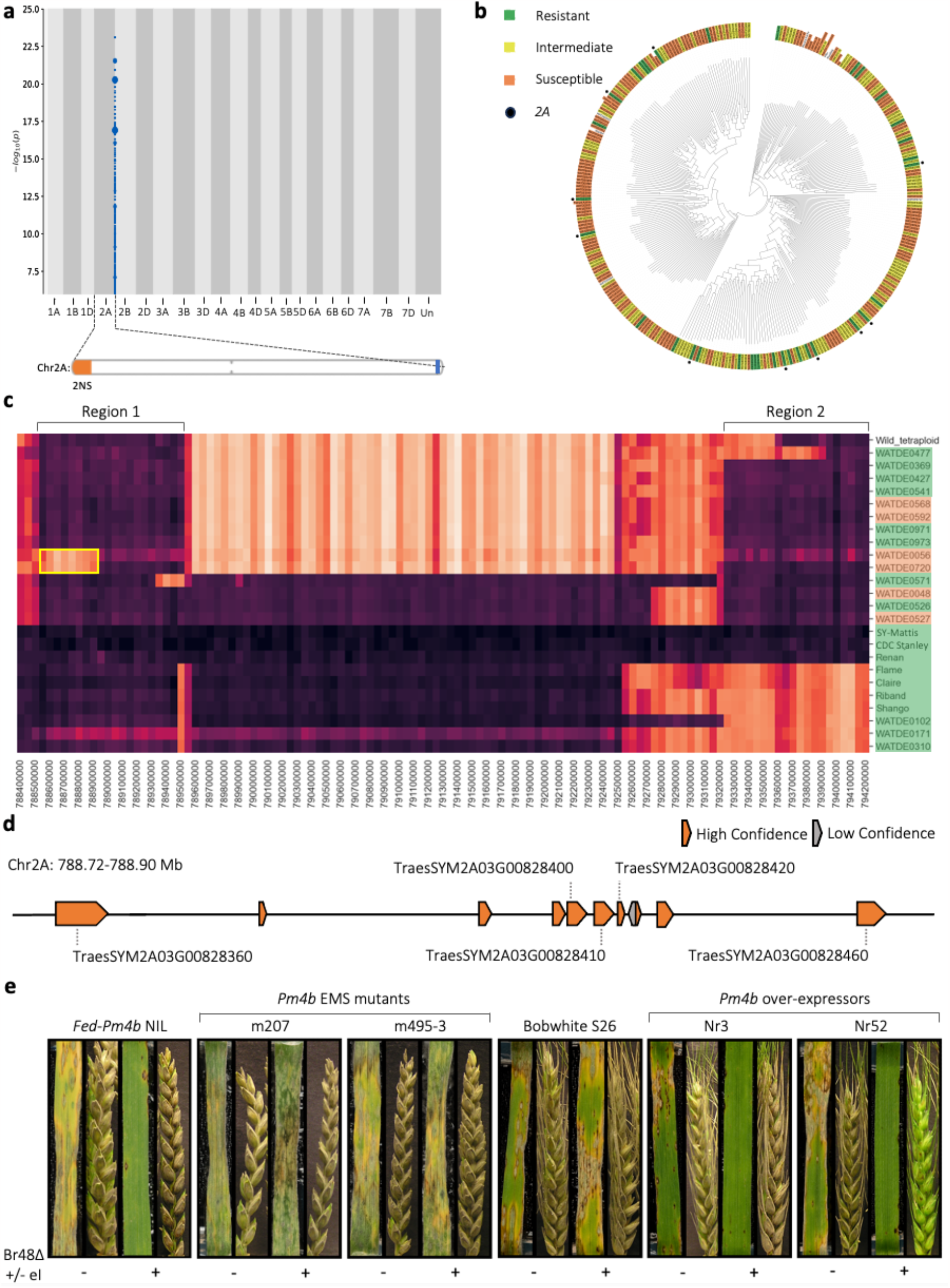
Genetic identification and validation of resistance to the wheat blast fungus effector AVR-Rmg8 by k-mer-based association mapping and haplotype analysis. **a**, *k*-mers (WATDE0310) associated with resistance to Py 15.1.018 mapped to the SY-Mattis genome. Points on the y-axis depict *k*-mers positively associated with resistance in blue. Point size is proportional to the number of *k*-mers. The association score is defined as the –log_10_ of the P value obtained using the likelihood ratio test for nested models. The ideogram shows the position of the *Ae*. ventricosa 2NS segment (orange) and the WatRenSeq 2A association (blue) on the distal ends of the short and long chromosome arms respectively. **b**, *k*-mer-based phylogeny of wheat landraces showing the phenotype of an accession after inoculation with Py 15.1.018. Phenotype of an accession after inoculation is indicated by the colour used to highlight the label of that accession (green = resistant (scores less than or equal to 3), yellow = intermediate (scores more than 3, less than 5) and orange = susceptible (scores equal to or greater than 5). Black circles indicate the presence of the chromosome 2A peak based on the WatRenSeq association plots. **c**, haplotype cluster heatmap for the chromosome 2A interval using SY-Mattis as the reference. The phenotype of an accession after inoculation with Py 15.1.018 is indicated by the colour used to highlight the label of that accession, as in (**b**). The darker the colour within a 50 kb window the more identical by state that sequence is to SY-Mattis. Among the accessions carrying the chromosome 2A interval, two regions were particularly similar to SY-Mattis, ‘Region 1’ (788,550,000 to 789,550,000) and ‘Region 2’ (793,250,000 to 794,250,000). Note that that the ‘Region 1’ haplotype block extends approximately 250 kb upstream of the 5.3 Mb 2A interval. Flame, Claire, Riband, Shango, WATDE0102, WATDE0171 and WATDE0310 were resistant but only contained ‘Region 1’. WATDE0056 and WATDE0720 were susceptible and lacked the first 400 kb of ‘Region 1’, highlighted in a yellow box. Source data are available from Zenodo under the DOI zenodo.org/record/8377152. **d**, Gene content of the 400 kb in Mattis according to the *de novo* gene models. Genes coloured in orange and grey correspond to high and low confidence genes respectively. Genes that show expression in young aerial tissue are labelled. **e**, Wheat blast detached leaf and spike assays for the *Pm4b* EMS-induced mutants of *Fed-Pm4b* and *Pm4b* over-expressors in the Bobwhite S26 background. Leaves and spikes were inoculated with *M. oryzae* isolates Br48ΔeI and Br48ΔeI+eI at 22 °C, denoted by ‘-’ and ‘+’ respectively.

We screened seedlings of a panel of 320 wheat lines including 300 landraces from the A. E. Watkins collection (Wingen et al. 2014) and wheat lines with chromosome-scale assemblies (Walkowiak et al. 2020) using the two *MoT* isolates described above (**Supplementary Table 3**). Only 13 accessions were highly resistant to the NO6047+*AVR8* isolate (score of 1.5 or less) (**Supplementary Table 4; Supplementary Fig. 2**), only ten accessions were highly resistant to Py 15.1.018 (score of 1.5 or less), and nine of these were also highly resistant to both isolates suggesting that, while resistance is rare, the majority of the resistance observed was due to the recognition of the same effector in the two isolates (**Figure 1b**; **Supplementary Table 4; Supplementary Fig. 3**). Of the cultivars with chromosome or scaffold-scale assemblies, three (SY-Mattis, CDC Stanley, Claire) were highly resistant to both isolates (but susceptible to the wildtype isolate NO6047), (**Supplementary Figs. 4 and 5**). The resistance of these three cultivars enables them to be used as references for subsequent analysis to locate and identify the causal gene.

**Fig. 2:**
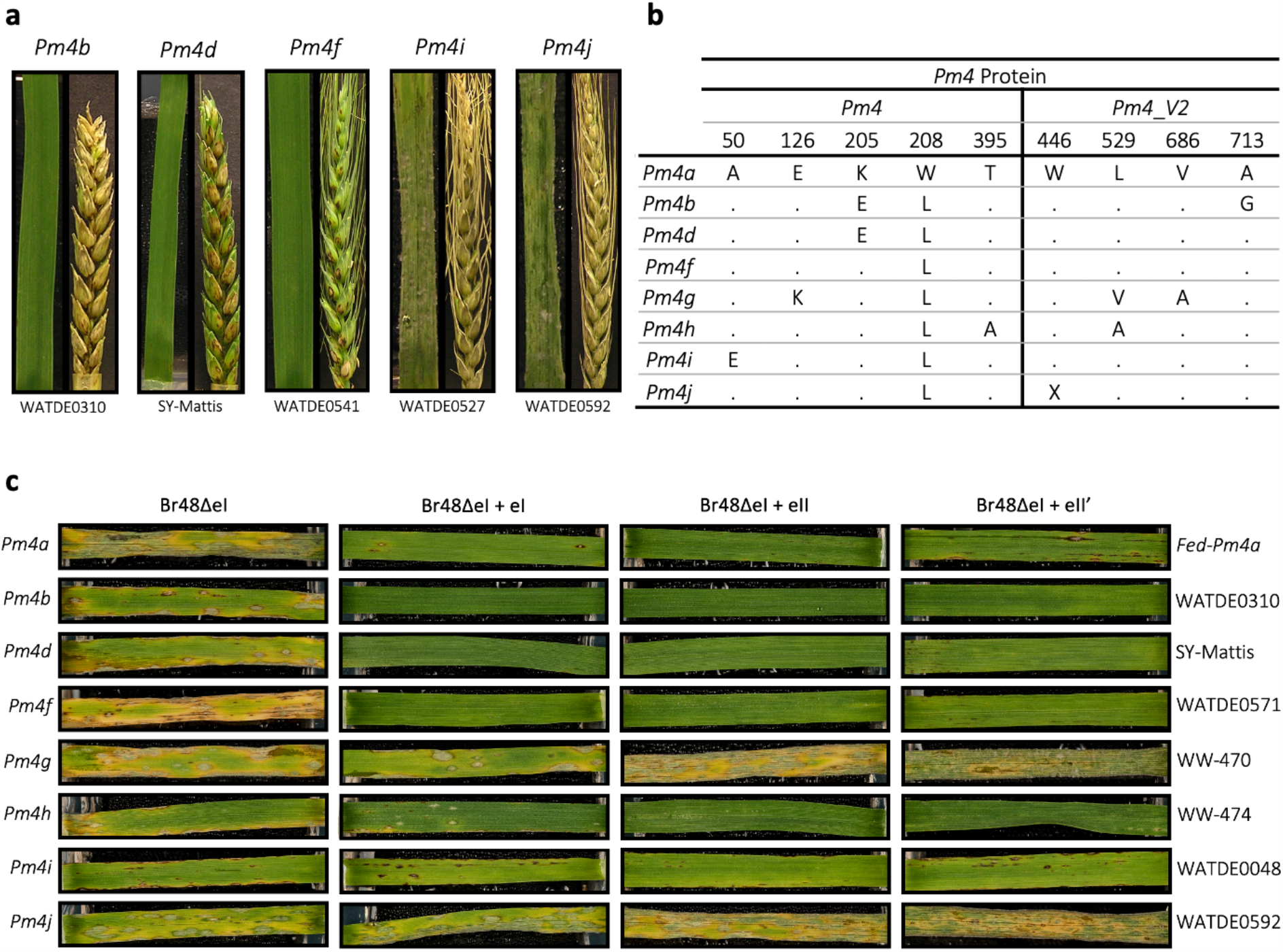
*The effect of allelic variation in Pm4 and AVR-Rmg8* on wheat blast symptoms. **a**, representative wheat blast detached leaf and spike assays for the *Pm4b, Pm4d, Pm4f, Pm4i* and *Pm4j* alleles, inoculated with Py 15.1.018 at 22 °C. **b**, protein sequence comparison of the known *Pm4* alleles. Dots represent the same amino acid present in *Pm4a*. **c**, representative wheat blast detached leaf and spike assays for the known *Pm4* alleles, inoculated with *M. oryzae* isolates Br48ΔeI, Br48ΔeI+eI, Br48ΔeI+eII and Br48ΔeI+eII’ at 22 °C.

**Fig. 3:**
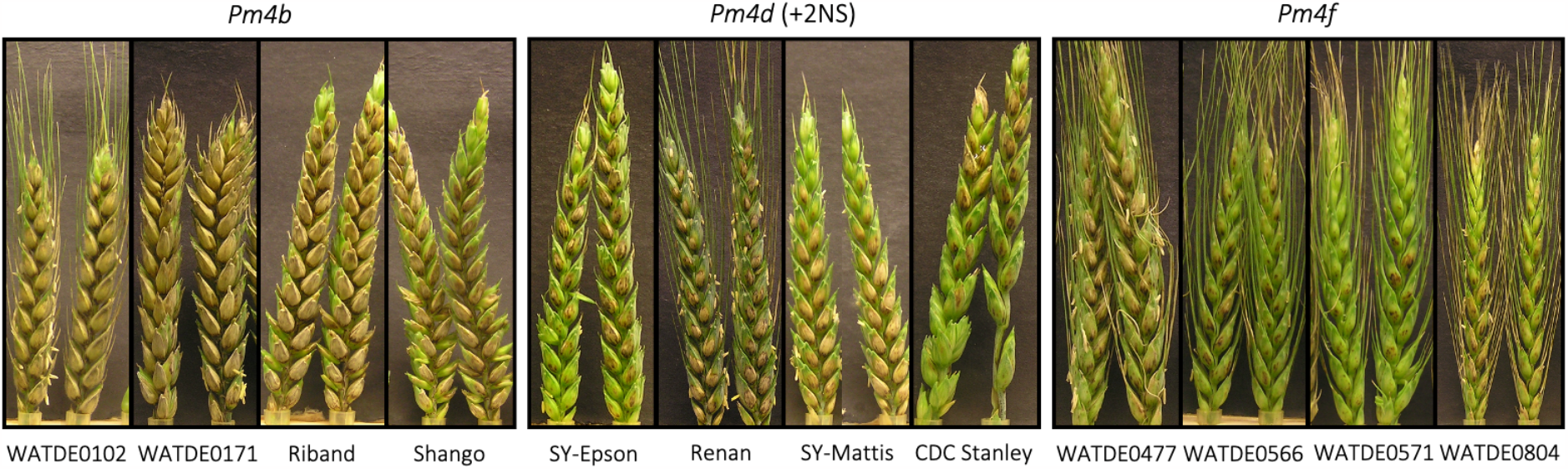
Identifying if Pm4 functions in the spike against Bangladeshi wheat blast isolates. Wheat blast detached spike assays for *Pm4b, Pm4d* and *Pm4f* alleles inoculated with Bangladeshi isolate BTJ4P-1 at 22 °C.

The NO6047+*AVR8* Watkins phenotype data was analysed using nucleotide-binding domain leucine rich repeat (NLR)-enriched *k*-mer based association genetics (henceforth referred to as WatRenSeq; (Arora et al., 2023)) with SY-Mattis as the reference genome. This produced a clear association peak on chromosome arm 2AL, spanning 788.8 to 794.1 Mbp. The Py 15.1.018 leaf disease scores were also analysed using SY-Mattis as the reference and produced an identical association peak (**Figure 1a**). Ten Watkins accessions highly resistant to Py 15.1.018 produced associations within the same interval on chromosome arm 2AL using SY-Mattis as the reference (**Figure 1a**; **Supplementary Figs. 6 and 7; Supplementary Table 4**). These data mapped the resistance to *AVR*-*Rmg8* to a 5.3 Mbp chromosome 2A interval on SY-Mattis.

### Interrogating the 5.3 Mbp chromosome 2A interval

Haplotype analysis was run across the genomic sequence of the *AVR*-*Rmg8* resistance interval in the full Watkins collection (827 accessions) and a selection of modern wheat varieties (218 cultivars) using SY-Mattis as the reference genome (Ahmed et al., 2023; Cheng et al., under review). A cluster heat map with 50 kb window size was generated to identify regions identical or near-identical to SY-Mattis which revealed that an additional 20 accessions of the Watkins collection carry sections of the 5.3 Mbp interval. Within the 5.3 Mbp interval, two 1 Mbp blocks of similarity among 63 accessions were observed, ‘Region 1’ (788,550,000 to 789,550,000 bp) and ‘Region 2’ (793,250,000 to 794,250,000 bp) (**Figure 1c**). These 20 accessions, along with an additional ten lines that lack the *AVR*-*Rmg8* resistance interval were phenotyped with isolates NO6047+*AVR8* and Py 15.1.018 (**Supplementary Table 5**). A cluster heat map containing the 20 additional Watkins lines is shown in **Supplementary Fig. 8**. Interestingly, the wild tetraploid wheat accession 33255 was highly similar to all of ‘Region 1’ and to 600 kb of ‘Region 2’ indicating that the interval may have originated from a hybridisation between hexaploid wheat with a wheat wild relative similar to *T. turgidum* (**Figure 1c**; **Supplementary Fig. 8**) (Walkowiak et al. 2020). Five Watkins lines (WATDE0102, WATDE0171, WATDE0310, WATDE0566 and WATDE0804) contained only ‘Region 1’ but showed resistance, suggesting that the *AVR*-*Rmg8* resistance was contained within this interval (**Figure 1c**). In addition, 22 modern European wheat cultivars also possessed only the ‘Region 1’ SY-Mattis haplotype (**Supplementary Fig. 8**). These 22 cultivars were all resistant to both NO6047+*AVR8* and Py 15.1.018 isolates (**Supplementary Table 6**) confirming that the *AVR*-*Rmg8* resistance is within the 1 Mbp interval termed Region 1. Among the Watkins lines containing ‘Region 1’, two lines (WATDE0056 and WATDE0720) both lacked the proximal 400 kb of ‘Region 1’ and both were susceptible to NO6047+*AVR8* and Py 15.1.018 isolates indicating that the *AVR*-*Rmg8* resistance lies within this 400 kb region (788550000 - 788950000 bp).

The 400 kb *AVR*-*Rmg8* resistance interval within ‘Region 1’ contains ten annotated genes (**Figure 1d**; **Supplementary Table 7**), only five of which were expressed in RNAseq data from whole aerial organs at the three-leaf stage (**Supplementary Table 8**). Significantly, four lines (WATDE0048, WATDE0527, WATDE0568 and WATDE0592) classified as containing ‘Region 1’ were susceptible to both isolates. We therefore compared the sequences of the five expressed genes between the four susceptible lines and resistant lines using the whole genome Watkins sequencing data and SY-Mattis as the reference (Cheng et al., under review). Two of the genes (TraesSYM2A03G00828410, TraesSYM2A03G00828450) were monomorphic among susceptible and resistant lines and two (TraesSYM2A03G00828400, TraseSYM2A03G00828460) had polymorphisms which did not associate with the resistance phenotype (**Supplementary Table 9**). In the remaining gene (TraesSYM2A03G00828360), two of the susceptible accessions (WATDE0568 and WATDE0592) contained a T/A SNP converting amino acid 446 from tryptophan to a stop codon (W446*) (**Supplementary Table 10**), while the other two susceptible accessions (WATDE0048 and WATDE0527) possessed an identical G/A SNP converting amino acid 50 from alanine to glutamic acid (A50E). The sequences of the five non-expressed genes in the interval were also examined and no polymorphisms that segregated with resistance were identified. Thus, the combined haplotype and allelic diversity analyses identified TraesSYM2A03G00828360 as a strong candidate gene for recognising and conferring resistance to isolates of *MoT* carrying *AVR-Rmg8*.

The RNAseq data revealed that TraesSYM2A03G00828360 is alternatively spliced resulting in two potential transcripts. The intron/exon structure for the first five exons was the same in both transcripts, while the last exons were distinct. Transcript 1 produced a protein of 560 amino acids, while in Transcript 2 the fifth intron extended an additional 1082 bp (encapsulating the sixth exon from transcript 1) and produced a 747 amino acids protein. BLAST analysis of Transcript 1 revealed it to be identical to the previously reported chimeric protein of a serine/threonine kinase and multiple C2 domains and transmembrane regions that functions as the wheat powdery mildew (*Blumeria graminis* f. sp. *tritici* (*Bgt*)) race specific resistance gene *Pm4* (Sánchez-Martín et al. 2021). This study also established that *Pm4* has alternate splicing, producing ‘isoforms’ Pm4b-V1 (560 amino acids) and Pm4b-V2 (747 amino acids), corresponding to TraesSYM2A03G00828360 Transcript 1 and Transcript 2, respectively. Both isoforms are required to confer resistance to wheat mildew (Sánchez-Martín et al. 2021). This suggests that the wheat blast *AVR*-*Rmg8* resistance is encoded by *Pm4*.

To confirm recognition of *AVR-Rmg8* by *Pm4*, we used the germplasm resources previously developed to characterize its role in resistance to powdery mildew. This included near-isogenic lines for two functionally distinct *Pm4* alleles (*Pm4a* and *Pm4b*) in the susceptible wheat cultivar Federation (*Fed-Pm4a, Fed-Pm4b*), *Pm4b* EMS-induced mutants in *Fed-Pm4b*, and transgenic lines of susceptible cultivar Bobwhite S26 overexpressing *Pm4b* (**Supplementary Table 11**) (Sánchez-Martín et al. 2021). In order to relate differences in response specifically to the presence or absence of the *eI* allele of *AVR-Rmg8*, we inoculated this germplasm with isogenic transformants of *MoT* isolate Br48 differing in the presence of only *AVR-Rmg8*. Isolate Br48ΔeI has been disrupted to remove *AVR-Rmg8 eI* while this gene has been replaced in isolate Br48ΔeI+eI (Horo et al. 2020). Federation was susceptible to both Br48ΔeI and Br48ΔeI+eI, while *Fed-Pm4b* and *Fed-Pm4a* (carrying different *Pm4* alleles) were both resistant in seedling leaves to Br48ΔeI+eI (**Fig. 1e**; **Supplementary Fig. 9**). All three lines were susceptible in spikes inoculated and incubated at 22 °C to both Br48ΔeI and Br48ΔeI+eI indicating that these alleles (*Pm4a* and *Pm4b*) only function in seedling resistance (**Fig. 1e**; **Supplementary Fig. 9**). Loss of wheat blast resistance in adult plants of many wheat varieties has been observed previously but the reasons for tissue- or stage-specific resistance is unknown (Ceresini et al. 2018).

All eight loss-of-function EMS-induced mutants of *Fed-Pm4b* were susceptible to both *MoT* isolates in seedling assays (**Fig. 1e**; **Supplementary Figs. 9 and 10**). Mutations were present in exon 6 and 7 specific to *Pm4b_V1* and *Pm4b_V2* respectively indicating that both transcripts are required for resistance to *MoT* as was found to be the case for *Bgt* (Sánchez-Martín et al. 2021). While Bobwhite S26 and lines S#3 and S#52 segregating from the T_1_ plants but lacking the transgene were susceptible to Br48ΔeI and Br48ΔeI+eI in seedling assays, both *Pm4b* over-expressing lines (Nr#3 and Nr#52) carrying the full length cDNAs of *Pm4b_V1* and *Pm4b_V2* were resistant to Br48ΔeI+eI. Surprisingly, Nr#52 was resistant to Br48ΔeI+eI in spikes while Nr#3 was susceptible (**Fig. 1e**). Expression of *Pm4b_V1* and *Pm4b_V2* was assessed in spike tissues of Nr#3 and Nr#52. Expression of *Pm4b_V1* was significantly higher in Nr#52 compared to Nr#3 (*p* value < 0.001) indicating that overexpression is sufficient to confer recognition and resistance in spike tissues to *MoT* isolates carrying *AVR-Rmg8* (**Supplementary Fig. 11**). The requirement for multiple copies or high expression of genes to provide full resistance has been reported recently suggesting that increased copy number may provide a route to increase disease resistance (Brabham et al. 2023), suggesting that increased copy number, or expression levels, may provide a route to increase disease resistance.

### Allelic variation

We designed PCR-based assays to detect *Pm4* and used these to investigate its prevalence in landraces and modern adapted varieties (primers detailed in **Supplementary Table 12**). We found *Pm4* to be uncommon among the landraces within the Watkins collection being present in only 29 of 827 (3.5%) accessions. The proportion of *Pm4*-containing varieties was higher (15.5%; 67 out of 432) in the ‘Gediflux’ collection of highly successful European varieties from the period 1945-2000 (Wingen et al. 2014) (**Supplementary Table 13**). This probably reflects the selection of *Pm4* by breeders in Europe to control mildew while this disease is of lesser importance in many other parts of the globe.

An allelic series of *Pm4*, each recognising different isolates of *B. graminis* f. sp. *tritici* has been reported and many of these originate from wild relatives of *T. aestivum* (Sánchez-Martín et al. 2021). *Pm4a* and *Pm4b* were introduced from tetraploid wheats (Briggle 1966; McIntosh and Bennett 1979), while *Pm4d* is believed to have been introgressed from *T. monococcum* (Schmolke et al. 2012). The origin of *Pm4f, Pm4g* and *Pm4h* is unknown. The two alleles of *Pm4* identified within this study (A50E and W446*) had sequences most similar to *Pm4f* but have not been reported previously. We designated these as *Pm4i* and *Pm4j*, respectively (**Table 2b**). *Pm4b* was the most common *Pm4* allele among modern wheat varieties (69%), but it was rare among the Watkins collection (11%). In contrast, *Pm4f* was absent in the modern varieties but was the most common allele within the Watkins collection (75%) (**Supplementary Table 14**). *Pm4d* was absent within the Watkins collection but was the second most common allele within modern varieties. This allele was only found in combination with the 2NS segment (33 Mbp (Gao et al. 2021)) on the short arm of chromosome 2A introgressed from *Aegilops ventricosa* into the wheat cultivar VPM1 (Vedel et al. 1981) (**Supplementary Table 14**). The 2NS segment on the short arm of chromosome 2A carries the rust resistance genes *Sr38, Yr17* and *Lr37* along with resistance to isolates of *MoT* (Cruz et al. 2016). It has been proposed that *Pm4* was introduced into the long arm of chromosome 2A from the *Triticum persicum* parent of VPM1 (Bariana and McIntosh 1993). The absence of *Pm4d* in Watkins accessions and its presence alongside the 2NS segment from *Ae. ventricosa* in modern varieties supports this view and indicates that the *Pm4d* allele may have been introduced only once into *T. aestivum* through VPM1 at the same time as 2NS but at the opposite end of the 2A chromosome and from a different wheat relative. This represents a second example of the serendipitous introduction of resistance into wheat from VPM1 as this line was originally developed to introduce the *Pch1* eyespot resistance gene on chromosome 7D^v^ of *Ae. ventricosa* into wheat (Doussinault et al. 1983) and the presence of the 2NS on the end of the short arm of chromosome 2A was not recognised.

The efficacy of seedling and spike resistance against *AVR-Rmg8* (isolate Py 15.1.018) was compared at 22 °C and 26 °C across wheat accessions carrying different *Pm4* alleles as it has been reported that resistance to *MoT* is often temperature sensitive (Anh et al. 2018). Carriers of *Pm4b, Pm4d, Pm4f* were all resistant at the seedling stage at both temperatures (**Fig 2a**; **Supplementary Fig. 12**). These three *Pm4* alleles, however, differed in efficacy in spikes. Carriers of *Pm4d* and *Pm4f* were resistant at 22 °C while *Pm4b* carriers were susceptible confirming the ineffectiveness of the *Pm4b* allele observed in the *Fed-Pm4b* NIL (**Figs. 1e and 2a**). The level of expression of the *V1* and *V2* transcripts of *Pm4b* and *Pm4f* in spikes were not significantly different among the wheat varieties examined (*p* value ≥ 0.314 and *p* value ≥ 0.750 or *V1* and *V2* transcripts respectively) indicating that differences in resistance more probably reflects differences in interaction between host and pathogen components than differences in expression of *Pm4* (**Supplementary Fig. 11**). Carriers of *Pm4d* expressed moderate resistance in the spikes at 26 °C while carriers of *Pm4b* and *Pm4f* were susceptible at this temperature. It should be noted that all the carriers of *Pm4d* also contained the 2NS segment that functions only in spike tissues (Cruz et al. 2016) and the resistance may reflect the presence of the two resistances in these varieties.

### The relative effectiveness of *Pm4* alleles against different alleles of *AVR-Rmg8*

Three alleles of *AVR-Rmg8* (*eI, eII, eII’*) were identified among *MoT* isolates collected in Brazil with *eII* being predominant (Wang et al 2018). The clonal lineage present in Bangladesh and Zambia, however, contains the eI allele (Horo et al. 2020). Isolates transformed to carry different alleles of *AVR-Rmg8* (*eI, eII and eII’*) differed in aggressiveness towards a wheat line (IL191) carrying *Rmg8*, with resistance being more pronounced against isolates carrying *eI* than those carrying *eII* or *eII’* (Horo et al. 2020).

The relative effectiveness of *Pm4* alleles against different alleles of *AVR-Rmg8* was examined by screening seedlings of wheat lines carrying different *Pm4* alleles for resistance to transformants lacking *AVR-Rmg8* or carrying *eI, eII* or *eII’* alleles (**Supplementary Table 15**) (Horo et al. 2020). The majority of *Pm4* alleles conferred resistance to all three *AVR-Rmg8* effector alleles (**Fig. 2c**). Alleles *Pm4g* (accession WW-470, (Pont et al. 2019)) and *Pm4j* did not confer resistance against any of the three *AVR-Rmg8* effector alleles. The lack of effectiveness of *Pm4j* was expected as this protein is truncated and *Pm4g* was previously reported to be a susceptible *Pm4* allele with respect to resistance to *Bgt* (Sánchez-Martín et al. 2021). Interestingly, the same study also reported *Pm4f* to be a *Bgt-*susceptible allele but it is effective against the three alleles of *AVR-Rmg8. Pm4a, Pm4b, Pm4d* and *Pm4h* (accession WW-474, (Pont et al. 2019)) are also highly effective against the three alleles of *AVR-Rmg8*. The two accessions carrying *Pm4i* (WATDE0048 and WATDE0527) showed greater resistance against Br48 carrying the *eI* or *eII* allele of *AVR-Rmg8* than against the same isolate carrying the *eII’* effector allele (**Fig 2c**; **Supplementary Table S15**). Differences in aggressiveness of isolates carrying different *AVR-Rmg8* alleles has also been noted previously (Horo et al. 2020). These two accessions, however, were highly susceptible to Py 15.1.018 (*eI*) and NO6047+*AVR8* (*eI* and *eII”‘*) (**Fig 2a, Supplementary Fig. 13**). We postulate that this may be due to the presence of additional effectors in these isolates that suppress *Pm4i* alleles in an equivalent manner to that reported for *PWT4* (Inoue et al. 2020).

The pandemic clonal lineage of MoT present in Bangladesh and Zambia contains the *eI* allele of *AVR-Rmg8* (Latorre et al. 2023) and so it was important to demonstrate whether *Pm4* alleles would function against this. Furthermore as the impacts of MoT infection are most dramatic for spike disease we screened spikes of wheat accessions carrying different *Pm4* alleles for resistance to the Bangladesh isolate BTJP4-1 (Latorre et al 2023). As anticipated from the studies above, accessions carrying *Pm4b* did not exhibit resistance to BTJP4-1 in spikes. In contrast, spikes of accessions carrying *Pm4f* were highly resistant to this isolate (**Fig. 3**). Spikes of accessions carrying *Pm4d* showed moderate resistance to BTJP4-1 but this probably reflects the presence of the 2NS segment in all these accessions. Assessment of CIMMYT’s international screening nurseries has revealed that the 2NS segment contributes almost all the wheat blast resistance present within both the Bread Wheat Screening Nurseries and the Semi-Arid Wheat Screning Nurseries (Juliana et al. 2020). These authors emphasized the urgent need to identify additional non-2NS sources of resistance. We believe that the *Pm4f* allele provides such a source.

## Discussion

No resistance gene against *MoT* has been cloned to date. In this study we used isolates carrying the *eI* allele of *AVR-Rmg8* to screen a genome-sequenced diversity panel of hexaploid wheat and clone a resistance gene recognising this effector. Surprisingly, this gene, encoding a serine-threonine kinase-MCTP protein, has previously been identified as *Pm4*, a race specific resistance gene to wheat powdery mildew (Sánchez-Martín et al. 2021). This is the second example of resistance to blast and mildew being conferred by the same gene. The NLR mildew resistance gene *MLA3* of barley also recognises the effector *PWL2* that acts as a host range determinant preventing *M. oryzae* from infecting weeping lovegrass (*Eragrostis curvula*) (Brabham et al. 2022). Furthermore, *Pm24*, an allele of the tandem kinase *Rwt4* that recognises *PWT4* of *M. oryzae*, (Arora et al. 2023) also confers resistance against powdery mildew of wheat (Lu et al 2020). The stem rust resistance gene *Sr62* from *Aegilops sharonensis* has been identified as an orthologue of *Pm24/Rwt4* (Yu et al. 2023). A small number of genes of different classes have been shown to confer resistance to multiple pathogens in wheat or its progenitors. These include the NLRs *Mla7* and *Mla8* (Bettgenhaeuser et al. 2021), and the receptor like kinase *TuRLK1* (Zou et al. 2022) both of which confer resistance to wheat powdery mildew and wheat yellow rust.

The predominant *Pm4* allele (*Pm4f*) identified within the Watkins Collection functions against wheat blast in both seedling and spike tissues but was not found within the collection of elite European varieties so this study also highlights the potential value of landrace materials as sources of resistance. It also appears that *Pm4* may have originated in wild tetraploid species in which a number of alleles were identified. Extending analysis to additional tetraploid species may reveal new *Pm4* alleles with greater efficacies against the different alleles of *AVR-Rmg8* and so improve resistance against wheat blast in Bangladesh, Zambia and South America. *Pm4* was not found in a selection of 565 CIMMYT wheat breeding lines carrying the 2NS translocation originating from wheat line VPM1 (**Supplementary table S17**). As VPM1 was also the donor of *Pm4d* into wheat this suggests that *Pm4d* was lost, probably because mildew resistance is not a major breeding target for CIMMYT. These findings are supported by genome wide association studies of wheat blast resistance in CIMMYT’s international screening nurseries in Bangladesh and Bolivia where the only robust and potent resistance was associated with 2NS, with no resistance identified in the region of *Pm4* (Juliana et al 2020; He et al 2021). This emphsises the importance of maintaining selection for as wide a range of diseases as practical because of the potential for unexpected benefits of resistances effective against multiple pathogens. Conventionally, the search for new resistances centres upon examining accessions originating from regions where the disease is believed to have originated on the assumption that co-evolution of host and pathogen will have led to selection for resistance. This approach is not possible with wheat blast as this disease first appeared in 1985. The finding that, at least some, resistances effective against powdery mildew in wheat also confer resistance to wheat blast suggests that the search for additional resistances might benefit from focus on non-obvious regions with cool, damp environments where wheat mildew is most problematic.

The urgent need to identify additional wheat blast resistance sources to complement and protect against loss of efficacy of the 2NS resistance is widely recognised (Phuke et al. 2022; Juliana et al. 2020). We believe that *Pm4* offers the first such resistance and provides an important entry point to lead to identification of additional resistances. Additional research is required to determine whether other mildew or rust resistances also confer resistance against wheat blast and to establish the basis of similarity in resistances against *P. oryzae, B. graminis* and *Puccinia* species through comparison of effectors, resistance genes and their interactions.

## Methods

### Phenotyping of the Watkins panel and additional materials for validation with wheat blast isolates

The *MoT* isolate Py 15.1.018 and the transformed isolates Br48ΔeI and Br48ΔeI+eI (Horo et al. 2020) were grown on complete medium agar. The *MoT* isolate NO6047 and the transformed isolate NO6047+*Avr*8 were grown on oatmeal media. Oatmeal media was prepared by adding powdered oats (40 g) to 500 ml of dH_2_O and placed in a water bath at 65°C for one hour, and then filtered through two layers of muslin. As much liquid was extracted from the oats as possible before dividing equally between two 1 L Schott bottles. Agar (10 g) (Sigma Aldrich) and 2.5 g sucrose were added to each bottle and the volume was made up to 500 ml before autoclaving. Fungal inoculum was prepared as described by Goddard et al. (2020). A conidial suspension of 0.2 – 0.4 × 10^6^ conidia per millilitre was used for all inoculations. Seedling and spike assays were carried out as described by Goddard et al. (2020) and scored for disease symptoms using a 0–6 scale as described by Arora et al. (2023). Five and three biological replicates were used for seedling and spike assays respectively.

### *k*-mer-based association mapping

Association mapping was performed using similar methods to Arora et al. (2023) using the reference genome SY-Mattis (Walkowiak et al. 2020).

### Haplotype analysis

Haplotype analysis was used to refine the chromosome 2A WatRenSeq candidate region. In brief, a visualization cluster heat map using the “*variations*” database described in https://github.com/Uauy-Lab/IBSpy was used to identify candidates sharing similarity with the SY-Mattis genome reference within the chromosome 2A interval. The complete *variations* data are available from Zenodo under the DOI zenodo.org/record/8355991. Samples with similar *variations* profiles in the target region were selected to run short reads alignments against the SY-Mattis reference (Walkowiak et al. 2020). Alignments for the ten Watkins accessions with the chromosome 2A WatRenSeq association, in addition to ten adapted varieties and the 20 Watkins accessions outside of the core panel identified as having the chromosome 2A interval were generated. The alignments were produced using bowtie2 (v.2.4.1) (**Supplementary Tables 9 and 10**) (Langmead and Salzberg 2012) (Cheng et al., under review). Alignments in SAM format were processed using samtools (v.1.7) (Li et al. 2009) and transformed to BAM format removing duplicates and filtering for mapping quality (MAPQ) >30. BAM files were visualized with IGV (v.2.8.0; (Robinson et al. 2011)) to detect candidate SNPs within the 400kb *Rmg8* interval.

### Genome annotations and RNAseq analysis

Pre-publication access was granted to the SY-Mattis RNA-Seq data and gene annotations (version 03G) generated by the 10+ Wheat Genome project. Note that the current public annotation on EnsemblPlants is version 01G.

Seedlings were grown in a Controlled Environment Room (CER) (Conviron BDW80; Conviron, Winnipeg, Canada) set at 16 h day/8 h night photoperiod, temperatures of 20/16 °C, respectively, and 60% relative humidity. Plants were sampled at the 3-leaf stage (Zadocks GS13), harvesting whole aerial organs separately 4 hours after dawn (09:00). Each biological replicate consisted of a single plant. RNA sequence data from two biological replicates were mapped to the SY-Mattis reference genome using the Linux programme HISAT2 (v.2.1.0) (Walkowiak et al. 2020; Kim et al. 2019). BAM files were viewed using IGV (v.2.8.0) (Robinson et al. 2011).

### KASP marker design to detect *Pm4* wheat cultivars, Watkins collection and Gediflux collection

The wheat reference genome Chinese Spring (IWGSC et al. 2018) lacks *Pm4* (Sánchez-Martín et al. 2021). A region in exon 7 differentiating the functional *Pm4* from its closest (non-functional) homeologue in Chinese Spring, within gene TraesCS2A01G557900, was used to design KASP (LGC genomics) markers to distinguish *Pm4/pm4* carriers in wheat (**Supplementary Table 12**) (Sánchez-Martín et al. 2021). The KASP markers were validated on genome-sequenced cultivars and a subset of Watkins and adapted lines (Walkowiak et al. 2020). Subsequently, KASP marker analysis was performed on the Gediflux Collection (497) to understand the distribution of *Pm4* within northern European wheat (**Supplementary Table 13**). 67 accessions were identified as containing a *Pm4* allele. All reactions were run using the following touchdown PCR programme using an Eppendorf vapo.protect Mastercycler pro 384 (Eppendorf AG, 22331, Hamburg, Germany): 94°C for 15 minutes, 94°C for 20 seconds followed by 65°C for 1 minute (repeated ten times, decreasing by 0.8°C each cycle to 57°C), 94°C for 20 seconds followed by 57°C for 1 minute (repeated 30 times) and held at 16°C. An additional five to ten cycles were sometimes required for full separation of the signals from the different genotypes. Plates were read using a PHERAstar microplate reader (BMG LABTECH, Allmendgrün 8, 77799, Ortenberg) and analysed using the genotyping data analysis software KlusterCaller.

### Data availability

The RenSeq 150-bp paired-end Illumina sequences (raw data) for the 300 Watkins and 20 non-Watkins accessions are available from NCBI study number PRJNA760793 (reviewer link: https://dataview.ncbi.nlm.nih.gov/object/PRJNA760793?reviewer=jjvh7733t34pu9ls4sdviut9nd). The *k*-mer matrix and the CLC assemblies of the 300 Watkins and 21 non-Watkins accessions are available from Zenodo under the DOIs: 10.5281/zenodo.5557564, 10.5281/zenodo.5557685, 10.5281/zenodo.5557721, 10.5281/zenodo.5557827, 10.5281/zenodo.5557838 and 10.5281/zenodo.5655720. Sequencing data was obtained from Sequence Read Archive (SRA) accession SRX9897426 (Wild_tetraploid) and from the National Genomics Data Center (NGDC) Genome Sequence Archive (GSA) BioProject accession number PRJCA019636.

### Quantitative Real-Time PCR

Expression of *Pm4b_V1* and *Pm4b_V2* in the spike was quantified through reverse transcription, quantitative real-time PCR (RT-qPCR) (**Supplementary Fig. 11**). Mature spikes were harvested and individually shock frozen in liquid nitrogen and stored at -70°C. Three biological replicates were sampled per genotype. RNA was extracted using RNeasy^®^ Plant Mini Kit (74904, Qiagen, Hilden, Germany) according to the manufacturer’s protocol. Immediately after extraction, the samples were purified using DNA Turbo DNA-*free*™ kit (01134216, Invitrogen) according to the manufacturer’s protocol. Samples were quantified using a NanoDrop™ spectrophotometer (Thermo Scientific™) and stored at -70°C. First strand cDNA was synthesised using SuperScript^®^ III First-Strand Synthesis System for RT-PCR kit (18080-051, Life technologies, Carlsbad, California, USA). 1 μL each of 50 μM oligo(dT)_20_ and 50 ng/μL random hexamers were included in the first-stand cDNA synthesis. Subsequent steps were carried out according to the manufacturer’s protocol. Samples were quantified using a NanoDrop™ spectrophotometer and stored at -20°C. RT-qPCR was performed with 4 μL of 10-fold diluted cDNA in technical duplicates using SYBR^®^ Green JumpStart™ Taq ReadyMix™ (S4438, Sigma-Aldrich, St. Louis, Missouri, USA) as described by Chen et al. 2006. All reactions were run using a CFX96 Real-Time system C1000™ Thermal Cycler (BioRad, Hercules, California, USA). Thermocycling conditions were 95°C for 4 minutes, followed by 40 cycles of 94°C for 10 seconds, then 60°C for 10 seconds, then 72°C for 30 seconds. Amplification specificity was confirmed using the ‘melting curve’ capability. Reference genes ADP and ZFL were used, as described by Sánchez-Martín et al. (2021). Average amplification efficiencies for the ADP, ZFL, *Pm4b_V1* and *Pm4b_V2* primers (**Supplementary Table 12**) were determined using a serial dilution (10-fold dilution decreasing to a 100,000-fold dilution) of a pool of six cDNA samples, to produce a standard curve. Target specific amplication efficiencies were calculated using the Agilent BioCalculator (https://www.agilent.com/store/bioCalcs.jsp#) and are given in **Supplementary Table 16**. Data are presented as the expression ratio of the target gene to the reference gene as described by Chen et al. (2006). Statistical analysis was performed using Genstat version 22.1 (VSN International 2022). GLM analysis using ‘replicate’ and ‘line’, to compare relative expression of *V1* and *V2* in Nr52 and Nr3. A two sampled t-test was used to compare relative expression of *V1* and *V2* in *Pm4b* and *Pm4f* alleles at *p* < 0.05.

## Code availability

Scripts for the Watkins *k*-mer matrix generation, phylogenetic tree construction and *k*-mer based association mapping can be found in the repository https://github.com/arorasanu/watkins_renseq. Scripts for the haplotype analysis are in https://github.com/Uauy-Lab/IBSpy.

## Acknowledgements

The high-performance computing resources and services used in this work were supported by the Norwich Bioscience Institutes Partnership (NBIP) Computing infrastructure for Science (CiS) group. We are grateful to the John Innes Centre (JIC) Horticultural Services for plant. We acknowledge the kind gift of wild type and transformed fungal isolates from Diane Saunders and Cassandra Jensen (John Innes Centre, UK), and from Soichiro Asuke and Yukio Tosa (Kobe University, Japan). We thank M. Spannagl (Helmholtz-Center Munich), Anthony Hall (Earlham Institute) and C. Pozniak (University of Saskatchewan) for pre-publication access to SY-Mattis annotation and RNA-Seq data. We are also grateful to Richard Gorum for assistance with DNA extractions and Phil Robinson for photography. This research was financed by the Biotechnology and Biological Sciences Research Council (BBSRC) Designing Future Wheat Cross-Institute Strategic Programme to BBHW, CU and PN (BBS/E/J/000PR9780) and BBSRC Norwich Research Park iCASE Doctoral Training Grant with Limagrain UK as industrial partner (BB/M011216/1) for supporting TOH. To the Mexican Consejo Nacional de Ciencia y Tecnología (CONACYT; 2018-000009-01EXTF-00306) and the JIC International Scholarships for supporting JQ-C.

## Author contributions

This work was conceived by PN, TOH and BBHW. Watkins panel configuration and sequence acquisition (LW, SC, CF). Association mapping (TOH, SA, KG), candidate gene discovery and analysis (TOH). Wheat haplotype analysis (JQ-C, RR-G, CU). Phylogenetic analysis (DG). Phenotyping of diversity panels, mutants and transformants (AS, RG, TOH), KASP marker design and analysis (TOH, AS), NILs, mutants, and wheat transformed lines (JS-M, BK), Statistical analysis (AS), Genotyping of CIMMYT lines (SD). Mutant confirmation and segregation (AS, RG, PN). Drafted manuscript (PN, TOH). Edited manuscript (BBHW, CU, BK, JSM) and designed figures (TOH, DG, JQ-C).

## Competing interests

The authors declare no competing interests.

## Notes

### Competing Interest Statement

The authors have declared no competing interest.

