## Supplementary figures for "The wheat powdery mildew resistance gene *Pm4* also confers resistance to wheat blast"

### Figure S1

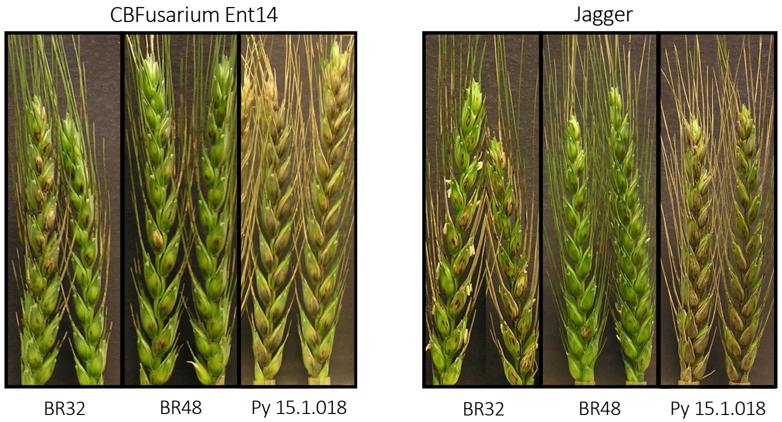


**Figure S1** - Wheat blast detached spike assays for *Ae. ventricosa* 2NS translocation containing cultivars CBFusarium Ent14 and Jagger. Spikes were inoculated with Brazilian isolates BR32, BR48 and Py 15.1.018 at 22 ºC. Images were taken at six and seven days post inoculation for Jagger and CB Fusarium Ent14, respectively.

### Figure S2

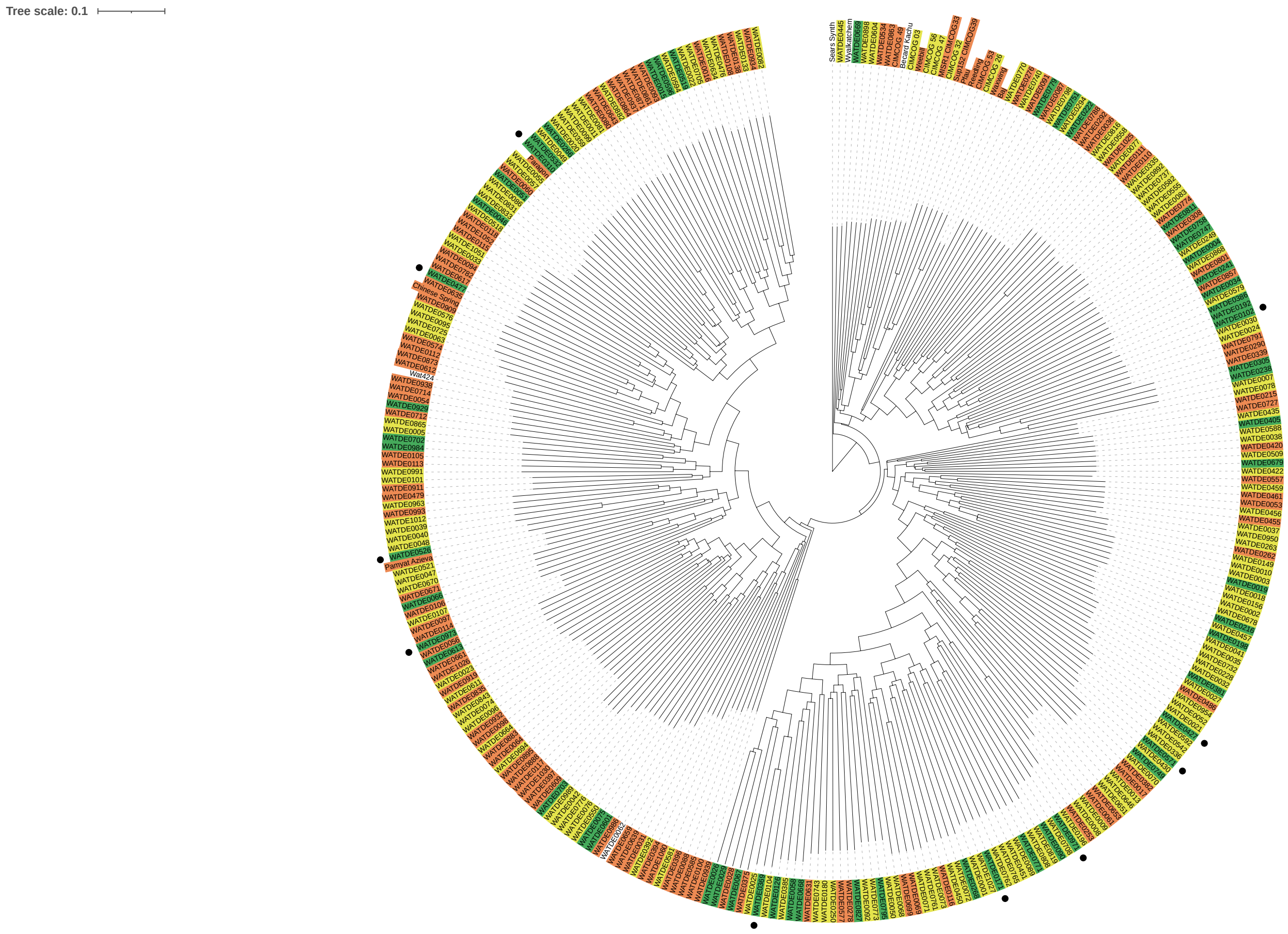


**Figure S2** - *k*-mer-based phylogeny of wheat landraces showing the phenotype of an accession after inoculation with NO6047+*AVR8.* Phenotype of an accession after inoculation is indicated by the colour used to highlight the label of that accession (green = resistant (scores less than or equal to 3 ), yellow = intermediate (scores more than 3, less than 5) and orange = susceptible (scores equal to greater than 5). Black circles indicate the presence of the chromosome 2A peak based on the WatRenSeq association plots.

### Figure S3

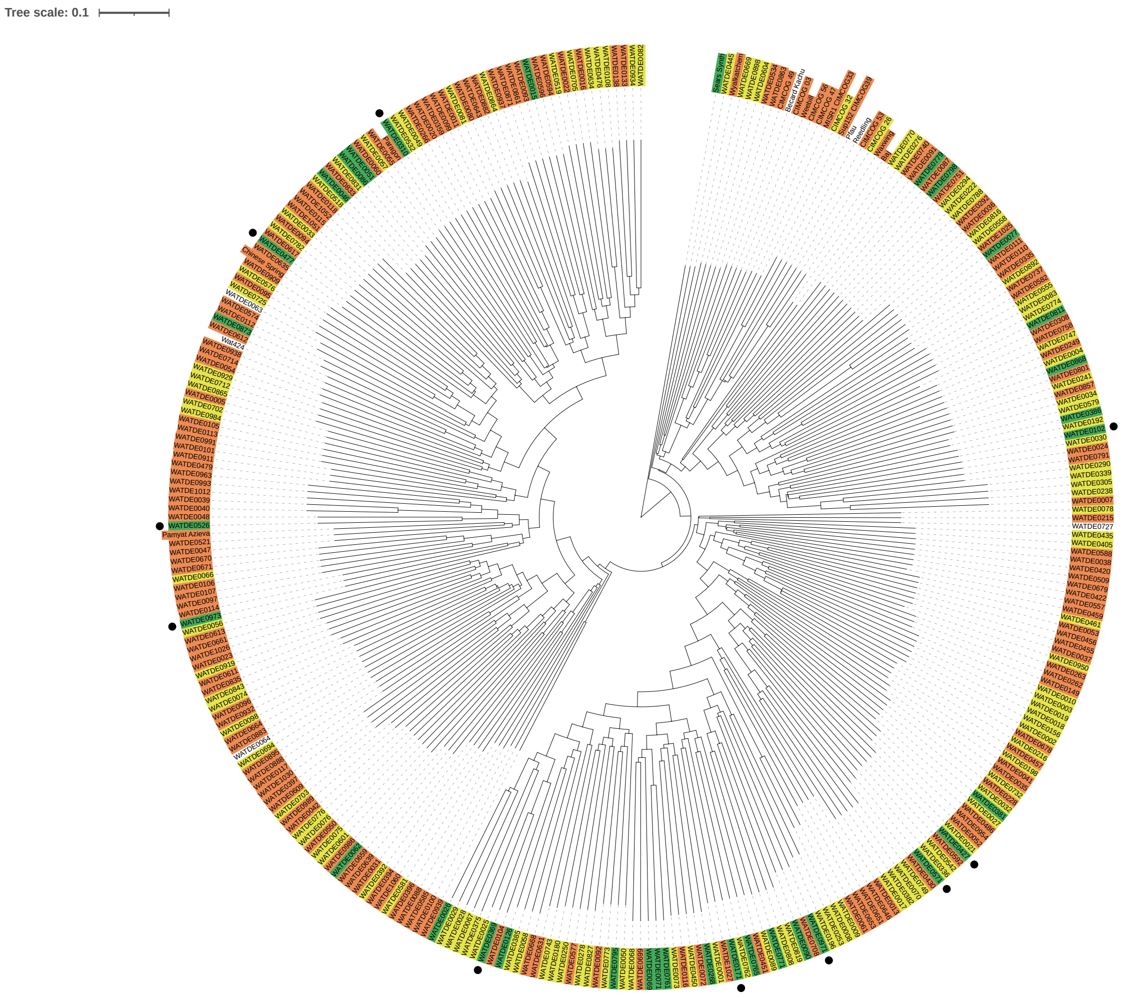


**Figure S3** - *k*-mer-based phylogeny of wheat landraces showing the phenotype of an accession after inoculation with Py 15.1.018. Phenotype of an accession after inoculation is indicated by the colour used to highlight the label of that accession (green = resistant (scores less than or equal to 3 ), yellow = intermediate (scores more than 3, less than 5) and orange = susceptible (scores equal to greater than 5). Black circles indicate the presence of the chromosome 2A peak based on the WatRenSeq association plots.

### Figure S4

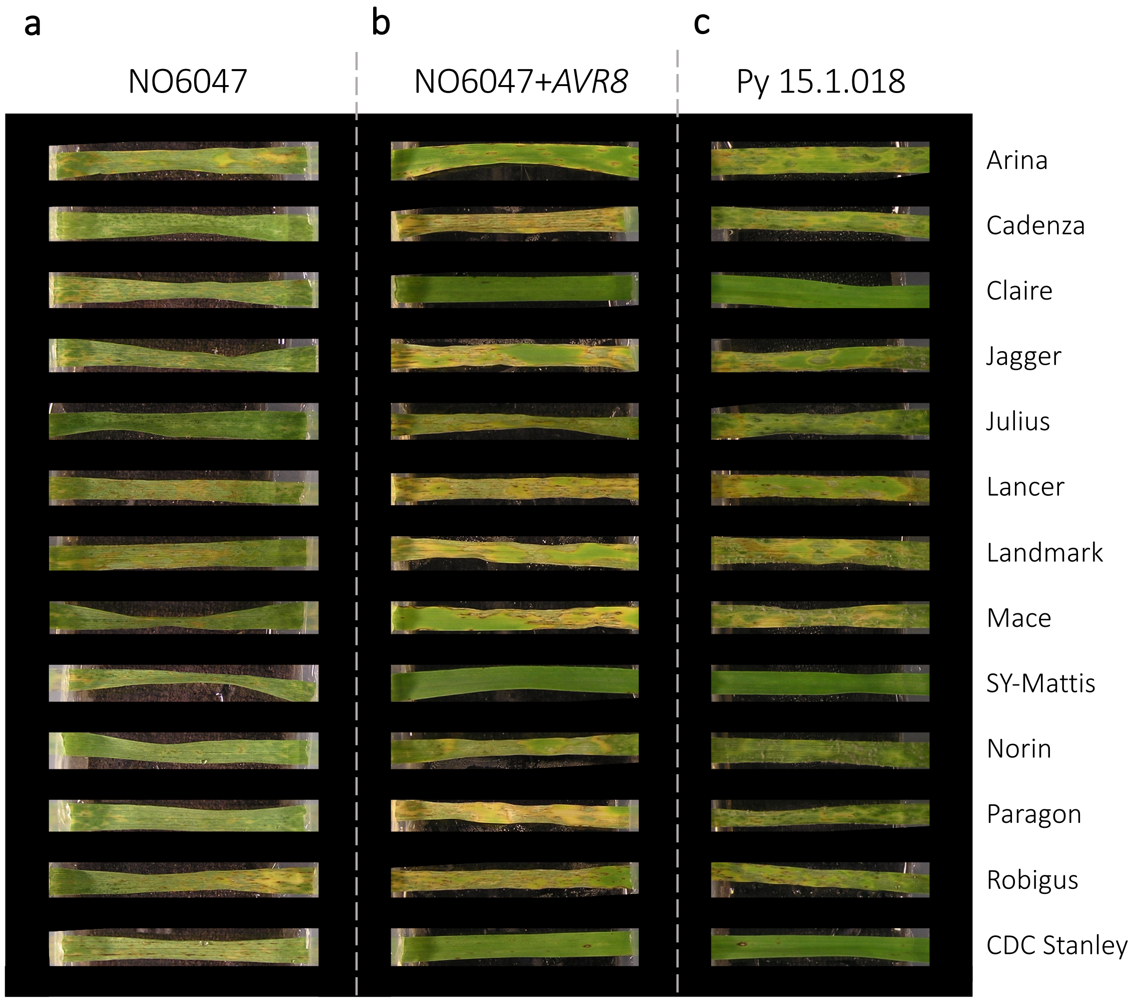


**Figure S4 -** Representative leaves of the 10 wheat cultivars with chromosome or scaffold-scale level assemblies used in this study. Leaves were inoculated at 22 ºC with **a**. NO6047, **b**. NO6047+*AVR8* and **c**. Py 15.1.018. Images were taken at five days post inoculation.

### Figure S5

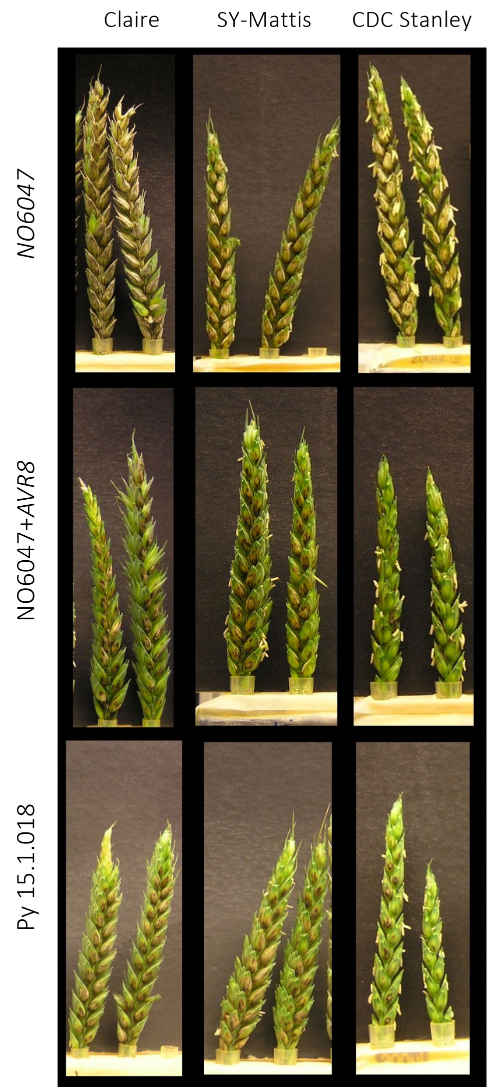


**Figure S5** – Comparison of detached spike phenotypes for Claire, SY-Mattis and CDC Stanley. Spikes were inoculated at 22 $^{o}$C with NO6047, NO6047+*AVR8* and Py 15.1.018. Images were taken at five to seven days post inoculation.

### Figure S6

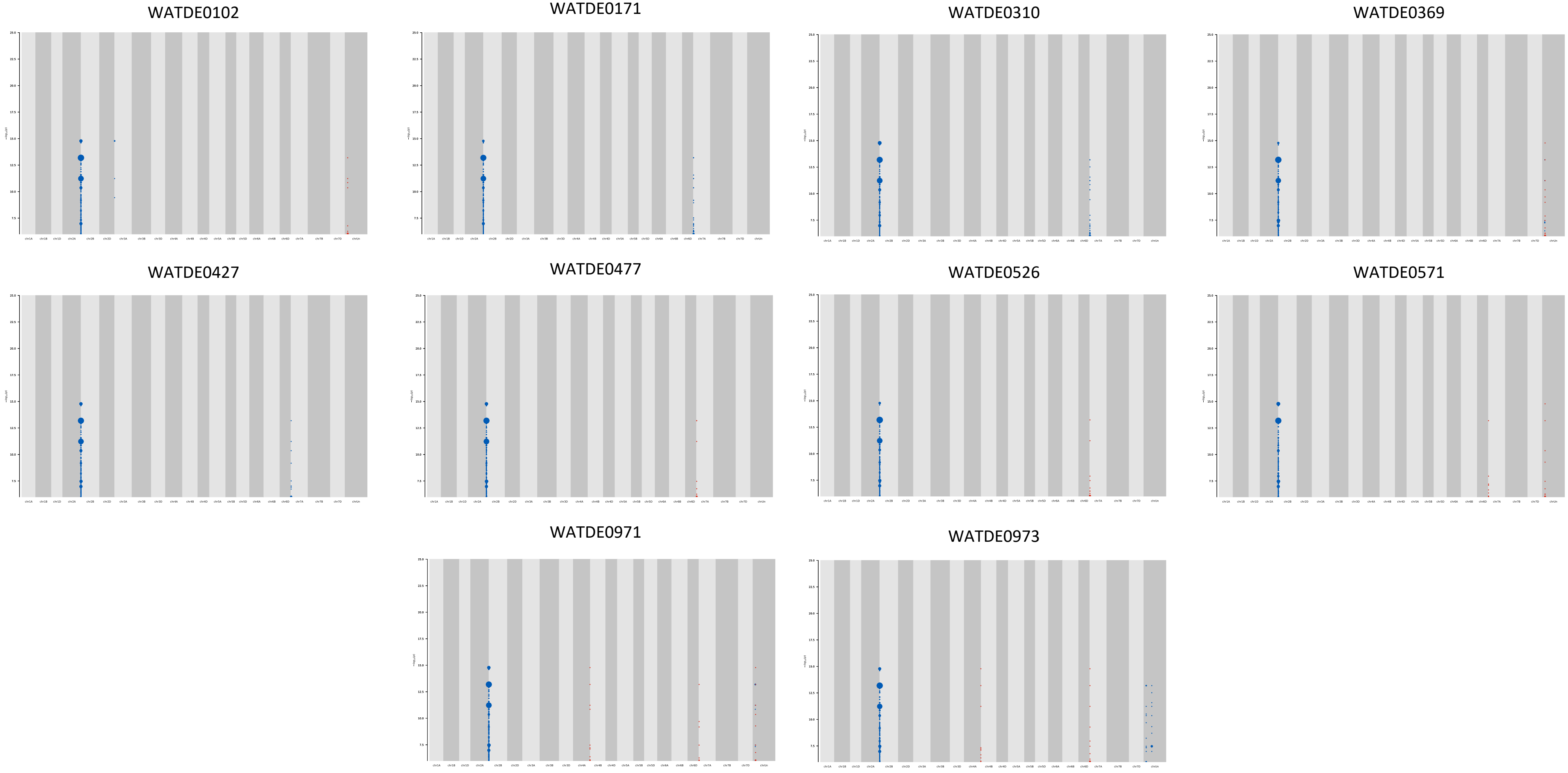


**Figure S6** - *k*-mers associated with resistance to NO6047+*AVR8* mapped to SY-Mattis for the *k*-mer compliment of the ten Watkins accessions that have the chromosome 2A association peak. Phenotype data was collected at five days post inoculation. Points on the *y* axis depict *k*-mers positively associated with resistance in blue, and negatively associated with resistance in red. Point size is proportional to the number of *k*-mers. The association score is defined as the –$\log_{10}$ of the *P* value obtained using the likelihood ratio test for nested models.

### Figure S7

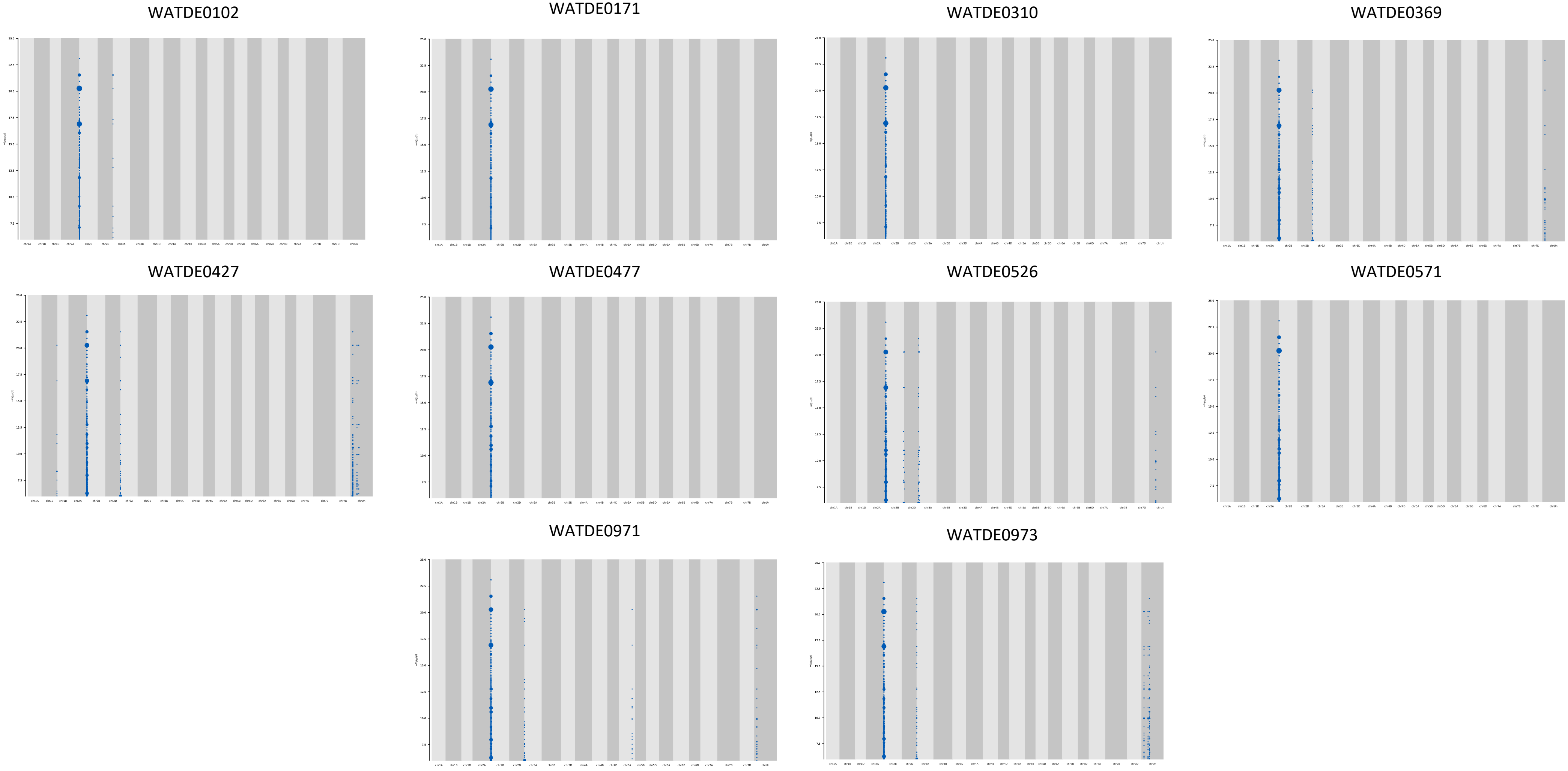


**Figure S7** - *k*-mers associated with resistance to Py 15.1.018 mapped to SY-Mattis for the *k*-mer compliment of the ten Watkins accessions that have the chromosome 2A association peak. Phenotype data was collected at five days post inoculation. Points on the *y* axis depict *k*-mers positively associated with resistance in blue, and negatively associated with resistance in red. Point size is proportional to the number of *k*-mers. The association score is defined as the $-\log_{10}$ of the *P* value obtained using the likelihood ratio test for nested models.

### Figure S8

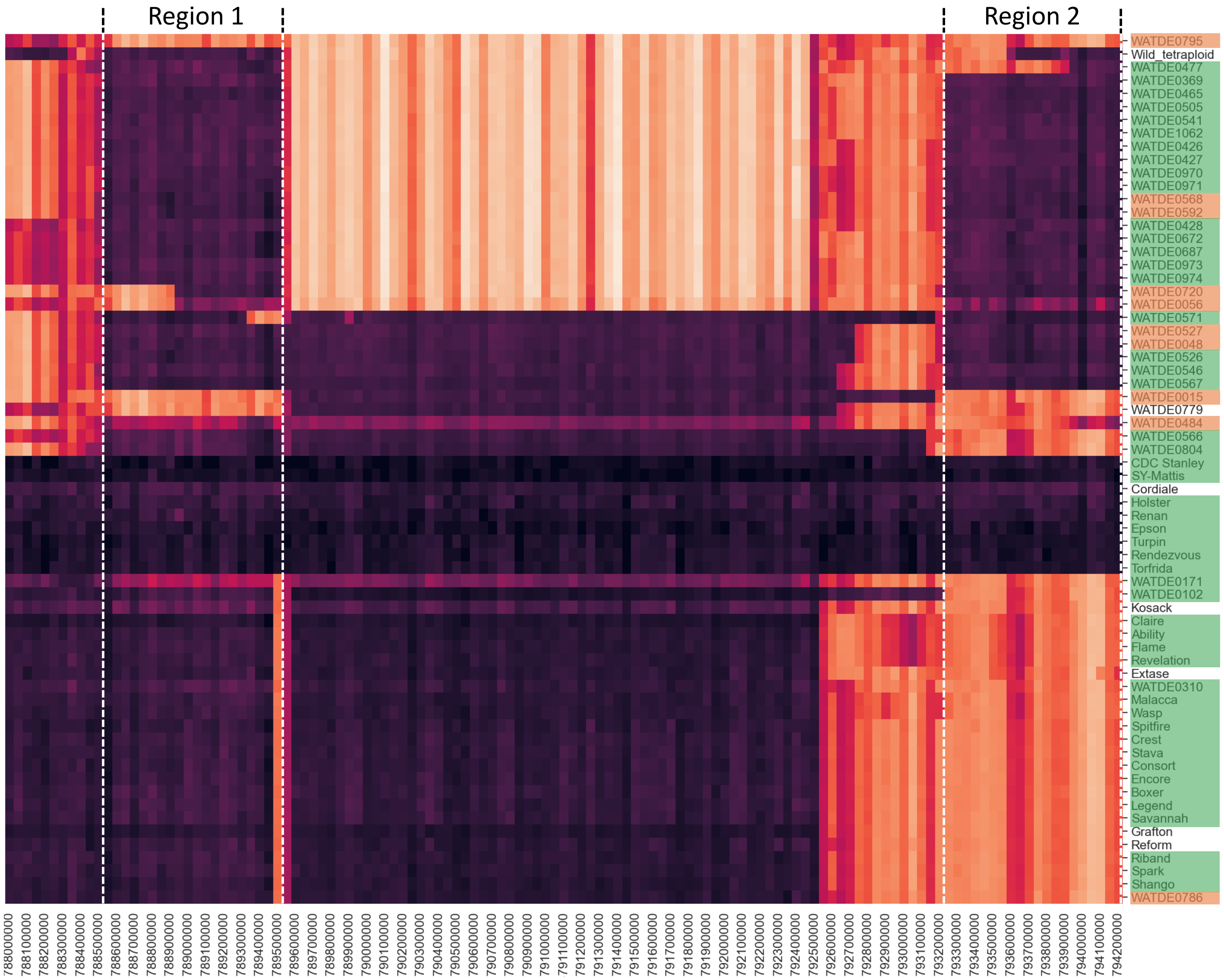


**Figure S8** - Haplotype cluster heatmap for the chromosome 2A interval using SY-Mattis as the reference. Phenotype of an accession after inoculation with Py 15.1.018 is indicated by the colour used to highlight the label of that accession, (green = resistant (scores less than or equal to 3), yellow = intermediate (scores more than 3, less than 5) and orange = susceptible (scores equal to greater than 5)). The darker the colour within a 50 kb window, the more sequence identity that sequence has to SY-Mattis. Two blocks of similarity were observed, 'Region 1' (788,550,000 to 789,550,000) and 'Region 2' (793,250,000 to 794,250,000). Note that the 'Region 1' haplotype block extends approximately 250 kb upstream of the 5.3 Mb chromosome 2A interval (788.8 to 794.1 Mbp). The variations data are available from Zenodo under the DOI zenodo.org/record/8377192.

### Figure S9

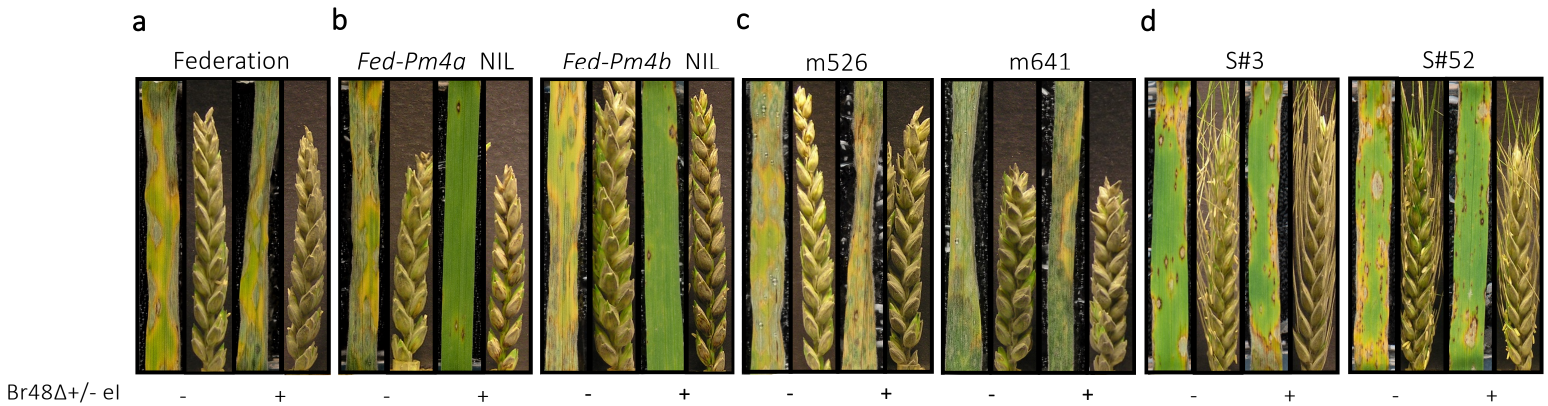


**Figure S9** - Wheat blast detached leaf and spike assays for **a**, the susceptible genetic background Federation(Fed). **b**, *Fed-Pm4a* and *Fed-Pm4b* NILs. **c**, *Pm4b* EMS-induced mutants of *Fed-Pm4b* NIL. Leaves and spikes were inoculated with Br48ΔeI and Br48ΔeI+eI, denoted by '-' and '+', respectively. **d**, Sister lines originating from the same $T_0$ as the transgenic *Pm4b* over-expressors in the Bobwhite S26 background that have lost the transgene through segregation, S#3 and S#52 are sister lines to Nr#3 and Nr#52, respectively (Fig. 1e).

### Figure S10

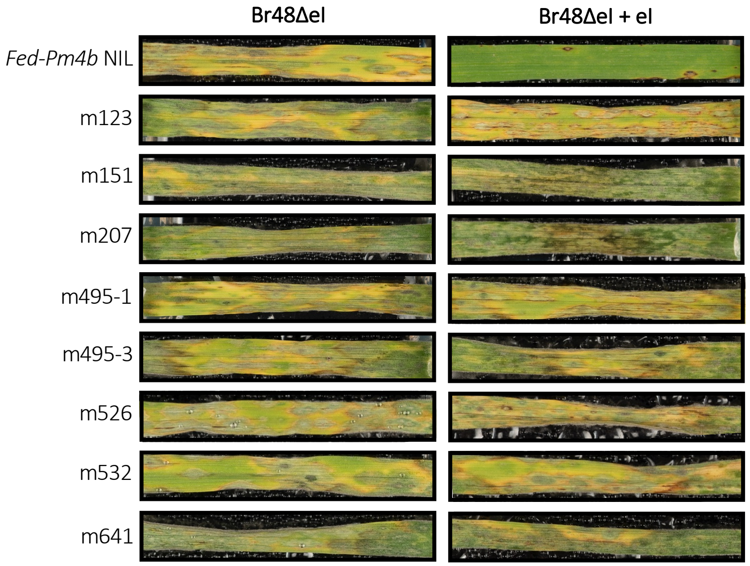


**Figure S10** - Wheat blast detached leaf assays for *Pm4b* EMS-induced mutants of *Fed-Pm4b* NIL. Leaves were inoculated with Br48ΔeI and Br48ΔeI+eI.

### Figure S11

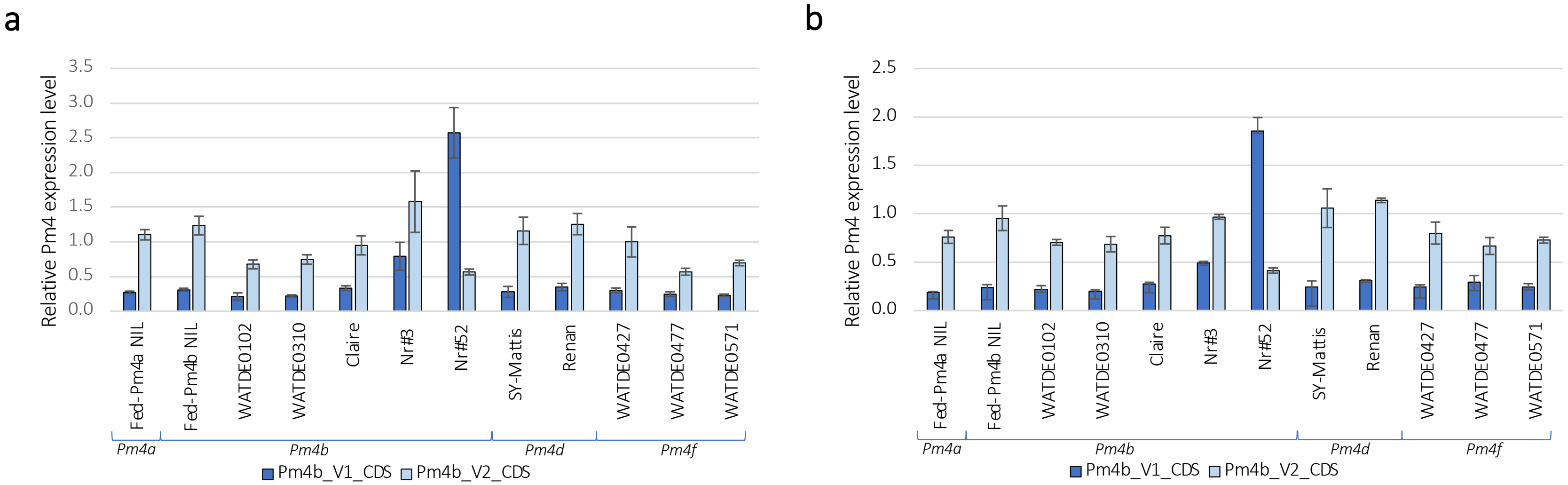


**Figure S11** – Expression levels of Pm4b_V1_CDS (dark blue) and Pm4b_V2_CDS (light blue) in the spike relative to reference genes (a) ADP and (b) ZFL. The data points comprise the mean of three biological replicates, each with two technical replicates . 'Nr#' = Pm4b transgenics (over expressors). The *Pm4* allele present in each accession is shown underneath the brackets. Error bars show the standard error. Statistical analysis was performed using Genstat version 22.1 (VSN International 2022).

### Figure S12

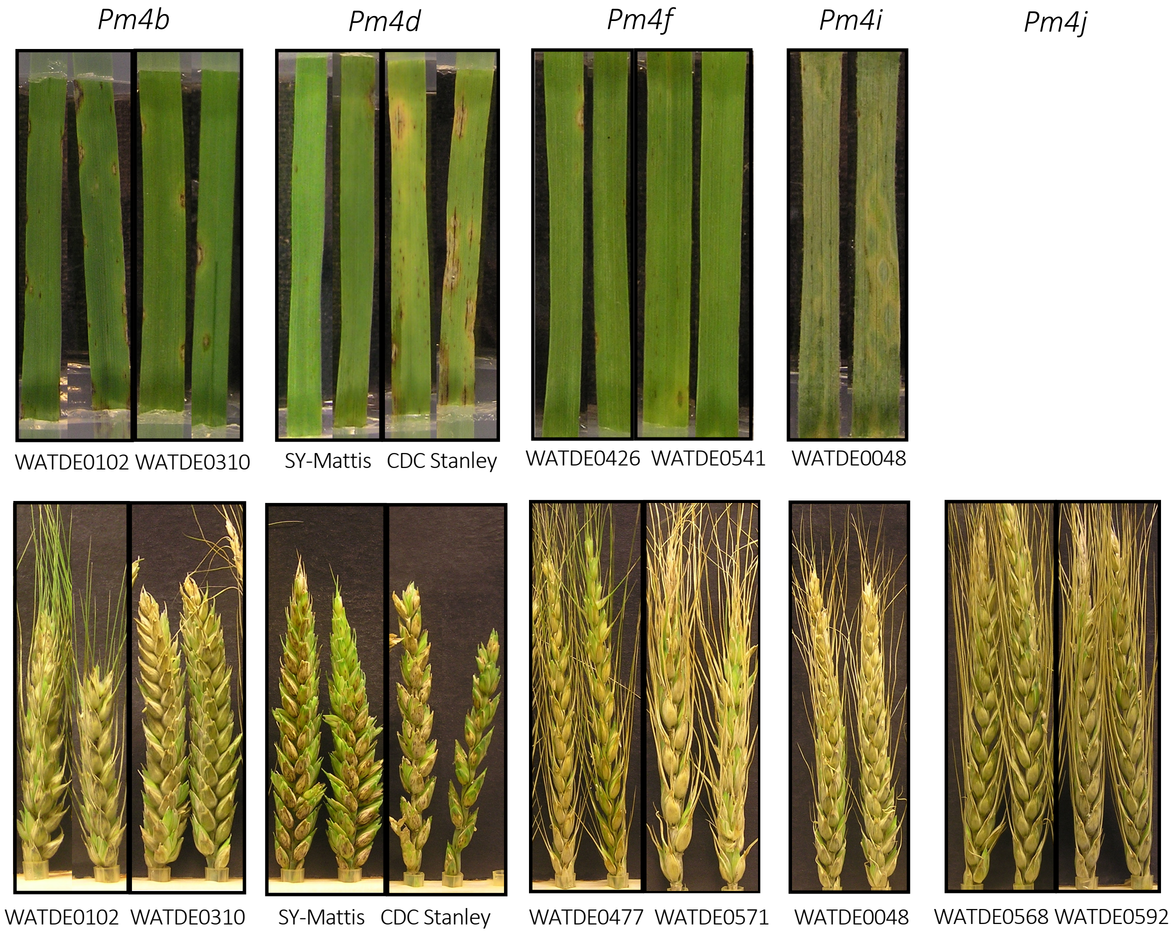


**Figure S12** - Wheat blast detached leaf and spike assays for *Pm4* alleles inoculated with *Py 15.1.018* at 26 ºC. Images were taken at five and four days post inoculation for leaves and spikes, respectively.

### Figure S13

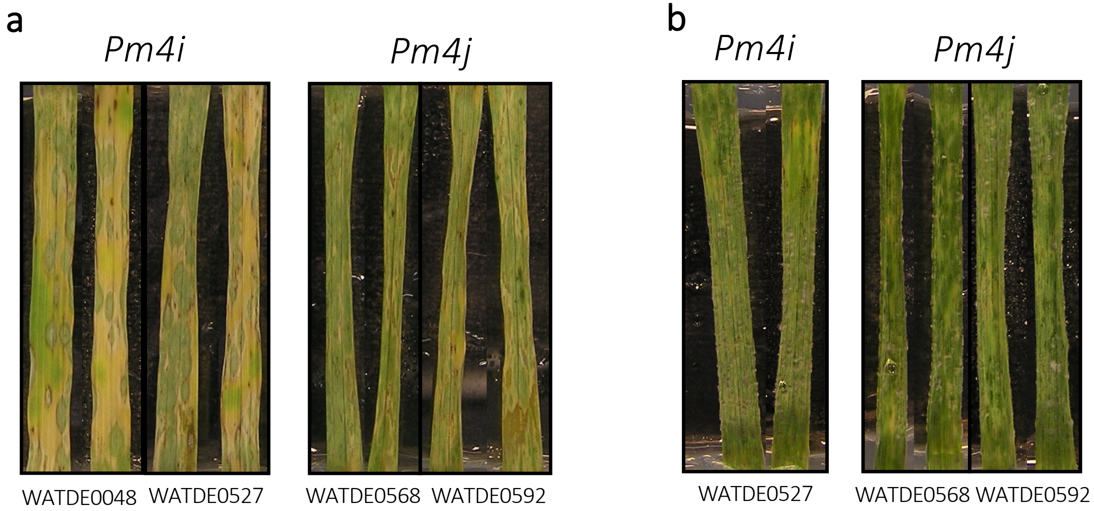


**Figure S13** - Wheat blast detached leaf and spike assays for *Pm4i* and *Pm4j* alleles inoculated with a. NO6047+*AVR8* **b**. Py 15.1.018 at 22ºC. Images were taken at five and six days post inoculation for NO6047+*AVR8* **and** Py 15.1.018 respectively.
