## Supplementary tables for "The wheat powdery mildew resistance gene *Pm4* also confers resistance to wheat blast"

**Table S1 -** List of blast resistance loci identified and mapped in wheat. Note only *Rwt3* and *Rwt4* have been cloned (Arora et al. 2023).

| **Gene** | **Source cultivar** | **Effective against** | ***Avr* genes** | **Chromosome** | **References** |
|---|---|---|---|---|---|
| *Rmg1 (Rwt4)* | Norin 4 | *Avena* isolate Br58 | *PWT4* | 1D (cloned) | (Takabayashi et al. 2002; Arora et al. 2023) |
| *Rmg2* | Thatcher | *Triticum* isolate Br48 | Not known | 7A | (Zhan et al. 2008) |
| *Rmg3* | Thatcher | *Triticum* isolate Br48 | Not known | 6D | (Zhan et al. 2008) |
| *Rmg4* | P168, Shin-chunaga, Norin 4, Norin 26, Norin 29 | *Digitaria* isolate | Not known | 4A | (Nga et al. 2009) |
| *Rmg5* | Red Egyptian and Salmon | *Digitaria* isolate | Not known | 6D | (Nga et al. 2009) |
| *Rmg6 (Rwt3)* | Chinese Spring, Shin-chunaga and Norin 4 | *Lolium* isolate TP2 | *PWT3* (or *A1*) | 1D (cloned) | (Vy et al. 2014; Arora et al. 2023) |
| *Rmg7* | St24, St17, St25 | *Triticum* isolate Br48 | *AVR-Rmg7* | 2A | (Tagle et al. 2015) |
| *Rmg8* | S-615 | *Triticum* isolate Br48 | *AVR-Rmg8* (=*AVR-Rmg7*) | 2B | (Anh et al. 2015; Anh et al. 2018) |
| *RmgGR119* | GR119 | *Triticum* isolate Br48 | Not known | - | (Wang et al. 2018) |
| 2NS | VPM1 | *Triticum* isolate Br48 | Not known | 2A | (Doussinault et al 1981) |

**Table S2** - The wheat cultivars with chromosome or scaffold-scale assemblies used in this study, their pedigree, origins and assembly type. Adapted from Supplementary Table 1 of Walkowiak et al. (2020). RQA: Reference Quality Pseudomolecule Assembly

| **Line** | **Pedigree** | **Growth Habit** | **Origin** | **Assembly Type** |
|---|---|---|---|---|
| Mace | WYALKATCHEM/STYLET//WYALKATCHEM | Spring | Australia | RQA |
| LongReach Lancer | VI184/Chara//Chara/3/Lang | Spring | Australia | RQA |
| CDC Stanley | CDC Teal//EE8/Kenyon35//AC Barrie | Spring | Canada | RQA |
| CDC Landmark | Unity/Waskada//Alsen/Superb | Spring | Canada | RQA |
| Julius | Asketis/Drifter | Winter | Germany | RQA |
| Norin 61 | Fukuoka Komugi 18/Shinchunaga | Facultative Spring | Japan | RQA |
| Arina*LrFor* | Arina*3/Forno | Winter | Switzerland | RQA |
| Jagger | KS-82-W-418/STEPHENS | Winter | USA | RQA |
| SY-Mattis | Apache/Intense | Winter | France | RQA |
| Cadenza | AXONA/TONIC | Spring | UK | Scaffold |
| Paragon | CSW-1724-19-5-69//Axona/Tonic | Spring | UK | Scaffold |
| Robigus | 1366/Z-836 | Winter | UK | Scaffold |
| Claire | WASP/FLAME | Winter | UK | Scaffold |

**Table S3** - *MoT* isolates used in this study.

| Isolate code | Origin | Date collected | Source |
|---|---|---|---|
| BR32 | Brazil | 1991 | Lesley Boyd, JIC |
| BR48 | Brazil | 1990 | Lesley Boyd, JIC |
| NO6047 | Brazil | 2006 | Diane Saunders, JIC |
| NO6047+*AVR8* | - | - | Diane Saunders, JIC |
| Py 15.1.018 | Brazil | 2015 | Embrapa |
| BTJ4P-1 | Bangladesh | 2016 | Tofazzal Islam, BSMRU, Bangladesh |
| BR48ΔeI | - | - | Yukio Tosa, Kobe University |
| BR48ΔeI+eI | - | - | Yukio Tosa, Kobe University |
| BR48ΔeI+eII | - | - | Yukio Tosa, Kobe University |
| BR48ΔeI+eII' | - | - | Yukio Tosa, Kobe University |

**Table S4** - Mean values of the detached leaf phenotype data for the Watkins core panel screened with NO6047+*AVR8* and Py 15.1.018. JIC GRU Store code refers to the accession identifier in the John Innes Centre Germplasm Resources Unit (www.seedstor.ac.uk).

| JIC GRU store code | Detached leaf assay phenotype (Mean score at 5 dpi) | | Contains 2A association peak |
|---|---|---|---|
| | NO6047+*AVR8* | Py 15.1.018 | |
| WATDE0477 | 0.00 | 0.00 | 2A |
| WATDE0310 | 0.67 | 0.00 | 2A |
| WATDE0571 | 0.67 | 0.00 | 2A |
| WATDE0971 | 0.67 | 0.00 | 2A |
| WATDE0427 | 1.00 | 0.00 | 2A |
| WATDE0973 | 0.33 | 0.33 | 2A |
| WATDE0015 | 2.33 | 0.50 | |
| WATDE0526 | 1.00 | 0.67 | 2A |
| WATDE0102 | 1.00 | 1.00 | 2A |
| WATDE0171 | 0.33 | 1.33 | 2A |
| WATDE0369 | 3.00 | 1.33 | 2A |
| Sears Synth Type | - | 1.33 | |
| WATDE0026 | 3.00 | 1.67 | |
| WATDE0873 | 5.33 | 1.67 | |
| WATDE0779 | 1.67 | 2.00 | |
| WATDE0090 | 2.33 | 2.00 | |
| WATDE0071 | 4.67 | 2.00 | |
| WATDE0062 | - | 2.00 | |
| WATDE0046 | 1.33 | 2.33 | |
| WATDE0795 | 2.33 | 2.33 | |
| WATDE0268 | 2.67 | 2.33 | |
| WATDE0126 | 1.00 | 2.50 | |
| WATDE0381 | 1.33 | 2.67 | |

| | | |
|---|---|---|
| WATDE0051 | 1.67 | 2.67 |
| WATDE0386 | 2.33 | 2.67 |
| WATDE0771 | 2.67 | 2.67 |
| WATDE0077 | 3.67 | 2.67 |
| WATDE0868 | 4.67 | 2.67 |
| WATDE0811 | 2.67 | 3.00 |
| WATDE0765 | 3.33 | 3.00 |
| WATDE0086 | 4.00 | 3.00 |
| WATDE0798 | 4.33 | 3.00 |
| WATDE0761 | 4.67 | 3.00 |
| WATDE0069 | 5.33 | 3.00 |
| WATDE0216 | 2.00 | 3.33 |
| WATDE0519 | 2.00 | 3.33 |
| WATDE0004 | 2.67 | 3.33 |
| WATDE0029 | 3.00 | 3.33 |
| WATDE0827 | 3.00 | 3.33 |
| WATDE0050 | 3.67 | 3.33 |
| WATDE0032 | 4.00 | 3.33 |
| WATDE0070 | 4.00 | 3.33 |
| WATDE0450 | 4.00 | 3.33 |
| WATDE0762 | 4.00 | 3.33 |
| WATDE0773 | 4.33 | 3.33 |
| WATDE0018 | 4.67 | 3.33 |
| WATDE0290 | 5.00 | 3.33 |
| WATDE0788 | 5.00 | 3.33 |
| WATDE0002 | 4.33 | 3.50 |
| WATDE0816 | 4.67 | 3.50 |
| WATDE0749 | 1.67 | 3.67 |

| | | |
|---|---|---|
| WATDE0019 | 2.33 | 3.67 |
| WATDE0075 | 2.67 | 3.67 |
| WATDE0198 | 2.67 | 3.67 |
| WATDE0532 | 2.67 | 3.67 |
| WATDE0929 | 3.00 | 3.67 |
| WATDE0003 | 3.33 | 3.67 |
| WATDE0579 | 3.33 | 3.67 |
| WATDE0898 | 3.33 | 3.67 |
| WATDE0073 | 3.67 | 3.67 |
| WATDE0033 | 4.00 | 3.67 |
| WATDE0074 | 4.00 | 3.67 |
| WATDE0776 | 4.00 | 3.67 |
| WATDE0009 | 4.33 | 3.67 |
| WATDE0385 | 4.33 | 3.67 |
| WATDE0250 | 4.67 | 3.67 |
| WATDE0808 | 4.67 | 3.67 |
| WATDE0253 | 5.00 | 3.67 |
| WATDE0276 | 6.00 | 3.67 |
| WATDE0601 | 1.67 | 4.00 |
| WATDE0405 | 2.00 | 4.00 |
| WATDE0747 | 2.00 | 4.00 |
| WATDE0034 | 3.00 | 4.00 |
| WATDE0192 | 3.00 | 4.00 |
| WATDE0703 | 3.00 | 4.00 |
| WATDE0008 | 3.33 | 4.00 |
| WATDE0581 | 3.33 | 4.00 |
| WATDE0604 | 3.33 | 4.00 |
| WATDE0819 | 3.33 | 4.00 |

| | | |
|---|---|---|
| WATDE0770 | 3.67 | 4.00 |
| WATDE0025 | 4.00 | 4.00 |
| WATDE0076 | 4.00 | 4.00 |
| WATDE0725 | 4.00 | 4.00 |
| WATDE0865 | 4.00 | 4.00 |
| WATDE0156 | 4.33 | 4.00 |
| WATDE0196 | 4.33 | 4.00 |
| WATDE0082 | 4.67 | 4.00 |
| WATDE0089 | 4.67 | 4.00 |
| WATDE0555 | 4.67 | 4.00 |
| WATDE0017 | 5.67 | 4.00 |
| WATDE0278 | 6.00 | 4.00 |
| WATDE0058 | 0.33 | 4.33 |
| WATDE0241 | 2.33 | 4.33 |
| WATDE0702 | 2.33 | 4.33 |
| WATDE0238 | 2.67 | 4.33 |
| WATDE0669 | 2.67 | 4.33 |
| WATDE0067 | 3.00 | 4.33 |
| WATDE0010 | 3.33 | 4.33 |
| WATDE0049 | 3.33 | 4.33 |
| WATDE0027 | 3.67 | 4.33 |
| WATDE0542 | 3.67 | 4.33 |
| WATDE0705 | 3.67 | 4.33 |
| WATDE0392 | 4.00 | 4.33 |
| WATDE0694 | 4.00 | 4.33 |
| CIMCOG 32 | 4.33 | 4.33 |
| WATDE0021 | 4.33 | 4.33 |
| WATDE0518 | 4.33 | 4.33 |

| | | |
|---|---|---|
| WATDE0634 | 4.33 | 4.33 |
| WATDE0892 | 4.33 | 4.33 |
| WATDE0068 | 4.67 | 4.33 |
| WATDE0078 | 4.67 | 4.33 |
| WATDE0294 | 4.67 | 4.33 |
| WATDE0435 | 4.67 | 4.33 |
| WATDE0558 | 4.67 | 4.33 |
| WATDE0782 | 5.33 | 4.33 |
| WATDE0934 | 5.33 | 4.33 |
| WATDE0028 | 5.67 | 4.33 |
| WATDE0375 | 6.00 | 4.33 |
| WATDE0774 | 6.00 | 4.33 |
| WATDE0001 | 3.67 | 4.50 |
| WATDE0864 | 5.33 | 4.50 |
| WATDE0066 | 1.67 | 4.67 |
| WATDE0984 | 2.00 | 4.67 |
| WATDE0222 | 3.00 | 4.67 |
| WATDE0305 | 3.00 | 4.67 |
| WATDE0732 | 3.33 | 4.67 |
| WATDE0083 | 3.67 | 4.67 |
| WATDE0336 | 4.00 | 4.67 |
| WATDE0476 | 4.00 | 4.67 |
| WATDE0843 | 4.00 | 4.67 |
| WATDE0081 | 4.33 | 4.67 |
| WATDE0180 | 4.33 | 4.67 |
| WATDE0743 | 4.33 | 4.67 |
| CIMCOG 26 | 4.67 | 4.67 |
| WATDE0030 | 4.67 | 4.67 |

| | | |
|---|---|---|
| WATDE0057 | 4.67 | 4.67 |
| WATDE0445 | 4.67 | 4.67 |
| WATDE0576 | 4.67 | 4.67 |
| WATDE0831 | 4.67 | 4.67 |
| WATDE0950 | 4.67 | 4.67 |
| WATDE0339 | 5.00 | 4.67 |
| WATDE0919 | 5.00 | 4.67 |
| WATDE0056 | 5.33 | 4.67 |
| WATDE0098 | 5.33 | 4.67 |
| WATDE0108 | 5.33 | 4.67 |
| WATDE0382 | 5.67 | 4.67 |
| WATDE0461 | 5.67 | 4.67 |
| WATDE0712 | 6.00 | 4.67 |
| WATDE0679 | 2.33 | 5.00 |
| WATDE0668 | 3.00 | 5.00 |
| WATDE0751 | 3.00 | 5.00 |
| WATDE0457 | 3.33 | 5.00 |
| WATDE0708 | 3.33 | 5.00 |
| WATDE0005 | 3.67 | 5.00 |
| WATDE0052 | 3.67 | 5.00 |
| WATDE0072 | 3.67 | 5.00 |
| WATDE0582 | 3.67 | 5.00 |
| WATDE0037 | 4.00 | 5.00 |
| WATDE0737 | 4.00 | 5.00 |
| WATDE0020 | 4.33 | 5.00 |
| WATDE0047 | 4.33 | 5.00 |
| WATDE0096 | 4.33 | 5.00 |
| WATDE0740 | 4.33 | 5.00 |

| | | |
|---|---|---|
| WATDE0991 | 4.33 | 5.00 |
| WATDE0013 | 4.67 | 5.00 |
| WATDE0963 | 4.67 | 5.00 |
| WATDE0801 | 5.00 | 5.00 |
| WATDE0863 | 5.00 | 5.00 |
| WATDE0574 | 5.33 | 5.00 |
| WATDE0861 | 5.33 | 5.00 |
| WATDE0939 | 5.33 | 5.00 |
| WATDE1052 | 5.33 | 5.00 |
| WATDE0053 | 5.67 | 5.00 |
| WATDE0308 | 5.67 | 5.00 |
| WATDE0699 | 5.67 | 5.00 |
| WATDE0791 | 6.00 | 5.00 |
| WATDE0932 | 6.00 | 5.00 |
| WATDE0937 | 6.00 | 5.00 |
| Wyalkatchem | - | 5.00 |
| WATDE0613 | 3.00 | 5.33 |
| WATDE0011 | 3.33 | 5.33 |
| WATDE0104 | 3.33 | 5.33 |
| WATDE0042 | 3.67 | 5.33 |
| WATDE0007 | 4.00 | 5.33 |
| WATDE0040 | 4.00 | 5.33 |
| WATDE0594 | 4.00 | 5.33 |
| WATDE0048 | 4.33 | 5.33 |
| WATDE0664 | 4.33 | 5.33 |
| WATDE1051 | 4.33 | 5.33 |
| WATDE0024 | 4.67 | 5.33 |
| WATDE0041 | 4.67 | 5.33 |

| | | |
|---|---|---|
| WATDE0249 | 4.67 | 5.33 |
| WATDE0359 | 4.67 | 5.33 |
| WATDE0588 | 4.67 | 5.33 |
| WATDE0954 | 4.67 | 5.33 |
| Sup152 CIMCOG39 | 5.00 | 5.33 |
| WATDE0031 | 5.00 | 5.33 |
| WATDE0100 | 5.00 | 5.33 |
| WATDE0714 | 5.00 | 5.33 |
| WATDE0871 | 5.00 | 5.33 |
| Weebil | 5.00 | 5.33 |
| WATDE0060 | 5.33 | 5.33 |
| WATDE0112 | 5.33 | 5.33 |
| WATDE0061 | 5.67 | 5.33 |
| WATDE0036 | 6.00 | 5.33 |
| WATDE0262 | 6.00 | 5.33 |
| WATDE0099 | 3.67 | 5.50 |
| WATDE0456 | 4.67 | 5.50 |
| WATDE0088 | 5.33 | 5.50 |
| WATDE0911 | 6.00 | 5.50 |
| WATDE0266 | 2.33 | 5.67 |
| WATDE0596 | 2.67 | 5.67 |
| WATDE0758 | 3.00 | 5.67 |
| WATDE0611 | 3.33 | 5.67 |
| CIMCOG 47 | 3.67 | 5.67 |
| WATDE0038 | 3.67 | 5.67 |
| WATDE0509 | 4.00 | 5.67 |
| WATDE0678 | 4.00 | 5.67 |
| WATDE0022 | 4.33 | 5.67 |

| | | |
|---|---|---|
| WATDE0055 | 4.33 | 5.67 |
| WATDE0335 | 4.33 | 5.67 |
| WATDE0430 | 4.33 | 5.67 |
| WATDE0451 | 4.67 | 5.67 |
| WATDE0646 | 4.67 | 5.67 |
| WATDE0833 | 4.67 | 5.67 |
| WATDE0054 | 5.00 | 5.67 |
| WATDE0612 | 5.00 | 5.67 |
| WATDE0643 | 5.00 | 5.67 |
| WATDE0016 | 5.33 | 5.67 |
| WATDE0585 | 5.33 | 5.67 |
| WATDE0857 | 5.33 | 5.67 |
| WATDE0094 | 5.67 | 5.67 |
| WATDE0106 | 5.67 | 5.67 |
| WATDE0455 | 5.67 | 5.67 |
| WATDE0479 | 5.67 | 5.67 |
| WATDE0986 | 5.67 | 5.67 |
| WATDE0105 | 6.00 | 5.67 |
| WATDE0635 | 6.00 | 5.67 |
| WATDE0835 | 6.00 | 5.67 |
| WATDE0938 | 6.00 | 5.67 |
| WATDE0133 | 3.67 | 6.00 |
| WATDE0550 | 3.67 | 6.00 |
| WATDE0651 | 3.67 | 6.00 |
| WATDE0670 | 3.67 | 6.00 |
| WATDE0107 | 4.00 | 6.00 |
| WATDE0149 | 4.00 | 6.00 |
| WATDE0035 | 4.33 | 6.00 |

| | | |
|---|---|---|
| WATDE0039 | 4.33 | 6.00 |
| WATDE0101 | 4.33 | 6.00 |
| WATDE0263 | 4.33 | 6.00 |
| WATDE0521 | 4.33 | 6.00 |
| WATDE0592 | 4.33 | 6.00 |
| WATDE0882 | 4.33 | 6.00 |
| WATDE0989 | 4.33 | 6.00 |
| WATDE1012 | 4.33 | 6.00 |
| WATDE0023 | 4.50 | 6.00 |
| CIMCOG 03 | 4.67 | 6.00 |
| CIMCOG 56 | 4.67 | 6.00 |
| WATDE0092 | 4.67 | 6.00 |
| WATDE0095 | 4.67 | 6.00 |
| WATDE0228 | 4.67 | 6.00 |
| WATDE0422 | 4.67 | 6.00 |
| WATDE0459 | 4.67 | 6.00 |
| WATDE1027 | 4.67 | 6.00 |
| CIMCOG 12 | 5.00 | 6.00 |
| Fielder | 5.00 | 6.00 |
| WATDE0113 | 5.00 | 6.00 |
| WATDE0138 | 5.00 | 6.00 |
| WATDE0215 | 5.00 | 6.00 |
| WATDE0292 | 5.00 | 6.00 |
| WATDE0639 | 5.00 | 6.00 |
| WATDE0909 | 5.00 | 6.00 |
| WATDE0993 | 5.00 | 6.00 |
| Baj | 5.33 | 6.00 |
| Chinese Spring | 5.33 | 6.00 |

| | | |
|---|---|---|
| Paragon | 5.33 | 6.00 |
| WATDE0091 | 5.33 | 6.00 |
| WATDE0110 | 5.33 | 6.00 |
| WATDE0116 | 5.33 | 6.00 |
| WATDE0420 | 5.33 | 6.00 |
| WATDE0534 | 5.33 | 6.00 |
| WATDE0557 | 5.33 | 6.00 |
| WATDE0609 | 5.33 | 6.00 |
| WATDE0631 | 5.33 | 6.00 |
| WATDE0661 | 5.33 | 6.00 |
| WATDE0671 | 5.33 | 6.00 |
| Waxwing | 5.33 | 6.00 |
| CIMCOG 49 | 5.67 | 6.00 |
| CIMCOG 53 | 5.67 | 6.00 |
| MISR1 CIMCOG33 | 5.67 | 6.00 |
| WATDE0093 | 5.67 | 6.00 |
| WATDE0097 | 5.67 | 6.00 |
| WATDE0111 | 5.67 | 6.00 |
| WATDE0394 | 5.67 | 6.00 |
| WATDE0577 | 5.67 | 6.00 |
| WATDE0617 | 5.67 | 6.00 |
| WATDE0659 | 5.67 | 6.00 |
| WATDE0883 | 5.67 | 6.00 |
| WATDE1025 | 5.67 | 6.00 |
| WATDE1030 | 5.67 | 6.00 |
| Pamyat Azieva | 6.00 | 6.00 |
| WATDE0080 | 6.00 | 6.00 |
| WATDE0087 | 6.00 | 6.00 |

| | | |
|---|---|---|
| WATDE0114 | 6.00 | 6.00 |
| WATDE0115 | 6.00 | 6.00 |
| WATDE0117 | 6.00 | 6.00 |
| WATDE0118 | 6.00 | 6.00 |
| WATDE0396 | 6.00 | 6.00 |
| WATDE0397 | 6.00 | 6.00 |
| WATDE0486 | 6.00 | 6.00 |
| WATDE0653 | 6.00 | 6.00 |
| WATDE0888 | 6.00 | 6.00 |
| WATDE0895 | 6.00 | 6.00 |
| WATDE1026 | 6.00 | 6.00 |
| WATDE1060 | 6.00 | 6.00 |
| WATDE0063 | 4.00 | - |
| Reedling | 5.00 | - |
| WATDE0727 | 5.00 | - |
| WATDE0064 | 5.33 | - |
| Pfau | 6.00 | - |

**Table S5** - Additional Watkins cultivars used to verify the 2A resistance, their origins, growth habit and detached leaf (DLA) and spike (DSA) assay phenotype at five days post inoculation with isolate Py 15.1.018. Phenotype was predicted based on the presense of 'Region 1' or 'Region 2' following haplotype analysis.

| GRU Store Code | Collection ID | Origin Country | Growth Habit | Predicted phenotype | Mean DLA score (5 dpi) | Mean DSA score (5 dpi) |
|---|---|---|---|---|---|---|
| WATDE0056 | Wat1190433-1 | India | Spring | Resistant | 5.20 | 4.90 |
| WATDE0426 | Wat1190267-1 | Spain | Spring | Resistant | 1.00 | 0.70 |
| WATDE0428 | Wat1190269-1 | Spain | Spring | Resistant | 1.00 | 2.70 |
| WATDE0465 | Wat1190309-1 | Iran | Winter | Resistant | 1.20 | - |
| WATDE0484 | Wat1190320-3 | China | Winter | Resistant | 1.60 | 3.00 |
| WATDE0505 | Wat1190337-1 | Hungary | Spring | Resistant | 1.20 | 5.00 |
| WATDE0527 | Wat1190357-2 | Yugoslavia | Spring | Resistant | 5.00 | 5.70 |
| WATDE0541 | Wat1190369-2 | Yugoslavia | Spring | Resistant | 0.80 | 2.70 |
| WATDE0546 | Wat1190373-1 | Iran | Spring | Resistant | 0.60 | 2.30 |
| WATDE0566 | Wat1190389-1 | Portugal | Spring | Resistant | 1.20 | 2.00 |
| WATDE0567 | Wat1190390-1 | Portugal | Spring | Resistant | 1.20 | 4.00 |
| WATDE0568 | Wat1190391-1 | Portugal | Spring | Resistant | 5.40 | 6.00 |
| WATDE0592 | Wat1190412-1 | India | Spring | Resistant | 6.00 | 6.00 |
| WATDE0672 | Wat1190488-1 | USSR | Winter | Resistant | 1.20 | 2.50 |
| WATDE0687 | Wat1190500-1 | Iraq | Winter | Resistant | 0.80 | 3.00 |
| WATDE0720 | Wat1190525-1 | India | Spring | Resistant | 5.40 | 6.00 |
| WATDE0122 | Wat1190003-4 | Iran | Spring | Susceptible | 5.60 | 5.30 |
| WATDE0141 | Wat1190017-2 | Spain | Winter | Susceptible | 5.40 | 6.00 |
| WATDE0152 | Wat1190027-1 | Australia | Spring | Susceptible | 5.60 | - |
| WATDE0162 | Wat1190037-1 | Poland | Winter | Susceptible | 6.00 | 6.00 |
| WATDE0175 | Wat1190049-1 | Spain | Spring | Susceptible | 6.00 | 6.00 |
| WATDE0185 | Wat1190058-1 | Portugal | Spring | Susceptible | 5.60 | - |

| | | | | | | |
|---|---|---|---|---|---|---|
| WATDE0199 | Wat1190068-1 | Spain | Spring | Susceptible | 5.80 | - |
| WATDE0207 | Wat1190075-1 | Yugoslavia | Winter | Susceptible | 6.00 | 5.70 |
| WATDE0220 | Wat1190086-2 | India | Spring | Susceptible | 4.60 | - |
| WATDE0232 | Wat1190097-1 | Poland | Winter | Susceptible | 6.00 | 4.80 |
| WATDE0804 | Wat1190601-1 | Spain | Spring | Resistant | 1.60 | 4.00 |
| WATDE0970 | Wat1190758-2 | Italy | Spring | Resistant | 1.00 | 2.70 |
| WATDE0974 | Wat1190761-1 | USSR | Spring | Resistant | 1.00 | 3.70 |
| WATDE1062 | Wat1190911-1 | Hungary | Spring | Resistant | 1.20 | 2.70 |

**Table S6** - Adapted wheat varieties used in this study, along with their detached leaf assay phenotype using NO6047+*AVR8* and Py 15.1.018 isolates. Scores were taken at six days post inoculation.

| Accession | JIC GRU store code | Detached leaf assay with NO6047+*AVR8* | Detached leaf assay with Py 15.1.018 |
|---|---|---|---|
| Ability | WGED0562 | 2.33 | 1.00 |
| Boxer | WGED0540 | 1.67 | 1.00 |
| Claire | PANG0005 | 1.20 | 1.20 |
| Consort | WGED0670 | 3.00 | 3.00 |
| Cordiale | W10003 | - | - |
| Crest | WGED0666 | 3.00 | 2.00 |
| Encore | WGED0671 | 2.33 | 2.00 |
| Flame | WGED0155 | 2.33 | 1.00 |
| Grafton | - | - | - |
| Holster | WGED0663 | 3.00 | 3.00 |
| Kosack | - | - | - |
| Legend | WGED0561 | 1.33 | 1.33 |
| Malacca | WGED0639 | 3.33 | 2.00 |
| SY-Mattis | PANG0015 | 1.80 | 1.20 |
| Reform | - | - | - |
| Renan | WGED0248 | 2.00 | 0.80 |
| Rendevous | - | 2.60 | 1.80 |
| Revelation | W10190 | 1.40 | 1.20 |
| Riband | WGED0128 | 3.00 | 1.67 |
| Savannah | WGED0280 | 2.67 | 1.67 |
| Shango | WGED0713 | 2.33 | 1.67 |
| Spark | WGED0159 | 0.67 | 1.33 |
| Spitfire | WGED0712 | 2.00 | 1.67 |
| CDC Stanley | PANG0004 | 2.60 | 1.20 |

| | | | |
|---|---|---|---|
| Stava | WGED0211 | 1.67 | 1.00 |
| SY-Epson | W10079 | 1.20 | 0.00 |
| Torfrida | WGED0161 | 3.00 | 1.33 |
| Turpin | WGED0629 | 3.67 | 1.67 |
| Wasp | WGED0630 | 2.00 | 1.00 |

**Table S7 -** High confidence gene content of the SY-Mattis 2A interval, their location and function from the *de novo* genome annotation. Gene content of the 400kb SY-Mattis haplotype (78855000 to 788950000 kb) is highlighted in yellow.

| Gene code | Coordinates (orientation) | Function |
|---|---|---|
| TraesSYM2A03G00828360 | 788728552 - 788738447(+) | Receptor-like kinase |
| TraesSYM2A03G00828370 | 788768076 - 788769438(+) | *unknown* |
| TraesSYM2A03G00828380 | 788811120 - 788813175(+) | *unknown* |
| TraesSYM2A03G00828390 | 788825472 - 788828016(+) | *unknown* |
| TraesSYM2A03G00828400 | 788828264 - 788832085(+) | methyltransferases superfamily protein |
| TraesSYM2A03G00828410 | 788833791 - 788837343(+) | RNase P 1 |
| TraesSYM2A03G00828420 | 788837936 - 788839387(+) | STAY-GREEN LIKE, chloroplastic |
| TraesSYM2A03G00828440 | 788841216 - 788842285(+) | *unknown* |
| TraesSYM2A03G00828450 | 788845950 - 788849102(+) | resistance protein RGA2 |
| TraesSYM2A03G00828460 | 788885045 - 788890732(+) | resistance protein RGA2 |
| TraesSYM2A03G00828500 | 788960342 - 788962593(+) | *unknown* |
| TraesSYM2A03G00828540 | 789110507 - 789111037(+) | *unknown* |
| TraesSYM2A03G00828580 | 789151116 - 789153155(+) | *unknown* |
| TraesSYM2A03G00828590 | 789161156 - 789162436(+) | F-box/kelch-repeat protein |
| TraesSYM2A03G00828600 | 789188895 - 789193763(+) | resistance protein RGA2/ transposon TNT |
| TraesSYM2A03G00828640 | 789217066 - 789233090(+) | F-box/kelch-repeat protein |
| TraesSYM2A03G00828700 | 789264657 - 789265305(-) | protein 5NG4 |
| TraesSYM2A03G00828720 | 789284536 - 789289459(+) | resistance protein RGA2 |
| TraesSYM2A03G00828810 | 789390224 - 789396435(+) | resistance protein RGA2 |
| TraesSYM2A03G00828830 | 789403251 - 789408104(+) | resistance protein RGA2 |

| | | |
|---|---|---|
| TraesSYM2A03G00828840 | 789408643 - 789409011(-) | peptidase subunit alpha |
| TraesSYM2A03G00828850 | 789416992 - 789419026(+) | phosphodiesterases superfamily protein |
| TraesSYM2A03G00828930 | 789745589 - 789748504(-) | 4-like protein |
| TraesSYM2A03G00829020 | 790122656 - 790126161(-) | *unknown* |
| TraesSYM2A03G00829080 | 790381831 - 790387386(+) | *unknown* |
| TraesSYM2A03G00829120 | 790568231 - 790571224(-) | *unknown* |
| TraesSYM2A03G00829200 | 790955915 - 790959358(-) | *unknown* |
| TraesSYM2A03G00829250 | 791221535 - 791225005(+) | *unknown* |
| TraesSYM2A03G00829350 | 791855004 - 791858879(-) | *unknown* |
| TraesSYM2A03G00829450 | 792130947 - 792134542(+) | *unknown* |
| TraesSYM2A03G00829510 | 792400216 - 792404527(-) | N-methyltransferase SUVR4 |
| TraesSYM2A03G00829520 | 792405026 - 792406692(+) | non-LTR retrotransposon |
| TraesSYM2A03G00829540 | 792410750 - 792414847(-) | glycosyltransferase subunit 1 |
| TraesSYM2A03G00829560 | 792465474 - 792467366(-) | beta-1,6-N-acetylglucosaminyltransferase family protein |
| TraesSYM2A03G00829570 | 792472433 - 792473359(+) | *unknown* |
| TraesSYM2A03G00829580 | 792519361 - 792525805(-) | toxin-like protein Hfr-2 |
| TraesSYM2A03G00829600 | 792527494 - 792528414(-) | *unknown* |
| TraesSYM2A03G00829610 | 792534358 - 792535178(+) | *unknown* |
| TraesSYM2A03G00829620 | 792554319 - 792557912(+) | domain-containing protein 78 |
| TraesSYM2A03G00829630 | 792559869 - 792562916(+) | protein Rab-18 |
| TraesSYM2A03G00829640 | 792660864 - 792662136(+) | *unknown* |
| TraesSYM2A03G00829650 | 792660869 - 792662507(-) | *unknown* |
| TraesSYM2A03G00829660 | 792670265 - 792673854(+) | resistance protein RGA2 |
| TraesSYM2A03G00829670 | 792679164 - 792680963(+) | Receptor-like kinase family |

| | | |
|---|---|---|
| TraesSYM2A03G00829710 | 792724770 - 792728735(-) | protein, putative, Ty1-copia |
| TraesSYM2A03G00829740 | 792765879 - 792768798(+) | *unknown* |
| TraesSYM2A03G00829750 | 792781806 - 792786738(+) | resistance protein RGA2 |
| TraesSYM2A03G00829800 | 792879278 - 792881875(-) | Protease |
| TraesSYM2A03G00829810 | 792928596 - 792934980(+) | Peptidase |
| TraesSYM2A03G00829820 | 792935138 - 792938913(-) | transporter NIPA (DUF803) |
| TraesSYM2A03G00829830 | 792983207 - 792991299(+) | *unknown* |
| TraesSYM2A03G00829850 | 793062876 - 793066852(+) | Putative protein, Ty1-copia |
| TraesSYM2A03G00830090 | 793345006 - 793347118(+) | *unknown* |
| TraesSYM2A03G00830110 | 793359286 - 793361111(+) | domain-containing protein 78 |
| TraesSYM2A03G00830160 | 793436071 - 793438232(-) | *unknown* |
| TraesSYM2A03G00830190 | 793470556 - 793474718(+) | ATP-dependent RNA helicase |
| TraesSYM2A03G00830210 | 793492346 - 793493598(+) | *unknown* |
| TraesSYM2A03G00830260 | 793585072 - 793585649(-) | *unknown* |
| TraesSYM2A03G00830280 | 793602733 - 793603198(-) | ATPase sarcoplasmic/ERCC-6-like protein |
| TraesSYM2A03G00830300 | 793618964 - 793626781(+) | excision repair protein |
| TraesSYM2A03G00830400 | 793840966 - 793844595(-) | *unknown* |
| TraesSYM2A03G00830410 | 793848007 - 793848435(+) | *unknown* |
| TraesSYM2A03G00830420 | 793853596 - 793855385(+) | *unknown* |
| TraesSYM2A03G00830430 | 793857902 - 793860706(-) | transcription factor 28 |
| TraesSYM2A03G00830440 | 793864325 - 793867928(-) | transcription factor 28 |
| TraesSYM2A03G00830450 | 793882890 - 793884880(+) | transcription factor 28 |
| TraesSYM2A03G00830490 | 793986412 - 793987375(-) | *unknown* |
| TraesSYM2A03G00830530 | 794021318 - 794024342(-) | phosphodiesterases superfamily protein |
| TraesSYM2A03G00830550 | 794036223 - 794041272(-) | resistance protein RGA2 - related family |

| | | |
|---|---|---|
| TraesSYM2A03G00830580 | 794071710 - 794073655(-) | putative protein, Mutator subclass |
| TraesSYM2A03G00830590 | 794082632 - 794085547(+) | protein putative, Ty1-copia |
| TraesSYM2A03G00830600 | 794135460 - 794136365(-) | SKIP23-like protein |
| TraesSYM2A03G00830620 | 794148636 - 794150360(+) | glucosyltransferase 1 |

**Table S8** - Readcounts for the high confidence gene content of the 400kb SY-Mattis interval.

| Gene code* | Coordinates (orientation) | Readcounts for dataset 1 | Readcounts for dataset 2 |
|---|---|---|---|
| TraesSYM2A03G00828360 | 788728552 - 788738447(+) | 1224 | 1274 |
| TraesSYM2A03G00828370 | 788768076 - 788769438(+) | 0 | 0 |
| TraesSYM2A03G00828380 | 788811120 - 788813175(+) | 0 | 0 |
| TraesSYM2A03G00828390 | 788825472 - 788828016(+) | 0 | 0 |
| TraesSYM2A03G00828400 | 788828264 - 788832085(+) | 1264 | 1172 |
| TraesSYM2A03G00828410 | 788833791 - 788837343(+) | 2202 | 2326 |
| TraesSYM2A03G00828420 | 788837936 - 788839387(+) | 8930 | 6822 |
| TraesSYM2A03G00828440 | 788841216 - 788842285(+) | 0 | 0 |
| TraesSYM2A03G00828450 | 788845950 - 788849102(+) | 0 | 0 |
| TraesSYM2A03G00828460 | 788885045 - 788890732(+) | 106 | 128 |

**Table S9** - Polymorphisms present in the nucleotide sequences of TraesSYM2A03G00828400, TraesSYM2A03G00828410, TraesSYM2A03G00828420, TraesSYM2A03G00828450 and TraesSYM2A03G00828460 for all adapted wheat varieties and Watkins accessions where alignments were generated. Sequences were mapped to SY-Mattis. Scores were taken at five days post inoculation from detached leaves.

| | | | SY-Mattis Chr2A | | | | | | | |
|---|---|---|---|---|---|---|---|---|---|---|
| | | | *TraesSYM2A03G00828400* | | *TraesSYM2A03G00828410* | *TraesSYM2A03G00828420* | *TraesSYM2A03G00828450* | *TraesSYM2A03G00828460* | | |
| Cultivar | *NO6047+ AVR8* | Py 15.1.018 | Exon 1 788,828,724 | Exon 5 788,830,642 | | Exon 1 788,838,013 | | 788,888,585 | 788,889,012 | 788,890,436 |
| Ability | 2.33 | 1.00 | - | - | - | - | - | - | T | - |
| Claire | 1.20 | 1.20 | - | - | - | - | - | - | T | - |
| SY-Epson | 1.20 | 0.00 | - | - | - | - | - | - | - | - |
| Extase | | | - | - | - | - | - | - | T | - |
| Cordiale | | | - | - | - | - | - | - | - | - |
| Grafton | | | - | - | - | - | - | - | T | - |
| Flame | 2.33 | 1.00 | - | - | - | - | - | - | T | - |
| Malacca | 3.33 | 2.00 | - | - | - | - | - | - | T | - |
| Renan | 2.00 | 0.80 | - | - | - | - | - | - | - | - |
| Revelation | 1.40 | 1.20 | - | - | - | - | - | - | T | - |
| Riband | 3.00 | 1.67 | - | - | - | - | - | - | T | - |
| Shango | 2.33 | 1.67 | - | - | - | - | - | - | T | - |
| Spark | 0.67 | 1.33 | - | - | - | - | - | - | T | - |
| SY-Mattis | 1.80 | 1.20 | - | - | - | - | - | - | - | - |
| Wasp | 2.00 | 1.00 | - | - | - | - | - | - | T | - |
| WATDE0048 | 4.60 | 5.60 | - | - | - | - | - | - | - | - |
| WATDE0102 | 2.60 | 2.20 | - | - | - | - | - | - | T | - |
| WATDE0171 | 1.80 | 0.80 | - | - | - | - | - | - | T | - |
| WATDE0310 | 1.20 | 0.80 | - | - | - | - | - | - | T | - |
| WATDE0369 | 3.00 | 1.30 | - | - | - | - | - | - | - | - |
| WATDE0426 | | 1.00 | - | - | - | - | - | T | - | - |
| WATDE0427 | 1.00 | 0.00 | - | - | - | - | - | T | - | - |
| WATDE0428 | | 1.00 | - | - | - | - | - | T | - | - |
| WATDE0465 | | 1.20 | - | - | - | - | - | - | - | - |
| WATDE0477 | 0.00 | 0.00 | - | A | - | G | - | - | - | - |
| WATDE0505 | | 1.20 | - | - | - | - | - | - | - | - |
| WATDE0526 | 1.00 | 0.67 | - | - | - | - | - | - | - | - |
| WATDE0527 | 4.60 | 5.80 | - | - | - | - | - | - | - | - |
| WATDE0541 | | 0.80 | - | - | - | - | - | - | - | - |
| WATDE0546 | | 1.20 | - | - | - | - | - | - | - | - |
| WATDE0566 | 1.60 | 2.20 | - | - | - | G | - | - | - | - |
| WATDE0567 | | 1.20 | - | - | - | - | - | - | - | - |
| WATDE0568 | 5.60 | 5.80 | - | - | - | - | - | - | - | - |
| WATDE0571 | 1.80 | 1.00 | - | - | - | - | - | - | - | - |
| WATDE0592 | 4.60 | 5.60 | - | - | - | - | - | - | - | - |
| WATDE0672 | | 1.20 | T | - | - | - | - | - | - | - |
| WATDE0687 | | 0.80 | T | - | - | - | - | - | - | - |
| WATDE0804 | 0.80 | 1.80 | - | - | - | G | - | - | - | - |
| WATDE0970 | | 1.00 | T* | - | - | - | - | - | - | - |
| WATDE0971 | 0.67 | 0.00 | - | - | - | - | - | - | - | - |
| WATDE0973 | 0.33 | 0.30 | - | - | - | - | - | - | - | A |
| WATDE0974 | | 1.00 | - | - | - | - | - | - | - | A |
| WATDE1062 | | 1.20 | - | - | - | - | - | - | - | - |

* only 1 read coverage

**Table S10** - Non-synonymous polymorphisms present in the ORF of *Pm4* for all adapted wheat varieties and Watkins accessions where alignments were generated. The codon position for each polymorphism is shown in the third row. Claire and SY-Mattis were also included. Sequences were mapped to SY-Mattis. Scores were taken at five days post inoculation from detached leaves.

| | | | *TraesSYM2A03G00828360 (Pm4)* | | | | | | | | | | | | | | |
|---|---|---|---|---|---|---|---|---|---|---|---|---|---|---|---|---|---|
| | | | EXON 1 | | | EXON 3 | | | EXON 7 | | | | | | | | |
| | Phenotype | | A50E | | | E205K | | | W446X | | | V697V | | | A713G | | |
| | | | ala - glu | | | glu - lys | | | trp - STOP | | | val - val | | | ala - gly | | |
| Cultivar | NO6047+ *AVR8* | Py 15.1.018 | G | C | A | G | A | A | T | G | G | G | T | C | G | C | C |
| Ability | 2.33 | 1.00 | - | - | - | - | - | - | - | - | - | - | - | G | - | G | - |
| Claire | 1.20 | 1.20 | - | - | - | - | - | - | - | - | - | - | - | G | - | G | - |
| SY-Epson | 1.20 | 0.00 | - | - | - | - | - | - | - | - | - | - | - | | - | | - |
| Flame | 2.33 | 1.00 | - | - | - | - | - | - | - | - | - | - | - | G | - | G | - |
| Malacca | 3.33 | 2.00 | - | - | - | - | - | - | - | - | - | - | - | G | - | G | - |
| Renan | 2.00 | 0.80 | - | - | - | - | - | - | - | - | - | - | - | | - | | - |
| Revelation | 1.40 | 1.20 | - | - | - | - | - | - | - | - | - | - | - | G | - | G | - |
| Riband | 3.00 | 1.67 | - | - | - | - | - | - | - | - | - | - | - | G | - | G | - |
| Shango | 2.33 | 1.67 | - | - | - | - | - | - | - | - | - | - | - | G | - | G | - |
| Spark | 0.67 | 1.33 | - | - | - | - | - | - | - | - | - | - | - | G | - | G | - |
| SY-Mattis | 1.80 | 1.20 | - | - | - | - | - | - | - | - | - | - | - | - | - | - | - |
| Wasp | 2.00 | 1.00 | - | - | - | - | - | - | - | - | - | - | - | G | - | G | - |
| WATDE0048 | 4.60 | 5.60 | - | A | - | A | - | - | - | - | - | - | - | - | - | - | - |
| WATDE0102 | 2.60 | 2.20 | - | - | - | - | - | - | - | - | - | - | - | G | - | G | - |
| WATDE0171 | 1.80 | 0.80 | - | - | - | - | - | - | - | - | - | - | - | G | - | G | - |
| WATDE0310 | 1.20 | 0.80 | - | - | - | - | - | - | - | - | - | - | - | G | - | G | - |
| WATDE0369 | 3.00 | 1.30 | - | - | - | A | - | - | - | - | - | - | - | - | - | - | - |
| WATDE0426 | | 1.00 | - | - | - | A | - | - | - | - | - | - | - | - | - | - | - |
| WATDE0427 | 1.00 | 0.00 | - | - | - | A | - | - | - | - | - | - | - | - | - | - | - |
| WATDE0428 | | 1.00 | - | - | - | A | - | - | - | - | - | - | - | - | - | - | - |
| WATDE0465 | | 1.20 | - | - | - | A | - | - | - | - | - | - | - | - | - | - | - |
| WATDE0477 | 0.00 | 0.00 | - | - | - | A | - | - | - | - | - | - | - | - | - | - | - |
| WATDE0505 | | 1.20 | - | - | - | A | - | - | - | - | - | - | - | - | - | - | - |
| WATDE0526 | 1.00 | 0.67 | - | - | - | A | - | - | - | - | - | - | - | - | - | - | - |
| WATDE0527 | 4.60 | 5.80 | - | A | - | - | - | - | - | - | - | - | - | - | - | - | - |
| WATDE0541 | | 0.80 | - | - | - | A | - | - | - | - | - | - | - | - | - | - | - |
| WATDE0546 | | 1.20 | - | - | - | A | - | - | - | - | - | - | - | - | - | - | - |
| WATDE0566 | 1.60 | 2.20 | - | - | - | A | - | - | - | - | - | - | - | - | - | - | - |
| WATDE0567 | | 1.20 | - | - | - | A | - | - | - | - | - | - | - | - | - | - | - |
| WATDE0568 | 5.60 | 5.80 | - | - | - | A | - | - | - | A | - | - | - | - | - | - | - |
| WATDE0571 | 1.80 | 1.00 | - | - | - | A | - | - | - | - | - | - | - | - | - | - | - |
| WATDE0592 | 4.60 | 5.60 | - | - | - | A | - | - | - | A | - | - | - | - | - | - | - |
| WATDE0672 | | 1.20 | - | - | - | A | - | - | - | - | - | - | - | - | - | - | - |
| WATDE0687 | | 0.80 | - | - | - | A | - | - | - | - | - | - | - | - | - | - | - |
| WATDE0804 | 0.80 | 1.80 | - | - | - | A | - | - | - | - | - | - | - | - | - | - | - |
| WATDE0970 | | 1.00 | - | - | - | A | - | - | - | - | - | - | - | - | - | - | - |
| WATDE0971 | 0.67 | 0.00 | - | - | - | A | - | - | - | - | - | - | - | - | - | - | - |
| WATDE0973 | 0.33 | 0.30 | - | - | - | A | - | - | - | - | - | - | - | - | - | - | - |
| WATDE0974 | | 1.00 | - | - | - | A | - | - | - | - | - | - | - | - | - | - | - |
| WATDE1062 | | 1.20 | - | - | - | A | - | - | - | - | - | - | - | - | - | - | - |

**Table S11** - Near-isogenic lines (NIL), EMS-induced mutant and overexpression lines used to validate the recognition of *AVR-Rmg8* by *Pm4.*

| **Line** | **Description** | **DLA phenotype** | | | | **Reference** |
|---|---|---|---|---|---|---|
| | | **NO6047+*Avr8*** | **Py 15.1.018** | **Br48ΔeI** | **Br48ΔeI+eI** | |
| **Bobwhite S26** | Susceptible background for overexpression transformants | S | S | S | S | - |
| **Federation** | Susceptible background for NILs | S | S | S | S | - |
| ***Fed-Pm4a*** **NIL** | *Pm4a* from Khapli introgressed into Chancellor, then Federation | R | R | S | R | McIntosh and Bennet, 1979 |
| ***Fed-Pm4b*** **NIL** | *Pm4b* from *T. carthlicum* introgressed into W804, then Federation*. Resistant background for Pm4b* EMS mutants. | R | R | S | R | Briggle, 1966 |
| ***Pm4b*_mutant_123** | EMS mutant (G132D) in *Fed-Pm4b* NIL background | S | S | S | S | Sanchéz-Martin et al, 2021 |
| ***Pm4b*_mutant_151** | EMS mutant (P184L) in *Fed-Pm4b* NIL background | S | S | S | S | Sanchéz-Martin et al, 2021 |
| ***Pm4b*_mutant_207** | EMS mutant (D170N) in *Fed-Pm4b* NIL background | S | S | S | S | Sanchéz-Martin et al, 2021 |
| ***Pm4b*_mutant_495_1** | EMS mutant (Q274X) in *Fed-Pm4b* NIL background | S | S | S | S | Sanchéz-Martin et al, 2022 |
| ***Pm4b*_mutant_495_3** | EMS mutant (Q274X) in *Fed-Pm4b* NIL background | S | S | S | S | Sanchéz-Martin et al, 2021 |
| ***Pm4b*_mutant_526** | EMS mutant (R291K) in *Fed-Pm4b* NIL background | S | S | S | S | Sanchéz-Martin et al, 2021 |
| ***Pm4b*_mutant_532** | EMS mutant (G104E) in *Fed-Pm4b* NIL background | S | S | S | S | Sanchéz-Martin et al, 2021 |
| ***Pm4b*_mutant_641** | EMS mutant (G45E) in *Fed-Pm4b* NIL background | S | S | S | S | Sanchéz-Martin et al, 2021 |
| ***Pm4b*_Nr#3** | *Pm4b* overexpression line | R | R | S | R | Sanchéz-Martin et al, 2021 |
| ***Pm4b*_S#3** | Susceptible sister line to *Pm4b*_Nr#3 | S | S | S | S | Sanchéz-Martin et al, 2021 |
| ***Pm4b*_Nr#52** | *Pm4b* overexpression line | R | R | S | R | Sanchéz-Martin et al, 2021 |
| ***Pm4b*_S#52** | Susceptible sister line to *Pm4b*_Nr#52 | S | S | S | S | Sanchéz-Martin et al, 2021 |

**Table S12 -** Primers used within this study.

| Name | Sequence (5' to 3') | Description | Function | Reference |
|---|---|---|---|---|
| GH414 | TAGGTTGGAGAGATCACAACGA | F; Exon 5-6; 179 bp | qRT-PCR *Pm4*_V1 expression | Sanchéz-Martin et al, 2021 |
| GH415 | CTGAGGTAGAGGAGGCAACTT | R; Exon 5-6; 179 bp | qRT-PCR *Pm4*_V1 expression | Sanchéz-Martin et al, 2021 |
| GH377 | AGAGTGCAGAGACTTCAATCCA | F; Exon 5-7; 159 bp | qRT-PCR *Pm4*_V2 expression | Sanchéz-Martin et al, 2021 |
| GH417 | TTCTTCGTACCCAGCAGGTC | R; Exon 5-7; 159 bp | qRT-PCR *Pm4*_V2 expression | Sanchéz-Martin et al, 2021 |
| GH094 | TCTCATGGTTGGTCTCGATG | F; Exon 2; 80 bp | qRT-PCR reference gene ADP | Giménez et al, as referenced by Sanchéz-Martin et al, 2021 |
| GH095 | GGATGGTGGTGACGATCTCT | R; Exon 2; 80 bp | qRT-PCR reference gene ADP | Giménez et al, as referenced by Sanchéz-Martin et al, 2021 |
| GH105 | CAGGCATCTCACTGGAGACT | F; Exon 1; 79 bp | qRT-PCR reference gene ZFL | Sanchéz-Martin et al, 2021 |
| GH106 | TGGCATCTCTCTTGCTTCTG | R; Exon 1; 79 bp | qRT-PCR reference gene ZFL | Sanchéz-Martin et al, 2021 |
| P1_F_hex | gaaggtcggagtcaacggatCAAGGCCAACTTCTACCGCT | F | *Pm4* presence/absence KASP (*Pm4*) | This study |
| P1_F_fam | gaaggtgaccaagttcatgctAAGGCCAACTTCTACCGCA | F | *Pm4* presence/absence KASP (*pm4*) | This study |
| P1_COM | ACTTGCAGATGCCGTCGA | R | *Pm4* presence/absence KASP | This study |

**Table S13** – Marker based prediction of the presence of *Pm4* within the Gediflux collection of North western European wheat cultivars. Cultivar information was proved by Luzie Wingen, JIC. Cultivars were genotyped using the '*Pm4* presence absence' KASP primer set (Table S12). In the '*Pm4* Genotype' column: 'Y:Y' indicates the cultivar contains an allele of *Pm4*; 'X:X' indicates the cultivar does not contain an allele of Pm4; '-' indicates insufficient/no DNA for genotyping. In the 'Country of origin' column, 'NL2000' indicates the accession was on the 'Great Britain national list 2000.

| Cultivar | Country of origin | Release decade | *Pm4* Genotype |
|---|---|---|---|
| Alba | Netherlands | 1940 | X:X |
| Holdfast | Great Britain | 1940 | X:X |
| Juliana | Great Britain | 1940 | X:X |
| Lovenik | Netherlands | 1940 | X:X |
| Mendel | Netherlands | 1940 | X:X |
| Redman | Great Britain | 1940 | X:X |
| Stedfast | Great Britain | 1940 | X:X |
| Blanco | Sweden | 1950 | - |
| Carstens 8 | East Germany | 1950 | X:X |
| Dr Lassers Dickkopf | Austria | 1950 | X:X |
| Loosdorfer Austro Bankut Grannen | Austria | 1950 | X:X |
| Minster | Great Britain | 1950 | X:X |
| Peragis | East Germany | 1950 | X:X |
| Pilot (GB) | Great Britain | 1950 | X:X |
| Stam 101 | Austria | 1950 | X:X |
| Strubes Dickkope | East Germany | 1950 | - |

| | | | |
|---|---|---|---|
| Svalov Kronen | East Germany | 1950 | X:X |
| Tassilo | Austria | 1950 | X:X |
| Admonter | Austria | 1960 | X:X |
| Apollo (NL) | Netherlands | 1960 | X:X |
| Cleo | Netherlands | 1960 | X:X |
| Dram Hofner Kolben | Austria | 1960 | X:X |
| Elite Leupeuple | Great Britain | 1960 | X:X |
| Felix | Netherlands | 1960 | - |
| Flevina | Netherlands | 1960 | X:X |
| Florian | East Germany | 1960 | - |
| Hubertusweisen | Austria | 1960 | X:X |
| Hybrid 46 | Great Britain | 1960 | X:X |
| Ibis | Netherlands | 1960 | Y:Y |
| Norda | Belgium | 1960 | X:X |
| Pontus | Austria | 1960 | X:X |
| Probus | Austria | 1960 | X:X |
| Professor Marchal | Great Britain | 1960 | X:X |
| Rabe | East Germany | 1960 | - |
| Record | Austria | 1960 | - |
| Schweigers Taca | Austria | 1960 | X:X |
| Stella | Netherlands | 1960 | - |
| Thor | Great Britain | 1960 | X:X |
| Triumph | Austria | 1960 | X:X |
| Almus | West Germany | 1970 | - |
| Atou | Great Britain | 1970 | X:X |
| Benno | East Germany | 1970 | X:X |
| Bouquet | Great Britain | 1970 | X:X |
| Cama | Great Britain | 1970 | - |

| | | | |
|---|---|---|---|
| Clément | France | 1970 | - |
| Courtot | France | 1970 | Y:Y |
| Cyrano | Netherlands | 1970 | X:X |
| Danubius | Austria | 1970 | X:X |
| Diplomat | East Germany | 1970 | - |
| Extrem | Austria | 1970 | X:X |
| Fakir | West Germany | 1970 | Y:Y |
| Fanal | West Germany | 1970 | X:X |
| Flinor | Great Britain | 1970 | X:X |
| Hardi | France | 1970 | - |
| Kador | Great Britain | 1970 | X:X |
| Kawkas | West Germany | 1970 | X:X |
| Kinsman | Great Britain | 1970 | X:X |
| Kormoran | East Germany | 1970 | - |
| Kranich | Sweden | 1970 | X:X |
| Lely | Netherlands | 1970 | X:X |
| Maris Freeman | Great Britain | 1970 | X:X |
| Maris Nimrod | Great Britain | 1970 | X:X |
| Maris Ranger | Great Britain | 1970 | X:X |
| Mega | Great Britain | 1970 | X:X |
| Mironowskaja | West Germany | 1970 | X:X |
| Multiweiss | Austria | 1970 | X:X |
| Poros | West Germany | 1970 | X:X |
| Solid | Sweden | 1970 | - |
| Sportsman | Great Britain | 1970 | Y:Y |
| Starke 2 | Sweden | 1970 | X:X |
| Top | France | 1970 | X:X |
| Walde | Sweden | 1970 | X:X |

| | | | |
|---|---|---|---|
| Winneton | West Germany | 1970 | X:X |
| Agron | Austria | 1980 | X:X |
| Albatross | Belgium | 1980 | X:X |
| Anja | Denmark | 1980 | X:X |
| Arkos | West Germany | 1980 | X:X |
| Avalon | Great Britain | 1980 | X:X |
| Beauchamp | France | 1980 | X:X |
| Bounty | Great Britain | 1980 | X:X |
| Brigand | Great Britain | 1980 | X:X |
| Brimstone | Great Britain | 1980 | - |
| Brock | Great Britain | 1980 | X:X |
| Calif | Germany | 1980 | X:X |
| Camp Rémy | France | 1980 | X:X |
| Capitole | Belgium | 1980 | - |
| Compal | West Germany | 1980 | Y:Y |
| David | Austria | 1980 | - |
| Fenman | Great Britain | 1980 | X:X |
| Festival | France | 1980 | X:X |
| Fidel | France | 1980 | X:X |
| Folke | Sweden | 1980 | X:X |
| Fontus | Austria | 1980 | - |
| Gamin | Belgium | 1980 | X:X |
| Granada | Netherlands | 1980 | X:X |
| Granta | Netherlands | 1980 | X:X |
| Helge | Sweden | 1980 | X:X |
| Hubertus | Austria | 1980 | - |
| Hustler | Great Britain | 1980 | X:X |
| Iena | Belgium | 1980 | X:X |

| | | | |
|---|---|---|---|
| Karat | Austria | 1980 | X:X |
| Longbow | Great Britain | 1980 | - |
| Martin | Austria | 1980 | X:X |
| Mission | Great Britain | 1980 | - |
| Moulin | France | 1980 | X:X |
| Norman | Great Britain | 1980 | - |
| Odeon | Belgium | 1980 | X:X |
| Perlo | Austria | 1980 | - |
| Pernel | France | 1980 | Y:Y |
| Rapier | Great Britain | 1980 | X:X |
| Regent | Austria | 1980 | X:X |
| Scipion | France | 1980 | X:X |
| Slejpner | Sweden | 1980 | X:X |
| Sperber | East Germany | 1980 | X:X |
| Stetson | Great Britain | 1980 | X:X |
| Taras | West Germany | 1980 | X:X |
| Titus | Austria | 1980 | X:X |
| Virtue | Great Britain | 1980 | X:X |
| Zemon | Belgium | 1980 | X:X |
| Abbot | Great Britain | 1990 | X:X |
| Altria | France | 1990 | - |
| Aztec | France | 1990 | X:X |
| Borenos | West Germany | 1990 | X:X |
| Brigadier | Great Britain | 1990 | Y:Y |
| Buchan | Great Britain | 1990 | Y:Y |
| Buzzard | Germany | 1990 | X:X |
| Cadenza | Great Britain | 1990 | X:X |
| Capo | Austria | 1990 | X:X |

| | | | |
|---|---|---|---|
| Cezanne | France | 1990 | X:X |
| Charger | France | 1990 | X:X |
| Claudius | Austria | 1990 | X:X |
| Contra | Germany | 1990 | Y:Y |
| Equinox | Great Britain | 1990 | X:X |
| Expert | Austria | 1990 | X:X |
| Faktor | Germany | 1990 | Y:Y |
| Flair | Germany | 1990 | X:X |
| Flame | Great Britain | 1990 | Y:Y |
| Florida | Denmark | 1990 | - |
| Genesis | Great Britain | 1990 | X:X |
| Georg | Austria | 1990 | X:X |
| Grief | Germany | 1990 | X:X |
| Haven | Great Britain | 1990 | X:X |
| Hunter | Great Britain | 1990 | X:X |
| Ibis (D) | Germany | 1990 | X:X |
| Ikarus | Austria | 1990 | - |
| Isengrain | France | 1990 | X:X |
| Kontrast | Germany | 1990 | X:X |
| Lindos | Austria | 1990 | Y:Y |
| Mikon | Germany | 1990 | X:X |
| Napier | Great Britain | 1990 | X:X |
| Obelisk | Germany | 1990 | X:X |
| Optimus | Austria | 1990 | X:X |
| Pallus | Germany | 1990 | X:X |
| Pastiche | Great Britain | 1990 | X:X |
| Pegassos | Germany | 1990 | X:X |
| Pepital | Denmark | 1990 | X:X |

| | | | |
|---|---|---|---|
| Renan | Austria | 1990 | Y:Y |
| Riband | Great Britain | 1990 | Y:Y |
| Ritmo | Germany | 1990 | X:X |
| Savannah | Great Britain | 1990 | Y:Y |
| Shamrock | Great Britain | 1990 | X:X |
| Sideral | France | 1990 | - |
| Silvius | Austria | 1990 | - |
| Spark | Great Britain | 1990 | Y:Y |
| Tambor | Austria | 1990 | X:X |
| Torfrida | Great Britain | 1990 | Y:Y |
| Toronto | Germany | 1990 | Y:Y |
| Trémie | France | 1990 | X:X |
| Zentra | Germany | 1990 | X:X |
| Ability | Great Britain (NL2000) | 2000 | Y:Y |
| Access | Great Britain (NL2000) | 2000 | Y:Y |
| Admiral | Great Britain (NL2000) | 2000 | X:X |
| Adroit | Great Britain (NL2000) | 2000 | Y:Y |
| Alcier | Great Britain (NL2000) | 2000 | Y:Y |
| Alert | Great Britain (NL2000) | 2000 | X:X |
| Ambassador | Great Britain (NL2000) | 2000 | X:X |
| Andante | Great Britain (NL2000) | 2000 | X:X |
| Anthem | Great Britain (NL2000) | 2000 | X:X |
| Apostle | Great Britain (NL2000) | 2000 | X:X |
| Aristocrat | Great Britain (NL2000) | 2000 | X:X |
| Assett | Great Britain (NL2000) | 2000 | - |
| Athlet | Great Britain (NL2000) | 2000 | X:X |
| Atla | Great Britain (NL2000) | 2000 | X:X |
| Attoll | Great Britain (NL2000) | 2000 | X:X |

| | | | |
|---|---|---|---|
| Avocet | Great Britain (NL2000) | 2000 | X:X |
| Axial | Great Britain (NL2000) | 2000 | X:X |
| Bandit | Great Britain (NL2000) | 2000 | X:X |
| Banner | Great Britain (NL2000) | 2000 | X:X |
| Baron | Great Britain (NL2000) | 2000 | X:X |
| Beaufort | Great Britain (NL2000) | 2000 | X:X |
| Belplaine | Great Britain (NL2000) | 2000 | X:X |
| Bercy | Great Britain (NL2000) | 2000 | X:X |
| Bert | Great Britain (NL2000) | 2000 | X:X |
| Biscay | Great Britain (NL2000) | 2000 | Y:Y |
| Blitz | Great Britain (NL2000) | 2000 | X:X |
| Bourbon | Great Britain (NL2000) | 2000 | X:X |
| Boxer | Great Britain (NL2000) | 2000 | Y:Y |
| Breval | Great Britain (NL2000) | 2000 | X:X |
| Brutus | Great Britain (NL2000) | 2000 | X:X |
| Bryden | Great Britain (NL2000) | 2000 | Y:Y |
| Bullet | Great Britain (NL2000) | 2000 | X:X |
| Buster | Great Britain (NL2000) | 2000 | - |
| Cambat | Great Britain (NL2000) | 2000 | - |
| Caprimus | Great Britain (NL2000) | 2000 | X:X |
| Captor | Great Britain (NL2000) | 2000 | X:X |
| Carolus | Great Britain (NL2000) | 2000 | X:X |
| Catamaran | Great Britain (NL2000) | 2000 | X:X |
| Caxton | Great Britain (NL2000) | 2000 | X:X |
| Cheetah | Great Britain (NL2000) | 2000 | X:X |
| Chianti | Great Britain (NL2000) | 2000 | Y:Y |
| Civic | Great Britain (NL2000) | 2000 | X:X |
| Claire | Great Britain (NL2000) | 2000 | Y:Y |

| | | | |
|---|---|---|---|
| Clove | Great Britain (NL2000) | 2000 | X:X |
| Club | Great Britain (NL2000) | 2000 | X:X |
| Cobalt | Great Britain (NL2000) | 2000 | X:X |
| Commodore | Great Britain (NL2000) | 2000 | X:X |
| Consort | Great Britain (NL2000) | 2000 | Y:Y |
| Contour | Great Britain (NL2000) | 2000 | X:X |
| Corinthian | Great Britain (NL2000) | 2000 | X:X |
| Corsaire | Great Britain (NL2000) | 2000 | X:X |
| Coxwain | Great Britain (NL2000) | 2000 | X:X |
| Creneau | Great Britain (NL2000) | 2000 | X:X |
| Crest | Great Britain (NL2000) | 2000 | Y:Y |
| Daphne | Great Britain (NL2000) | 2000 | X:X |
| Dean | Great Britain (NL2000) | 2000 | X:X |
| Deben | Great Britain (NL2000) | 2000 | Y:Y |
| Denver | Great Britain (NL2000) | 2000 | X:X |
| Depot | Great Britain (NL2000) | 2000 | X:X |
| Destroyer | Great Britain (NL2000) | 2000 | Y:Y |
| Diablo | Great Britain (NL2000) | 2000 | - |
| Dorby | Great Britain (NL2000) | 2000 | - |
| Drake | Great Britain (NL2000) | 2000 | X:X |
| Druid | Great Britain (NL2000) | 2000 | X:X |
| Dynamo | Great Britain (NL2000) | 2000 | X:X |
| Eagle | Great Britain (NL2000) | 2000 | - |
| Emblem | Great Britain (NL2000) | 2000 | Y:Y |
| Encore | Great Britain (NL2000) | 2000 | Y:Y |
| Erland | Great Britain (NL2000) | 2000 | X:X |
| Estica | Great Britain (NL2000) | 2000 | X:X |
| Estorial | Great Britain (NL2000) | 2000 | X:X |

| | | | |
|---|---|---|---|
| Fenda | Great Britain (NL2000) | 2000 | X:X |
| Feuvert | Great Britain (NL2000) | 2000 | X:X |
| Flash | Great Britain (NL2000) | 2000 | Y:Y |
| Fletum | Great Britain (NL2000) | 2000 | - |
| Focus | Great Britain (NL2000) | 2000 | X:X |
| Foreman | Great Britain (NL2000) | 2000 | X:X |
| Fortress | Great Britain (NL2000) | 2000 | X:X |
| Fresco | Great Britain (NL2000) | 2000 | X:X |
| Frista | Great Britain (NL2000) | 2000 | X:X |
| Fromendor | Great Britain (NL2000) | 2000 | X:X |
| Gallatea | Great Britain (NL2000) | 2000 | X:X |
| Galliard | Great Britain (NL2000) | 2000 | X:X |
| Gambit | Great Britain (NL2000) | 2000 | X:X |
| Gondola | Great Britain (NL2000) | 2000 | X:X |
| Governor | Great Britain (NL2000) | 2000 | X:X |
| Guardian | Great Britain (NL2000) | 2000 | X:X |
| Hanno | Great Britain (NL2000) | 2000 | X:X |
| Harrier | Great Britain (NL2000) | 2000 | Y:Y |
| Heinrich | Great Britain (NL2000) | 2000 | - |
| Hickory | Great Britain (NL2000) | 2000 | Y:Y |
| Holster | Great Britain (NL2000) | 2000 | Y:Y |
| Hudson | Great Britain (NL2000) | 2000 | Y:Y |
| Hussar | Great Britain (NL2000) | 2000 | - |
| Imola | Great Britain (NL2000) | 2000 | X:X |
| Jubilatka | Great Britain (NL2000) | 2000 | X:X |
| Kontiki | Great Britain (NL2000) | 2000 | X:X |
| Kronjewel | Great Britain (NL2000) | 2000 | Y:Y |
| Kyalami | Great Britain (NL2000) | 2000 | X:X |

| | | | |
|---|---|---|---|
| Lancelot | Great Britain (NL2000) | 2000 | X:X |
| Legend | Great Britain (NL2000) | 2000 | Y:Y |
| Leo | Great Britain (NL2000) | 2000 | X:X |
| Lynx | Great Britain (NL2000) | 2000 | X:X |
| Madrigal | Great Britain (NL2000) | 2000 | X:X |
| Magellan | Great Britain (NL2000) | 2000 | Y:Y |
| Malacca | Great Britain (NL2000) | 2000 | Y:Y |
| Mandate | Great Britain (NL2000) | 2000 | X:X |
| Mantle | Great Britain (NL2000) | 2000 | X:X |
| Mars | Great Britain (NL2000) | 2000 | X:X |
| Meteor | Great Britain (NL2000) | 2000 | X:X |
| Morell | Great Britain (NL2000) | 2000 | Y:Y |
| Motto | Great Britain (NL2000) | 2000 | X:X |
| Newhaven | Great Britain (NL2000) | 2000 | X:X |
| Norsman | Great Britain (NL2000) | 2000 | X:X |
| Option | Great Britain (NL2000) | 2000 | X:X |
| Orqual | Great Britain (NL2000) | 2000 | X:X |
| Ostara | Great Britain (NL2000) | 2000 | X:X |
| Parade | Great Britain (NL2000) | 2000 | X:X |
| Patience | Great Britain (NL2000) | 2000 | X:X |
| Peacock | Great Britain (NL2000) | 2000 | Y:Y |
| Piccadilly | Great Britain (NL2000) | 2000 | X:X |
| Pistol | Great Britain (NL2000) | 2000 | X:X |
| Poet | Great Britain (NL2000) | 2000 | X:X |
| Profet | Great Britain (NL2000) | 2000 | X:X |
| Profi | Great Britain (NL2000) | 2000 | Y:Y |
| Prospect | Great Britain (NL2000) | 2000 | X:X |
| Puma | Great Britain (NL2000) | 2000 | Y:Y |

| | | | |
|---|---|---|---|
| Raleigh | Great Britain (NL2000) | 2000 | X:X |
| Reaper | Great Britain (NL2000) | 2000 | X:X |
| Rebel | Great Britain (NL2000) | 2000 | - |
| Renard | Great Britain (NL2000) | 2000 | X:X |
| Rendezvous | Great Britain (NL2000) | 2000 | Y:Y |
| Renown | Great Britain (NL2000) | 2000 | X:X |
| Rhino | Great Britain (NL2000) | 2000 | Y:Y |
| Rialto | Great Britain (NL2000) | 2000 | X:X |
| Rifle | Great Britain (NL2000) | 2000 | X:X |
| Ritz | Great Britain (NL2000) | 2000 | X:X |
| Rocket | Great Britain (NL2000) | 2000 | X:X |
| Rooster | Great Britain (NL2000) | 2000 | Y:Y |
| Rostrum | Great Britain (NL2000) | 2000 | X:X |
| Rubens | Great Britain (NL2000) | 2000 | X:X |
| Russett | Great Britain (NL2000) | 2000 | Y:Y |
| Sabre | Great Britain (NL2000) | 2000 | X:X |
| Samson | Great Britain (NL2000) | 2000 | X:X |
| Sarek | Great Britain (NL2000) | 2000 | X:X |
| Sarsen | Great Britain (NL2000) | 2000 | Y:Y |
| Saxon | Great Britain (NL2000) | 2000 | X:X |
| Semper | Great Britain (NL2000) | 2000 | X:X |
| Sennet | Great Britain (NL2000) | 2000 | X:X |
| Shango | Great Britain (NL2000) | 2000 | Y:Y |
| Shannon | Great Britain (NL2000) | 2000 | X:X |
| Sickle | Great Britain (NL2000) | 2000 | Y:Y |
| Sirius | Great Britain (NL2000) | 2000 | X:X |
| Sitka | Great Britain (NL2000) | 2000 | X:X |
| Sniper | Great Britain (NL2000) | 2000 | X:X |

| | | | |
|---|---|---|---|
| Soleil | Great Britain (NL2000) | 2000 | X:X |
| Solstice | Great Britain (NL2000) | 2000 | X:X |
| Spice | Great Britain (NL2000) | 2000 | X:X |
| Spitfire | Great Britain (NL2000) | 2000 | Y:Y |
| Spray | Great Britain (NL2000) | 2000 | X:X |
| Squadron | Great Britain (NL2000) | 2000 | Y:Y |
| Stag | Great Britain (NL2000) | 2000 | X:X |
| Stallion | Great Britain (NL2000) | 2000 | X:X |
| Tallon | Great Britain (NL2000) | 2000 | X:X |
| Tandero | Great Britain (NL2000) | 2000 | X:X |
| Tanker | Great Britain (NL2000) | 2000 | X:X |
| Tara | Great Britain (NL2000) | 2000 | X:X |
| Tessa | Great Britain (NL2000) | 2000 | X:X |
| Texel | Great Britain (NL2000) | 2000 | X:X |
| Thunder | Great Britain (NL2000) | 2000 | X:X |
| Tilburi | Great Britain (NL2000) | 2000 | X:X |
| Tjalk | Great Britain (NL2000) | 2000 | X:X |
| Token | Great Britain (NL2000) | 2000 | X:X |
| Tomo | Great Britain (NL2000) | 2000 | Y:Y |
| Torch | Great Britain (NL2000) | 2000 | X:X |
| Toucan | Great Britain (NL2000) | 2000 | Y:Y |
| Trader | Great Britain (NL2000) | 2000 | X:X |
| Trafalgar | Great Britain (NL2000) | 2000 | X:X |
| Trawler | Great Britain (NL2000) | 2000 | X:X |
| Trend | Great Britain (NL2000) | 2000 | Y:Y |
| Turpin | Great Britain (NL2000) | 2000 | Y:Y |
| Vauntless | Great Britain (NL2000) | 2000 | - |
| Veritas | Great Britain (NL2000) | 2000 | X:X |

| | | | |
|---|---|---|---|
| Victo | Great Britain (NL2000) | 2000 | - |
| Vivant | Great Britain (NL2000) | 2000 | X:X |
| Vocal | Great Britain (NL2000) | 2000 | X:X |
| Voyage | Great Britain (NL2000) | 2000 | X:X |
| Warrior | Great Britain (NL2000) | 2000 | - |
| Wasp | Great Britain (NL2000) | 2000 | Y:Y |
| Welton | Great Britain (NL2000) | 2000 | X:X |
| Wizard | Great Britain (NL2000) | 2000 | X:X |
| Woodstock | Great Britain (NL2000) | 2000 | X:X |
| Wykenham | Great Britain (NL2000) | 2000 | Y:Y |
| Xi 19 | Great Britain (NL2000) | 2000 | X:X |
| Yacht | Great Britain (NL2000) | 2000 | X:X |
| Zodiac | Great Britain (NL2000) | 2000 | X:X |
| Banco | - | - | X:X |
| Bledor | Belgium | - | X:X |
| Capitaine | - | - | X:X |
| Capitole | - | - | X:X |
| Caribo | - | - | X:X |
| Combat | - | - | X:X |
| Dauntless | - | - | Y:Y |
| Ergo | Sweden | - | X:X |
| Eroica | Sweden | - | X:X |
| Eroica II | Sweden | - | X:X |
| Ertus | Sweden | - | X:X |
| Jarl | Sweden | - | X:X |
| Meredien | Sweden | - | - |
| Odin | Sweden | - | - |
| Probstdorfer Stabil | - | - | X:X |

| | | | |
|---|---|---|---|
| Roi Albert | Belgium | - | X:X |
| Rufus | Belgium | - | X:X |
| Schreibers Sturmweizen | - | - | X:X |
| Skandia | Sweden | - | X:X |
| Stava | Sweden | - | Y:Y |
| Svale | Sweden | - | X:X |
| Svalov 0987 | - | - | X:X |
| Terra | Sweden | - | X:X |
| Virgo | Sweden | - | X:X |
| Virtus | Sweden | - | X:X |
| William | Sweden | - | Y:Y |
| Flair Eigeno Nachbau | Germany? | ? | X:X |
| Glicevka | Germany? | ? | - |
| Bersee | Great Britain | 1940-50 | X:X |
| Criewener 152 | East Germany | 1940-50 | X:X |
| Ebersbacher Weiss | East Germany | 1940-50 | X:X |
| Heine 4 | East Germany | 1940-50 | X:X |
| Kadolzer | Austria | 1940-50 | X:X |
| Mahndorfer Tempo | East Germany | 1940-50 | X:X |
| Rimpaus Bastard 2 | East Germany | 1940-50 | - |
| Rimpaus Bastard 2 | - | 1940-50 | - |
| Rimpaus Braun | East Germany | 1940-50 | - |
| Ritzlhofer Neu | Austria | 1940-50 | X:X |
| Salzmunder Standard | East Germany | 1940-50 | X:X |
| Starling | Great Britain | 1940-50 | X:X |
| Svalon 0907 | East Germany | 1940-50 | - |
| TscherWaks Bergrannter Machfeld | Austria | 1940-50 | X:X |
| Vague d'épis | France | 1940-50 | X:X |

| | | | |
|---|---|---|---|
| Vilmorlin 27 | France | 1940-50 | X:X |
| Warden | Great Britain | 1940-50 | X:X |
| Cappelle Desprez | France | 1950-60 | X:X |
| Carstens | | 1950-60 | X:X |
| Carstens | Netherlands | 1950-60 | X:X |
| Eros | West Germany | 1950-60 | X:X |
| Etoile de Choisy | France | 1950-60 | X:X |
| Flamingo | Great Britain | 1950-60 | X:X |
| Heine 7 | East Germany | 1950-60 | X:X |
| Hochland | West Germany | 1950-60 | X:X |
| Leda | Netherlands | 1950-60 | X:X |
| Muck | West Germany | 1950-60 | X:X |
| Triumph (NC) | Netherlands | 1950-60 | X:X |
| Vilmorin 53 | France | 1950-60 | X:X |
| Werla | East Germany | 1950-60 | X:X |
| Capitole | France | 1960-70 | - |
| Champlein | France | 1960-70 | X:X |
| Erla Kolben | Austria | 1960-70 | X:X |
| Joss Cambier | France | 1960-70 | X:X |
| Jubilar | East Germany | 1960-70 | X:X |
| Manella | Netherlands | 1960-70 | X:X |
| Maris Widgeon | Great Britain | 1960-70 | X:X |
| Moisson | France | 1960-70 | X:X |
| Orlando | West Germany | 1960-70 | X:X |
| Remois | France | 1960-70 | X:X |
| Starke | Sweden | 1960-70 | X:X |
| Tadorna | Netherlands | 1960-70 | X:X |
| Adam | Austria | 1970-80 | X:X |

| | | | |
|---|---|---|---|
| Alcedo | West Germany | 1970-80 | X:X |
| Aquila | Great Britain | 1970-80 | X:X |
| Armada | Great Britain | 1970-80 | Y:Y |
| Arminda | France | 1970-80 | X:X |
| Capiso | - | 1970-80 | - |
| Disponent | East Germany | 1970-80 | X:X |
| Flanders | Great Britain | 1970-80 | X:X |
| Hildur | Sweden | 1970-80 | X:X |
| Hobbit | Great Britain | 1970-80 | X:X |
| Holme | Sweden | 1970-80 | X:X |
| Lutin | France | 1970-80 | X:X |
| Mardler | Great Britain | 1970-80 | X:X |
| Maris Huntsman | Great Britain | 1970-80 | X:X |
| Micronowskata | West Germany | 1970-80 | X:X |
| Nautica | Netherlands | 1970-80 | X:X |
| Oenus | Austria | 1970-80 | X:X |
| Okapi | East Germany | 1970-80 | - |
| Talent | France | 1970-80 | X:X |
| Vuka | East Germany | 1970-80 | X:X |
| Apollo (D) | East Germany | 1980-90 | X:X |
| Beaver | Great Britain | 1980-90 | X:X |
| Escoria | Belgium | 1980-90 | X:X |
| Galahad | Great Britain | 1980-90 | X:X |
| Gawain | Denmark | 1980-90 | X:X |
| Hereward | Great Britain | 1980-90 | X:X |
| Hornet | Great Britain | 1980-90 | X:X |
| Kanzler | East Germany | 1980-90 | X:X |
| Kosak | Sweden | 1980-90 | Y:Y |

| | | | |
|---|---|---|---|
| Mercia | Great Britain | 1980-90 | X:X |
| Miras | West Germany | 1980-90 | X:X |
| Recital | France | 1980-90 | - |
| Regina | West Germany | 1980-90 | X:X |
| Rektor | East Germany | 1980-90 | X:X |
| Soissons | France | 1980-90 | X:X |
| Thesee | France | 1980-90 | - |
| Urban | Denmark | 1980-90 | - |
| Cama | Belgium | - | X:X |
| Celesta | Belgium | - | X:X |
| Chinese Spring | China | - | X:X |
| Clovis | Belgium | - | Y:Y |
| Hesbinion | Belgium | - | X:X |
| Jason | Belgium | - | X:X |
| Marco | Belgium | - | X:X |
| Marisa | Belgium | - | X:X |
| Mina | Belgium | - | X:X |
| Mutant Odeon | Belgium | - | X:X |
| Mutant Odeon II | Belgium | - | X:X |
| Orestis | Germany | - | X:X |
| Pony | Belgium | - | X:X |
| Prima | Belgium | - | X:X |
| Stella | Belgium | - | - |

**Table S14** - Watkins accessions and adapted wheat varieties tested within this study that carry *Pm4.* Scores were taken from detached leaves at six-days post inoculation.

| Line | *Pm4* allele | NO6047+*AVR8* (6 dpi) | Py 15.1.018 (6 dpi) |
|---|---|---|---|
| Ability | *Pm4b* | 2.33 | 1.00 |
| Claire | | 1.20 | 1.20 |
| Flame | | 2.33 | 1.00 |
| Malacca | | 3.33 | 2.00 |
| Revelation | | 1.40 | 1.20 |
| Riband | | 3.00 | 1.67 |
| Shango | | 2.33 | 1.67 |
| Spark | | 0.67 | 1.33 |
| Wasp | | 2.00 | 1.00 |
| WATDE0102 | | 2.60 | 2.20 |
| WATDE0171 | | 1.80 | 0.80 |
| WATDE0310 | | 1.20 | 0.80 |
| SY-Epson* | *Pm4d* | 1.20 | 0.00 |
| Renan* | | 2.00 | 0.80 |
| SY-Mattis* | | 1.80 | 1.20 |
| CDC Stanley* | | 2.60 | 1.20 |
| WATDE0369 | *Pm4f* | 3.00 | 1.30 |
| WATDE0426 | | | 1.00 |
| WATDE0427 | | 1.00 | 0.00 |
| WATDE0428 | | | 1.00 |

| | | | |
|---|---|---|---|
| WATDE0465 | | | 1.20 |
| WATDE0477 | | 0.00 | 0.00 |
| WATDE0505 | | | 1.20 |
| WATDE0526 | | 1.00 | 0.67 |
| WATDE0541 | | | 0.80 |
| WATDE0546 | | | 1.20 |
| WATDE0566 | | 1.60 | 2.20 |
| WATDE0567 | | | 1.20 |
| WATDE0571 | | 1.80 | 1.00 |
| WATDE0672 | | | 1.20 |
| WATDE0687 | | | 0.80 |
| WATDE0804 | | 0.80 | 1.80 |
| WATDE0970 | | | 1.00 |
| WATDE0971 | | 0.67 | 0.00 |
| WATDE0973 | | 0.33 | 0.30 |
| WATDE0974 | | | 1.00 |
| WATDE1062 | | | 1.20 |
| WATDE0048 | *Pm4i* | 4.60 | 5.60 |
| WATDE0527 | | 4.60 | 5.80 |
| WATDE0568 | *Pm4j* | 5.60 | 5.80 |
| WATDE0592 | | 4.60 | 5.60 |

* contains the *Ae. ventricosa* 2NS translocation

**Table S15** - BR48 isogenic series detached leaf and spike assay scores. () indicate which *Pm4* allele the line contains. Scores were taken at five or six days post inoculation.

| Accession (*Pm4* allele) | Mean detached leaf assay score | | | | Detached spike assay | |
|---|---|---|---|---|---|---|
| | **BR48ΔeI** | **BR48ΔeI+eI** | **BR48ΔeI+eII** | **BR48ΔeI+eII'** | **BR48ΔeI** | **BR48ΔeI+eI** |
| Bobwhite | 4.00 | 4.00 | 4.25 | 5.75 | 6.00 | 6.00 |
| Federation | 5.75 | 5.50 | 5.50 | 6.00 | 6.00 | 6.00 |
| *Fed-Pm4a*-NIL | 5.75 | 1.75 | 0.75 | 1.75 | 6.00 | 5.00 |
| *Fed-Pm4b*-NIL | 5.75 | 1.25 | 1.50 | 2.75 | 6.00 | 4.00 |
| Nr#3 | 3.50 | 1.25 | 1.75 | 1.50 | 6.00 | 5.67 |
| Nr#52 | 4.00 | 1.25 | 1.00 | 2.25 | 5.67 | 2.00 |
| S3 | 3.75 | 4.00 | 4.25 | 5.00 | 6.00 | 6.00 |
| S52 | 3.50 | 3.75 | 3.75 | 3.50 | 6.00 | 6.00 |
| *Pm4b*_Mut_207 | 5.50 | 6.00 | 5.50 | 6.00 | 6.00 | 6.00 |
| *Pm4b*_Mut_495-3 | 5.50 | 5.50 | 6.00 | 6.00 | 6.00 | 6.00 |
| *Pm4b*_Mut_526 | 4.25 | 4.25 | 4.00 | 4.25 | 6.00 | 6.00 |
| *Pm4b*_Mut_641 | 5.50 | 3.50 | 4.75 | 4.50 | 6.00 | 6.00 |
| WATDE0102(*Pm4b*) | 1.50 | 0.50 | 0.00 | 0.75 | - | - |
| WATDE0310(*Pm4b*) | 3.75 | 1.25 | 1.00 | 0.25 | - | - |
| SY-Mattis(*Pm4d*) | 3.75 | 1.25 | 0.00 | 1.00 | - | - |
| CDC Stanley(*Pm4d*) | 3.25 | 1.00 | 0.75 | 1.50 | - | - |
| VPM1(*Pm4d*) | 3.50 | 1.00 | 0.25 | 1.25 | - | - |
| WATDE0427(*Pm4f*) | 3.75 | 1.50 | 1.25 | 1.00 | - | - |
| WATDE0477(*Pm4f*) | 4.50 | 1.25 | 0.25 | 2.00 | - | - |
| WATDE0571(*Pm4f*) | 4.50 | 0.75 | 0.50 | 1.25 | - | - |
| WATDE0048(*Pm4i*) | 2.00 | 1.75 | 1.00 | 2.50 | - | - |

| | | | | | | |
|---|---|---|---|---|---|---|
| WATDE0527(*Pm4i*) | 2.75 | 1.75 | 1.00 | 3.50 | - | - |
| WATDE0568(*Pm4j*) | 4.75 | 4.50 | 4.75 | 5.75 | - | - |
| WATDE0592(*Pm4j*) | 5.50 | 5.25 | 5.00 | 5.75 | - | - |
| WW-093(*Pm4g*) | 2.75 | 3.00 | 3.25 | 4.25 | - | - |
| WW-213(*Pm4g*) | 3.75 | 3.50 | 4.00 | 5.25 | - | - |
| WW-470(*Pm4h*) | 5.00 | 4.50 | 5.50 | 5.75 | - | - |
| WW-474(*Pm4h*) | 3.00 | 1.63 | 1.75 | 2.13 | - | - |

**Table S16** - Target-specific amplification efficiencies of the splicing variants *Pm4b_V1* and *Pm4b_V2* and the reference genes used for RT-qPCR in this study.

| gene / Target | gene ID | position | primer | amplicon length bp | reference | efficiency (%) | Slope | r2 of calibration curve |
|---|---|---|---|---|---|---|---|---|
| Pm4_V1 | | Exon 5-6 | F: TAGGTTGGAGAGATCACAACGA (GH414)<br>R: CTGAGGTAGAGGAGGCAACTT (GH415) | 179 | Sanchéz-Martin et al, 2021 | 100.75 | -3.304 | 0.9983 |
| Pm4_V2 | | Exon 5-7 | F: AGAGTGCAGAGACTTCAATCCA (GH377)<br>R: TTCTTCGTACCCAGCAGGTC (GH417) | 159 | Sanchéz-Martin et al, 2021 | 83.44 | -3.795 | 0.9918 |
| ADP | TraesCS3B01G368600, TraesCS3D01G330500 (TA.2291) | Exon 2 | F: TCTCATGGTTGGTCTCGATG (GH094)<br>R: GGATGGTGGTGACGATCTCT (GH095) | 80 | Giménez et al, as referenced by Sanchéz-Martin et al, 2021 | 98.92 | -3.348 | 0.9985 |
| ZFL | TraesCS3D01G432800, TraesCS3A01G440000 | Exon 1 | F: CAGGCATCTCACTGGAGACT (GH105)<br>R: TGGCATCTCTCTTGCTTCTG (GH106) | 79 | Sanchéz-Martin et al, 2021 | 98.6 | -3.356 | 0.9933 |

**Table S17** - Marker based prediction of the presence of *Pm4* within a selection of 565 CIMMYT wheat lines. Cultivars were genotyped for *Pm4* using the '*Pm4* presence/absence' KASP primer set (Table S12) and the 2NS translocation originating from wheat line VPM1 using marker set CIMwMAS0004 (Helguera et al. 2003). In the 'AVR-Rmg8-R/S' column: 'T:T' indicates the line contains an allele of *Pm4*; 'A:A' indicates the line does not contain an allele of *Pm4*. In the '2NS' column: 'T:T' indicates the line contains the VPM1 translocation; 'C:C' indicates the line does not contain the VPM1 translocation; 'C:T' indicates the line is heterozygous for this marker. The 'CIMwMAS' codes correspond to the CIMMYT marker ID.

| SampleID | Pedigree | Marker set<br>AVR-Rmg8-R/S<br>CIMwMAS1217 | Marker set<br>2NS<br>CIMwMAS0004 |
|---|---|---|---|
| BW22GS018742 | NINGA #1 | A:A | T:T |
| BW22GS018743 | BORL14//BECARD/QUAIU #1 | A:A | T:T |
| BW22GS018744 | MUCUY*2//SUP152/BAJ #1 | A:A | T:T |
| BW22GS018745 | BORL14//SUP152/FRNCLN/3/KASUKO | A:A | T:T |
| BW22GS018746 | BORL14/9/WAXWING/7/TNMU/6/CEP80111/CEP81165/5/IAC5/4/YKT406/3/AG/ASN//ATR/8/ATTILA/3*BCN//BAV 92/3/TILHI/4/SHA7/VEE#5//ARIV92/10/KASUKO | A:A | T:T |
| BW22GS018747 | MUNAL #1/CHIPAK//KASUKO | A:A | T:T |
| BW22GS018748 | AKEPA/BOKOTA/3/BORL14//KFA/2*KACHU | A:A | T:T |
| BW22GS018749 | CHIPAK*2/4/KACHU/3/WHEAR//2*PRL/2*PASTOR | A:A | T:T |
| BW22GS018750 | CHIPAK*2/4/KACHU/3/WHEAR//2*PRL/2*PASTOR | A:A | T:T |
| BW22GS018751 | MUCUY/3/KACHU//KIRITATI/2*TRCH/4/MOKUE #1 | A:A | T:T |
| BW22GS018752 | WHEAR//2*PRL/2*PASTOR/3/QUAIU #1/4/SWSR22T.B.//TACUPETO F2001*2/BRAMBLING/3/2*TACUPETO F2001*2/BRAMBLING/5/BORL14 | A:A | T:T |
| BW22GS018753 | KISKADEE #1/5/KAUZ*2/MNV//KAUZ/3/MILAN/4/BAV92/6/WHEAR//2*PRL/2*PASTOR/7/MUTUS/DANPHE #1/4/C80.1/3*BATAVIA//2*WBLL1/3/C80.1/3*QT4522//2*PASTOR/8/MUNAL #1/CIRO16 | A:A | T:T |
| BW22GS018754 | ISENGRAIN/KBIRD//MUNAL #1/3/SUP152/KENYA SUNBIRD/4/FRNCLN*2/KINGBIRD #1 | A:A | T:T |
| BW22GS018755 | KACHU/SAUAL/3/TACUPETO F2001/BRAMBLING//KIRITATI*2/4/FRET2/TUKURU//FRET2/3/MUNAL #1 | A:A | T:T |
| BW22GS018756 | TRCH/3/ROLF07/YANAC//TACUPETO F2001/BRAMBLING/4/PRL/2*PASTOR/5/BORL14/6/KASUKO | A:A | T:T |
| BW22GS018757 | TRCH/7/TUKURU//BAV92/RAYON/6/NG8201/KAUZ/4/SHA7//PRL/VEE#6/3/FASAN/5/MILAN/KAUZ/8/CIRO16/9/B ORL14/10/MOKUE #1 | A:A | T:T |
| BW22GS018758 | NL971*2/4/HUW234+LR34/PRINIA//INQALAB 91*2/KUKUNA/3/FRET2*2/SHAMA*2/5/MUCUY | A:A | T:T |
| BW22GS018759 | FRANCOLIN #1/3/PBW343*2/KUKUNA*2//YANAC/4/KINGBIRD #1//INQALAB 91*2/TUKURU*2/5/MUNAL #1 | A:A | T:T |

| | | | |
|---|---|---|---|
| BW22GS018760 | FRANCOLIN #1/3/PBW343*2/KUKUNA*2//YANAC/4/KINGBIRD #1//INQALAB 91*2/TUKURU*2/5/KINGBIRD #1//INQALAB 91*2/TUKURU | A:A | T:T |
| BW22GS018761 | FRANCOLIN #1/3/PBW343*2/KUKUNA*2//YANAC/4/KINGBIRD #1//INQALAB 91*2/TUKURU*2/5/KINGBIRD #1//INQALAB 91*2/TUKURU | A:A | T:T |
| BW22GS018762 | FRANCOLIN #1/3/PBW343*2/KUKUNA*2//YANAC/4/KINGBIRD #1//INQALAB 91*2/TUKURU*2/5/CHIPAK | A:A | T:T |
| BW22GS018763 | CAL/NH//H567.71/3/SERI/4/CAL/NH//H567.71/5/2*KAUZ/6/WH576/7/WH 542/8/WAXWING/9/ATTILA*2/PBW65//PIHA/3/ATTILA/2*PASTOR/10/UP2338*2/KKTS*2//YANAC/11/BORL14/12 /KASUKO | A:A | T:T |
| BW22GS018764 | ATTILA*2/PBW65/5/CNO79//PF70354/MUS/3/PASTOR/4/BAV92/6/KINGBIRD #1/7/COPIO*2/8/BORL14 | A:A | T:T |
| BW22GS018765 | KACHU/SUP152//KASUKO/3/KASUKO | A:A | T:T |
| BW22GS018766 | KASUKO*2/4/BECARD/AKURI*2/3/PBW343*2/KUKUNA*2//FRTL/PIFED | A:A | T:T |
| BW22GS018767 | KASUKO*2/CHIPAK | A:A | T:T |
| BW22GS018768 | KASUKO*2/CHIPAK | A:A | T:T |
| BW22GS018769 | KIRITATI//PRL/2*PASTOR/5/OASIS/SKAUZ//4*BCN/3/PASTOR/4/KAUZ*2/YACO//KAUZ/6/KIRITATI//PRL/2*PASTOR /7/PBW343*2/KUKUNA*2//FRTL/PIFED*2/8/KASUKO | A:A | T:T |
| BW22GS018770 | BORL14/MOKUE #1 | A:A | T:T |
| BW22GS018771 | BORL14/7/KFA/2*KACHU/5/WBLL1*2/4/BABAX/LR42//BABAX/3/BABAX/LR42//BABAX/6/KFA/2*KACHU | A:A | T:T |
| BW22GS018772 | BORL14/KASUKO | A:A | T:T |
| BW22GS018773 | BORL14/KASUKO | A:A | T:T |
| BW22GS018774 | BORL14/KASUKO | A:A | T:T |
| BW22GS018775 | BORL14/KASUKO | A:A | T:T |
| BW22GS018776 | BORL14/6/KSW/SAUAL//SAUAL/3/TRCH/HUIRIVIS #1/5/UP2338*2/SHAMA/3/MILAN/KAUZ//CHIL/CHUM18/4/UP2338*2/SHAMA | A:A | T:T |
| BW22GS018777 | BORL14/6/KSW/SAUAL//SAUAL/3/TRCH/HUIRIVIS #1/5/UP2338*2/SHAMA/3/MILAN/KAUZ//CHIL/CHUM18/4/UP2338*2/SHAMA | A:A | T:T |
| BW22GS018778 | BORL14/6/KSW/SAUAL//SAUAL/3/TRCH/HUIRIVIS #1/5/UP2338*2/SHAMA/3/MILAN/KAUZ//CHIL/CHUM18/4/UP2338*2/SHAMA | A:A | T:T |
| BW22GS018779 | BORL14/3/KFA/2*KACHU//KACHU/KIRITATI | A:A | T:T |
| BW22GS018780 | MUNAL #1/KASUKO | A:A | T:T |
| BW22GS018781 | KINGBIRD #1//INQALAB 91*2/TUKURU/3/KASUKO | A:A | T:T |
| BW22GS018782 | KINGBIRD #1//INQALAB 91*2/TUKURU/3/KASUKO | A:A | T:T |
| BW22GS018783 | KINGBIRD #1//INQALAB 91*2/TUKURU/3/KASUKO | A:A | T:T |
| BW22GS018784 | KINGBIRD #1//INQALAB 91*2/TUKURU/6/KSW/SAUAL//SAUAL/3/TRCH/HUIRIVIS #1/5/UP2338*2/SHAMA/3/MILAN/KAUZ//CHIL/CHUM18/4/UP2338*2/SHAMA | A:A | T:T |

| | | | |
|---|---|---|---|
| BW22GS018785 | CHIPAK/KASUKO | A:A | T:T |
| BW22GS018786 | MUCUY/3/BORL14*2//KFA/2*KACHU | A:A | T:T |
| BW22GS018787 | KACHU//KIRITATI/2*TRCH/3/KASUKO | A:A | T:T |
| BW22GS018788 | CHIBIA//PRLII/CM65531/3/FISCAL/4/DANPHE #1/5/CHIBIA//PRLII/CM65531/3/MISR 2/6/KASUKO | A:A | T:T |
| BW22GS018789 | CHIBIA//PRLII/CM65531/3/FISCAL/4/DANPHE #1/5/CHIBIA//PRLII/CM65531/3/MISR 2/6/KASUKO | A:A | T:T |
| BW22GS018790 | TAM200/PASTOR//TOBA97/3/FRNCLN/4/WHEAR//2*PRL/2*PASTOR/5/KASUKO | A:A | T:T |
| BW22GS018791 | TAM200/PASTOR//TOBA97/3/FRNCLN/4/WHEAR//2*PRL/2*PASTOR/5/KASUKO | A:A | T:T |
| BW22GS018792 | SUP152/FRNCLN//KASUKO | A:A | T:T |
| BW22GS018793 | MUNAL*2/WESTONIA//MOKUE #1 | A:A | C:T |
| BW22GS018794 | MUNAL*2/WESTONIA//KASUKO | A:A | C:C |
| BW22GS018795 | MUNAL*2/CHONTE//KASUKO | A:A | T:T |
| BW22GS018796 | MUNAL*2/CHONTE//KASUKO | A:A | T:T |
| BW22GS018797 | KACHU*2/3/ND643//2*PRL/2*PASTOR/4/KASUKO | A:A | T:T |
| BW22GS018798 | BAJ #1*2/5/SW89.5277/BORL95//SKAUZ/3/PRL/2*PASTOR/4/HEILO/6/KASUKO | A:A | T:T |
| BW22GS018799 | KACHU/3/WHEAR//2*PRL/2*PASTOR/4/KASUKO | A:A | T:T |
| BW22GS018800 | KACHU/3/WHEAR//2*PRL/2*PASTOR/4/MOKUE #1 | A:A | T:T |
| BW22GS018801 | KACHU*2/3/PBW343*2/KUKUNA//PBW343*2/KUKUNA/4/BORL14*2//KFA/2*KACHU | A:A | T:T |
| BW22GS018802 | SHA7//PRL/VEE#6/3/FASAN/4/HAAS8446/2*FASAN/5/CBRD/KAUZ/6/MILAN/AMSEL/7/FRET2*2/KUKUNA/8/TRAP #1/BOW/3/VEE/PJN//2*TUI/4/BAV92/RAYON/5/KACHU #1/9/COPIO/10/SHORTENED SR26 TRANSLOCATION//2*WBLL1*2/KKTS/3/BECARD | A:A | T:T |
| BW22GS018803 | TACUPETO F2001/BRAMBLING/5/NAC/TH.AC//3*PVN/3/MIRLO/BUC/4/2*PASTOR*2/6/WAXWING/SRTU//WAXWING/KIRITATI/7/MOKUE #1 | A:A | T:T |
| BW22GS018804 | PBW343*2/KUKUNA*2//KITE/3/ATTILA*2/PBW65*2//YANAC/4/SAUAL/YANAC//SAUAL/6/WAXWING/KIRITATI*2/3/C80.1/3*BATAVIA//2*WBLL1/4/COPIO/5/ND643//2*ATTILA*2/PASTOR/3/WBLL1*2/KURUKU/4/WBLL1*2/BRAMBLING | A:A | T:T |
| BW22GS018805 | BAVIS #1*2/4/PASTOR//HXL7573/2*BAU/3/SOKOLL/WBLL1/5/BORL14 | A:A | C:C |
| BW22GS018806 | NELOKI/4/ATTILA*2/PBW65//PIHA/3/ATTILA/2*PASTOR/8/TACUPETO F2001/6/CNDO/R143//ENTE/MEXI_2/3/AEGILOPS SQUARROSA (TAUS)/4/WEAVER/5/PASTOR/7/ROLF07/9/KASUKO | A:A | T:T |
| BW22GS018807 | PARUS/FRANCOLIN #1/4/MUU #1//PBW343*2/KUKUNA/3/MUU/5/BORL14*2//KFA/2*KACHU | A:A | T:T |
| BW22GS018808 | WHEAR//2*PRL/2*PASTOR/3/WAXBI/4/COPIO/5/NELOKI*2//KACHU/KIRITATI | A:A | T:T |
| BW22GS018809 | WBLL1*2/KKTS//PASTOR/KUKUNA/3/KINGBIRD #1//INQALAB 91*2/TUKURU/5/KAUZ//ALTAR 84/AOS/3/MILAN/KAUZ/4/SAUAL/6/MOKUE #1 | A:A | T:T |

| | | | |
|---|---|---|---|
| BW22GS018810 | WBLL1*2/KKTS//PASTOR/KUKUNA/3/KINGBIRD #1//INQALAB 91*2/TUKURU/5/KAUZ//ALTAR 84/AOS/3/MILAN/KAUZ/4/SAUAL/6/MOKUE #1 | A:A | T:T |
| BW22GS018811 | WBLL1*2/KKTS//PASTOR/KUKUNA/3/KINGBIRD #1//INQALAB 91*2/TUKURU/5/KAUZ//ALTAR 84/AOS/3/MILAN/KAUZ/4/SAUAL/6/MOKUE #1 | A:A | T:T |
| BW22GS018812 | WBLL1*2/KKTS//PASTOR/KUKUNA/3/KINGBIRD #1//INQALAB 91*2/TUKURU/5/KAUZ//ALTAR 84/AOS/3/MILAN/KAUZ/4/SAUAL/6/MOKUE #1 | A:A | T:T |
| BW22GS018813 | WBLL1*2/KKTS//PASTOR/KUKUNA/3/KINGBIRD #1//INQALAB 91*2/TUKURU/5/KAUZ//ALTAR 84/AOS/3/MILAN/KAUZ/4/SAUAL/6/KASUKO | A:A | T:T |
| BW22GS018814 | WBLL1*2/KKTS//PASTOR/KUKUNA/3/KINGBIRD #1//INQALAB 91*2/TUKURU/5/KAUZ//ALTAR 84/AOS/3/MILAN/KAUZ/4/SAUAL/6/BORL14*2//KFA/2*KACHU | A:A | T:T |
| BW22GS018815 | WBLL1*2/KKTS//PASTOR/KUKUNA/3/KINGBIRD #1//INQALAB 91*2/TUKURU/5/KAUZ//ALTAR 84/AOS/3/MILAN/KAUZ/4/SAUAL/6/BORL14*2//KFA/2*KACHU | A:A | T:T |
| BW22GS018816 | WBLL1*2/KKTS//PASTOR/KUKUNA/3/KINGBIRD #1//INQALAB 91*2/TUKURU/5/KAUZ//ALTAR 84/AOS/3/MILAN/KAUZ/4/SAUAL/6/KUTZ//KFA/2*KACHU | A:A | T:T |
| BW22GS018817 | AMUR*2/CIRO16//KASUKO | A:A | T:T |
| BW22GS018818 | FRET2/KUKUNA//FRET2/3/YANAC/4/FRET2/KIRITATI/5/2*UP2338*2/SHAMA/3/MILAN/KAUZ//CHIL/CHUM18/4/UP2338*2/SHAMA/6/KACHU/3/WHEAR//2*PRL/2*PASTOR | A:A | T:T |
| BW22GS018819 | FRET2/KUKUNA//FRET2/3/YANAC/4/FRET2/KIRITATI/5/2*UP2338*2/SHAMA/3/MILAN/KAUZ//CHIL/CHUM18/4/UP2338*2/SHAMA/6/BORL14*2//KFA/2*KACHU | A:A | T:T |
| BW22GS018820 | FRET2/KUKUNA//FRET2/3/YANAC/4/FRET2/KIRITATI/5/2*UP2338*2/SHAMA/3/MILAN/KAUZ//CHIL/CHUM18/4/UP2338*2/SHAMA/6/BORL14*2//KFA/2*KACHU | A:A | T:T |
| BW22GS018821 | FRET2/KUKUNA//FRET2/3/YANAC/4/FRET2/KIRITATI/5/2*UP2338*2/SHAMA/3/MILAN/KAUZ//CHIL/CHUM18/4/UP2338*2/SHAMA/6/BORL14*2//KFA/2*KACHU | A:A | T:T |
| BW22GS018822 | TACUPETO F2001/6/CNDO/R143//ENTE/MEXI_2/3/AEGILOPS SQUARROSA (TAUS)/4/WEAVER/5/PASTOR/7/ROLF07*2/8/SAUAL/YANAC//SAUAL/9/WBLL1*2/KKTS//PASTOR/KUKUNA/3/KINGBIRD #1//INQALAB 91*2/TUKURU/5/KAUZ//ALTAR 84/AOS/3/MILAN/KAUZ/4/SAUAL | A:A | T:T |
| BW22GS018823 | WADER/KASUKO | A:A | T:T |
| BW22GS018824 | WADER/4/KACHU//WBLL1*2/BRAMBLING*2/3/KACHU/KIRITATI | A:A | T:T |
| BW22GS018825 | KFA/5/REH/HARE//2*BCN/3/CROC_1/AE.SQUARROSA (213)//PGO/4/HUITES/6/REH/HARE//2*BCN/3/CROC_1/AE.SQUARROSA (213)//PGO/4/HUITES/7/BOKOTA/8/BOKOTA/9/KFA/2*KACHU//KACHU/KIRITATI | A:A | T:T |
| BW22GS018826 | BECARD/AKURI/4/WBLL1*2/BRAMBLING//JUCHI/3/WBLL1*2/BRAMBLING/5/KASUKO | A:A | T:T |
| BW22GS018827 | KSW/SAUAL//SAUAL/3/2*BORL14 | A:A | T:T |
| BW22GS018828 | KSW/SAUAL//SAUAL/3/2*BORL14 | A:A | T:T |

| | | | |
|---|---|---|---|
| BW22GS018829 | KSW/SAUAL//SAUAL/3/2*BORL14 | A:A | T:T |
| BW22GS018830 | KSW/SAUAL//SAUAL/3/2*BORL14 | A:A | T:T |
| BW22GS018831 | KUTZ//KFA/2*KACHU/3/NADI | A:A | T:T |
| BW22GS018832 | MOKUE #1/CHIPAK | A:A | T:T |
| BW22GS018833 | MOKUE #1/CHIPAK | A:A | T:T |
| BW22GS018834 | MOKUE #1/CHIPAK | A:A | T:T |
| BW22GS018835 | BECARD/QUAIU #1//BORL14/3/BORL14*2//KFA/2*KACHU | A:A | T:T |
| BW22GS018836 | SITE/MO//PASTOR/3/TILHI/4/MUNAL #1/5/MUNAL/6/MUCUY/7/MOKUE #1 | A:A | T:T |
| BW22GS018837 | NADI#1/3/PBW343*2/KUKUNA*2//FRTL/PIFED/4/NADI#2/5/FRANCOLIN #1/3/PBW343*2/KUKUNA*2//YANAC/4/KINGBIRD #1//INQALAB 91*2/TUKURU | A:A | T:T |
| BW22GS018838 | NADI#1/3/PBW343*2/KUKUNA*2//FRTL/PIFED/4/NADI#2/5/FRANCOLIN #1/3/PBW343*2/KUKUNA*2//YANAC/4/KINGBIRD #1//INQALAB 91*2/TUKURU | A:A | T:T |
| BW22GS018839 | NAINA #2/KASUKO | A:A | T:T |
| BW22GS018840 | KACHU #1/3/C80.1/3*BATAVIA//2*WBLL1/4/KACHU/8/TACUPETO F2001/6/CNDO/R143//ENTE/MEXI_2/3/AEGILOPS SQUARROSA (TAUS)/4/WEAVER/5/PASTOR/7/ROLF07/9/KFA/2*KACHU/10/KASUKO | A:A | T:T |
| BW22GS018841 | TACUPETO F2001/6/CNDO/R143//ENTE/MEXI_2/3/AEGILOPS SQUARROSA (TAUS)/4/WEAVER/5/PASTOR/7/ROLF07*2/8/SAUAL/YANAC//SAUAL/9/KASUKO | A:A | T:T |
| BW22GS018842 | BABAX/LR42//BABAX/3/ER2000/5/BABAX/LR39//BABAX*2/4/KABY/BAV92/3/CROC_1/AE.SQUARROSA (224)//OPATA/6/KUTZ//KFA/2*KACHU | A:A | T:T |
| BW22GS018843 | BAJ #1*2/PREMIO//MOKUE #1 | A:A | T:T |
| BW22GS018844 | PBW343*2/KUKUNA//PARUS/3/PBW343*2/KUKUNA/4/BAJ #1/AKURI/5/MOKUE #1 | A:A | T:T |
| BW22GS018845 | TACUPETO F2001/BRAMBLING//KACHU/8/REH/HARE//2*BCN/3/CROC_1/AE.SQUARROSA (213)//PGO/4/HUITES/5/T.DICOCCON PI94624/AE.SQUARROSA (409)//BCN/6/REH/HARE//2*BCN/3/CROC_1/AE.SQUARROSA (213)//PGO/4/HUITES/7/MUTUS/9/SUP152//WBLL1*2/BRAMBLING*2/3/KSW/SAUAL//SA | A:A | T:T |
| BW22GS018846 | TACUPETO F2001/BRAMBLING//KACHU/8/REH/HARE//2*BCN/3/CROC_1/AE.SQUARROSA (213)//PGO/4/HUITES/5/T.DICOCCON PI94624/AE.SQUARROSA (409)//BCN/6/REH/HARE//2*BCN/3/CROC_1/AE.SQUARROSA (213)//PGO/4/HUITES/7/MUTUS/9/SUP152//WBLL1*2/BRAMBLING*2/3/KSW/SAUAL//SA | A:A | T:T |
| BW22GS018847 | TACUPETO F2001/BRAMBLING//KACHU/8/REH/HARE//2*BCN/3/CROC_1/AE.SQUARROSA (213)//PGO/4/HUITES/5/T.DICOCCON PI94624/AE.SQUARROSA (409)//BCN/6/REH/HARE//2*BCN/3/CROC_1/AE.SQUARROSA (213)//PGO/4/HUITES/7/MUTUS/9/SUP152//WBLL1*2/BRAMBLING*2/3/KSW/SAUAL//SA | A:A | T:T |
| BW22GS018848 | SUP152/BAJ #1/3/KINGBIRD #1//INQALAB 91*2/TUKURU/8/ATTILA/3*BCN//BAV92/3/TILHI/4/SUP152/5/SUP152/6/KFA/2*KACHU/7/ATTILA/3*BCN//BAV92 /3/PASTOR/4/TACUPETO F2001*2/BRAMBLING/5/PAURAQ | A:A | T:T |

| | | | |
|---|---|---|---|
| BW22GS018849 | WBLL1*2/BRAMBLING//VORB/FISCAL/3/BECARD/4/ABLEU/5/KASUKO | A:A | T:T |
| BW22GS018850 | BECARD/FRNCLN//BORL14/3/BORL14*2//KFA/2*KACHU | A:A | T:T |
| BW22GS018851 | BORL14*2/FITIS/4/KACHU/3/WHEAR//2*PRL/2*PASTOR | A:A | T:T |
| BW22GS018852 | BORL14*2/3/KBIRD//WBLL1*2/KURUKU/4/KASUKO | A:A | T:T |
| BW22GS018853 | BORL14*2/3/WBLL1*2/TUKURU//CROSBILL #1/4/MOKUE #1 | A:A | T:T |
| BW22GS018854 | BORL14*2/3/WBLL1*2/TUKURU//CROSBILL #1/4/MOKUE #1 | A:A | T:T |
| BW22GS018855 | BORL14*2/3/WBLL1*2/TUKURU//CROSBILL #1/4/MOKUE #1 | A:A | T:T |
| BW22GS018856 | BORL14*2/3/WBLL1*2/TUKURU//CROSBILL #1/4/MOKUE #1 | A:A | T:T |
| BW22GS018857 | BORL14*2/3/WBLL1*2/TUKURU//CROSBILL #1/4/KASUKO | A:A | T:T |
| BW22GS018858 | BORL14*2//KFA/2*KACHU/3/KASUKO | A:A | T:T |
| BW22GS018859 | BORL14*2//BECARD/QUAIU #1/3/KASUKO | A:A | T:T |
| BW22GS018860 | WBLL1*2/BRAMBLING//CHYAK*2/3/KINGBIRD #1//INQALAB 91*2/TUKURU/4/BORL14*2//KFA/2*KACHU | A:A | T:T |
| BW22GS018861 | GRACK/CHYAK/6/ROLF07*2/5/FCT/3/GOV/AZ//MUS/4/DOVE/BUC/7/KACHU/3/WHEAR//2*PRL/2*PASTOR | A:A | T:T |
| BW22GS018862 | GRACK/CHYAK/6/ROLF07*2/5/FCT/3/GOV/AZ//MUS/4/DOVE/BUC/7/KASUKO | A:A | T:T |
| BW22GS018863 | SUP152/HUIRIVIS #1//2*BORL14/3/KASUKO | A:A | T:T |
| BW22GS018864 | SUP152/HUIRIVIS #1//2*BORL14/3/KASUKO | A:A | T:T |
| BW22GS018865 | BECARD/AKURI/3/KACHU//WBLL1*2/BRAMBLING/4/MUTUS/AKURI/5/MOKUE #1 | A:A | T:T |
| BW22GS018866 | BECARD/AKURI/3/KACHU//WBLL1*2/BRAMBLING/4/MUTUS/AKURI/5/BORL14*2//KFA/2*KACHU | A:A | T:T |
| BW22GS018867 | KSW/SAUAL//SAUAL/3/TRCH/HUIRIVIS #1/5/UP2338*2/SHAMA/3/MILAN/KAUZ//CHIL/CHUM18/4/UP2338*2/SHAMA/6/KASUKO | A:A | T:T |
| BW22GS018868 | KSW/SAUAL//SAUAL/3/TRCH/HUIRIVIS #1/5/UP2338*2/SHAMA/3/MILAN/KAUZ//CHIL/CHUM18/4/UP2338*2/SHAMA/6/KSW/SAUAL//SAUAL/3/TRCH/HUIRIVIS #1/5/UP2338*2/SHAMA/3/MILAN/KAUZ//CHIL/CHUM18/4/UP2338*2/SHAMA | A:A | T:T |
| BW22GS018869 | KSW/SAUAL//SAUAL/3/TRCH/HUIRIVIS #1/5/UP2338*2/SHAMA/3/MILAN/KAUZ//CHIL/CHUM18/4/UP2338*2/SHAMA/6/PRL/2*PASTOR//KACHU | A:A | T:T |
| BW22GS018870 | KSW/SAUAL//SAUAL/3/TRCH/HUIRIVIS #1/5/UP2338*2/SHAMA/3/MILAN/KAUZ//CHIL/CHUM18/4/UP2338*2/SHAMA/6/WBLL1*2/KKTS//PASTOR/KUKUNA/3/KINGBIRD #1//INQALAB 91*2/TUKURU/5/KAUZ//ALTAR 84/AOS/3/MILAN/KAUZ/4/SAUAL | A:A | T:T |

| | | | |
|---|---|---|---|
| BW22GS018871 | KSW/SAUAL//SAUAL/3/TRCH/HUIRIVIS #1/5/UP2338*2/SHAMA/3/MILAN/KAUZ//CHIL/CHUM18/4/UP2338*2/SHAMA/6/WBLL1*2/KKTS//PASTOR/KUKUNA/3/KINGBIRD #1//INQALAB 91*2/TUKURU/5/KAUZ//ALTAR 84/AOS/3/MILAN/KAUZ/4/SAUAL | A:A | T:T |
| BW22GS018872 | WBLL1*2/KURUKU//HEILO/3/WBLL1*2/KURUKU/4/SUP152/BAJ #1/5/SUP152/BAJ #1/6/KASUKO | A:A | T:T |
| BW22GS018873 | SUP152/BAJ #1/4/BAJ #1/3/KIRITATI//ATTILA*2/PASTOR/5/SUP152/BAJ #1/6/KSW/SAUAL//SAUAL/3/TRCH/HUIRIVIS #1/5/UP2338*2/SHAMA/3/MILAN/KAUZ//CHIL/CHUM18/4/UP2338*2/SHAMA | A:A | T:T |
| BW22GS018874 | KACHU #1/KIRITATI//KACHU*2/3/GRACK/CHYAK/4/KASUKO | A:A | T:T |
| BW22GS018875 | TUKURU//BAV92/RAYON/3/MUNAL #1/4/2*KFA/2*KACHU/5/BORL14*2//KFA/2*KACHU | A:A | T:T |
| BW22GS018876 | BORL14/MUNAL #1//MOKUE #1 | A:A | T:T |
| BW22GS018877 | BORL14/MUNAL #1//MOKUE #1 | A:A | T:T |
| BW22GS018878 | KACHU//WBLL1*2/BRAMBLING/3/KACHU/KIRITATI/4/NELOKI*2//KACHU/KIRITATI | A:A | T:T |
| BW22GS018879 | FRANCOLIN #1*2/HAWFINCH #1//2*MUCUY/4/MUTUS*2/KINGBIRD #1/3/KSW/SAUAL//SAUAL | A:A | T:T |
| BW22GS018880 | SUP152/QUAIU #2//BECARD/QUAIU #1/6/KSW/SAUAL//SAUAL/3/TRCH/HUIRIVIS #1/5/UP2338*2/SHAMA/3/MILAN/KAUZ//CHIL/CHUM18/4/UP2338*2/SHAMA | A:A | T:T |
| BW22GS018881 | SUP152//WBLL1*2/BRAMBLING*2/3/KSW/SAUAL//SAUAL/4/KASUKO | A:A | T:T |
| BW22GS018882 | ALD/CEP75630//CEP75234/PT7219/3/BUC/BJY/4/MILAN/5/3*BORL14 | A:A | T:T |
| BW22GS018883 | ALD/CEP75630//CEP75234/PT7219/3/BUC/BJY/4/MILAN/5/3*MUCUY | A:A | T:T |
| BW22GS018884 | BORL14/MOKUE #1//KASUKO | A:A | T:T |
| BW22GS018885 | BORL14/MOKUE #1//KASUKO | A:A | T:T |
| BW22GS018886 | AKEPA/KASUKO//KASUKO | A:A | T:T |
| BW22GS018887 | AKEPA/KASUKO//KASUKO | A:A | T:T |
| BW22GS018888 | BOKOTA*2/MOKUE #1 | A:A | T:T |
| BW22GS018889 | BOKOTA*2/MOKUE #1 | A:A | T:T |
| BW22GS018890 | BOKOTA*2/MOKUE #1 | A:A | T:T |
| BW22GS018891 | WHEAR/VIVITSI//WHEAR/3/PANDORA INIA/4/BORL14*2//KFA/2*KACHU/5/BORL14*2//MUNAL #1/FRANCOLIN #1 | A:A | T:T |
| BW22GS018892 | BLOUK #1/4/WHEAR/KUKUNA/3/C80.1/3*BATAVIA//2*WBLL1/5/MUNAL #1*2/6/MOKUE #1 | A:A | T:T |
| BW22GS018893 | BLOUK #1/4/WHEAR/KUKUNA/3/C80.1/3*BATAVIA//2*WBLL1/5/MUNAL #1/6/KASUKO/7/KASUKO | A:A | T:T |
| BW22GS018894 | SUP152/FRNCLN//KASUKO/3/BORL14*2//MUNAL #1/FRANCOLIN #1 | A:A | T:T |
| BW22GS018895 | BAVIS//ATTILA*2/PBW65/3/2*NELOKI*2//KACHU/KIRITATI | A:A | T:T |
| BW22GS018896 | GLADIUS/3/2*KA/NAC//TRCH/4/KUTZ//KFA/2*KACHU/5/KUTZ//KFA/2*KACHU | A:A | T:T |

| | | | |
|---|---|---|---|
| BW22GS018897 | MELON//FILIN/MILAN/3/FILIN/4/PRINIA/PASTOR//HUITES/3/MILAN/OTUS//ATTILA/3*BCN/5/MELON//FILIN/MILAN/3/FILIN/6/BORL14*2//KFA/2*KACHU/7/ROLF07*2/DIAMONDBIRD//TRCH/HUIRIVIS #1/3/BORL14 | A:A | T:T |
| BW22GS018898 | SOKOLL/3/PASTOR//HXL7573/2*BAU/4/MASSIV/PPR47.89C/5/2*BORL14*2//KFA/2*KACHU | A:A | T:T |
| BW22GS018899 | ROLF07/YANAC//TACUPETO F2001/BRAMBLING/5/KAUZ//ALTAR 84/AOS/3/MILAN/KAUZ/4/SAUAL/6/2*BORL14//KFA/2*KACHU | A:A | T:T |
| BW22GS018900 | PRL/2*PASTOR//KACHU*2/3/BORL14*2//KFA/2*KACHU | A:A | T:T |
| BW22GS018901 | SUP152/CIRO16*2//KASUKO | A:A | T:T |
| BW22GS018902 | BAV92//IRENA/KAUZ/3/HUITES/4/PVN/5/CIRO16/6/2*MOKUE #1 | A:A | T:T |
| BW22GS018903 | UP2338*2/SHAMA/3/MILAN/KAUZ//CHIL/CHUM18/4/UP2338*2/SHAMA/5/COPIO*2/6/SHORTENED SR26 TRANSLOCATION//2*WBLL1*2/KKTS/3/BECARD | A:A | T:T |
| BW22GS018904 | UP2338*2/SHAMA/3/MILAN/KAUZ//CHIL/CHUM18/4/UP2338*2/SHAMA/5/COPIO*2/6/SHORTENED SR26 TRANSLOCATION//2*WBLL1*2/KKTS/3/BECARD | A:A | T:T |
| BW22GS018905 | COPIO/5/UP2338*2/SHAMA/3/MILAN/KAUZ//CHIL/CHUM18/4/UP2338*2/SHAMA/6/2*KACHU//WBLL1*2/BRAMBLING/3/KACHU/KIRITATI | A:A | T:T |
| BW22GS018906 | FRANCOLIN #1/3/PBW343*2/KUKUNA*2//YANAC/4/KINGBIRD #1//INQALAB 91*2/TUKURU/5/KASUKO/6/KASUKO | A:A | T:T |
| BW22GS018907 | FRANCOLIN #1/3/PBW343*2/KUKUNA*2//YANAC/4/KINGBIRD #1//INQALAB 91*2/TUKURU/5/BORL14*2//KFA/2*KACHU/6/BORL14*2//MUNAL #1/FRANCOLIN #1 | A:A | T:T |
| BW22GS018908 | NELOKI/5/FRET2/KUKUNA//FRET2/3/TNMU/4/FRET2*2/SHAMA/6/KINGBIRD #1//INQALAB 91*2/TUKURU/7/KASUKO/8/KASUKO | A:A | T:T |
| BW22GS018909 | TACUPETO F2001/BRAMBLING/5/NAC/TH.AC//3*PVN/3/MIRLO/BUC/4/2*PASTOR*2/6/WAXWING/SRTU//WAXWING/KIRITATI/7/KUTZ//KFA/2*KACHU/8/ATTILA/3*BCN//BAV92/3/PASTOR/4/TACUPETO F2001*2/BRAMBLING/5/PAURAQ/6/KFA/2*KACHU | A:A | T:T |
| BW22GS018910 | TACUPETO F2001/BRAMBLING/5/NAC/TH.AC//3*PVN/3/MIRLO/BUC/4/2*PASTOR*2/6/WAXWING/SRTU//WAXWING/KIRITATI/7/2*SUP152//WBLL1*2/BRAMBLING*2/3/KSW/SAUAL//SAUAL | NA | T:T |
| BW22GS018911 | TACUPETO F2001/BRAMBLING/5/NAC/TH.AC//3*PVN/3/MIRLO/BUC/4/2*PASTOR*2/6/WAXWING/SRTU//WAXWING/KIRITATI/7/2*SUP152//WBLL1*2/BRAMBLING*2/3/KSW/SAUAL//SAUAL | A:A | T:T |
| BW22GS018912 | SAUAL/MUTUS*2//CIRO16*2/3/MOKUE #1 | A:A | T:T |
| BW22GS018913 | SAUAL/MUTUS*2//CIRO16*2/3/MOKUE #1 | A:A | T:T |
| BW22GS018914 | SAUAL/MUTUS*2//CIRO16*2/3/MOKUE #1 | A:A | T:T |
| BW22GS018915 | SAUAL/MUTUS*2//CIRO16*2/3/MOKUE #1 | A:A | T:T |
| BW22GS018916 | SAUAL/MUTUS*2//CIRO16*2/3/MOKUE #1 | A:A | T:T |

| | | | |
|---|---|---|---|
| BW22GS018917 | QUAIU #2/BAVIS #1//2*KASUKO | A:A | T:T |
| BW22GS018918 | NELOKI/4/ATTILA*2/PBW65//PIHA/3/ATTILA/2*PASTOR/8/TACUPETO F2001/6/CNDO/R143//ENTE/MEXI_2/3/AEGILOPS SQUARROSA (TAUS)/4/WEAVER/5/PASTOR/7/ROLF07/9/2*KASUKO | A:A | T:T |
| BW22GS018919 | NELOKI/4/ATTILA*2/PBW65//PIHA/3/ATTILA/2*PASTOR/8/TACUPETO F2001/6/CNDO/R143//ENTE/MEXI_2/3/AEGILOPS SQUARROSA (TAUS)/4/WEAVER/5/PASTOR/7/ROLF07/9/2*KASUKO | A:A | T:T |
| BW22GS018920 | SHA7/VEE#5//ARIV92/3/PBW343*2/KUKUNA/4/2*VARIS/MISR 2/3/FRET2/KUKUNA//FRET2/5/2*KASUKO | A:A | T:T |
| BW22GS018921 | WBLL1*2/KKTS//PASTOR/KUKUNA/3/KINGBIRD #1//INQALAB 91*2/TUKURU/5/KAUZ//ALTAR 84/AOS/3/MILAN/KAUZ/4/SAUAL*2/6/BORL14*2//KFA/2*KACHU | A:A | T:T |
| BW22GS018922 | BECARD/AKURI*2/3/KINGBIRD #1//INQALAB 91*2/TUKURU*2/4/NELOKI*2//KACHU/KIRITATI | A:A | T:T |
| BW22GS018923 | KFA/5/REH/HARE//2*BCN/3/CROC_1/AE.SQUARROSA (213)//PGO/4/HUITES/6/REH/HARE//2*BCN/3/CROC_1/AE.SQUARROSA (213)//PGO/4/HUITES/7/BOKOTA/8/BOKOTA/9/BORL14*2//KFA/2*KACHU/10/BORL14*2//KFA/2*KACHU | A:A | T:T |
| BW22GS018924 | KFA/5/REH/HARE//2*BCN/3/CROC_1/AE.SQUARROSA (213)//PGO/4/HUITES/6/REH/HARE//2*BCN/3/CROC_1/AE.SQUARROSA (213)//PGO/4/HUITES/7/BOKOTA/8/BOKOTA*2/9/KACHU*2/3/ND643//2*PRL/2*PASTOR | A:A | T:T |
| BW22GS018925 | BECARD/AKURI/4/WBLL1*2/BRAMBLING//JUCHI/3/WBLL1*2/BRAMBLING/5/KASUKO/6/KASUKO | A:A | T:T |
| BW22GS018926 | KASUKO*2/3/PRL/2*PASTOR//KACHU | A:A | T:T |
| BW22GS018927 | KUTZ//KFA/2*KACHU/3/NADI/5/NADI#1/3/PBW343*2/KUKUNA*2//FRTL/PIFED/4/NADI#2 | A:A | T:T |
| BW22GS018928 | BECARD/QUAIU #1//BORL14/3/2*BORL14*2//KFA/2*KACHU | A:A | T:T |
| BW22GS018929 | AMUR/3/KINGBIRD #1//INQALAB 91*2/TUKURU/4/AMUR*2/5/BORL14*2//KFA/2*KACHU | A:A | T:T |
| BW22GS018930 | WAXWING/KIRITATI*2/3/C80.1/3*BATAVIA//2*WBLL1/4/COPIO/5/ND643//2*ATTILA*2/PASTOR/3/WBLL1*2/KURUKU/4/WBLL1*2/BRAMBLING/6/BORL14/7/KASUKO | A:A | T:T |
| BW22GS018931 | WAXWING/KIRITATI*2/3/C80.1/3*BATAVIA//2*WBLL1/4/COPIO/5/ND643//2*ATTILA*2/PASTOR/3/WBLL1*2/KURUKU/4/WBLL1*2/BRAMBLING/6/BORL14/7/KASUKO | A:A | T:T |
| BW22GS018932 | WBLL1*2/BRAMBLING//JUCHI/5/KIRITATI/4/2*BAV92//IRENA/KAUZ/3/HUITES/6/WBLL1*2/BRAMBLING//KACHU*2/7/KUTZ//KFA/2*KACHU | A:A | T:T |
| BW22GS018933 | LIVINGSTON/6/2*MTRWA92.161/PRINIA/5/SERI*3//RL6010/4*YR/3/PASTOR/4/BAV92*2/7/KASUKO | A:A | T:T |
| BW22GS018934 | LIVINGSTON/6/2*MTRWA92.161/PRINIA/5/SERI*3//RL6010/4*YR/3/PASTOR/4/BAV92*2/7/SUP152//WBLL1*2/BRAMBLING*2/3/KSW/SAUAL//SAUAL | A:A | T:T |

| | | | |
|---|---|---|---|
| BW22GS018935 | C80.1/3*BATAVIA//2*WBLL1/5/REH/HARE//2*BCN/3/CROC_1/AE.SQUARROSA (213)//PGO/4/HUITES/6/PBW343*2/KUKUNA*2//FRTL/PIFED/7/C80.1/3*BATAVIA//2*WBLL1/5/REH/HARE//2*BCN/3/CROC_1/AE.SQUARROSA (213)//PGO/4/HUITES/8/KASUKO/9/BORL14*2//MUNAL #1/FRANCOLIN #1 | A:A | T:T |
| BW22GS018936 | TACUPETO F2001/6/CNDO/R143//ENTE/MEXI_2/3/AEGILOPS SQUARROSA (TAUS)/4/WEAVER/5/PASTOR/7/ROLF07*2/8/SAUAL/YANAC//SAUAL*2/9/KASUKO | A:A | T:T |
| BW22GS018937 | TACUPETO F2001/BRAMBLING//KACHU/8/REH/HARE//2*BCN/3/CROC_1/AE.SQUARROSA (213)//PGO/4/HUITES/5/T.DICOCCON PI94624/AE.SQUARROSA (409)//BCN/6/REH/HARE//2*BCN/3/CROC_1/AE.SQUARROSA (213)//PGO/4/HUITES/7/MUTUS*2/9/BORL14*2//KFA/2*KACHU | A:A | T:T |
| BW22GS018938 | TACUPETO F2001/BRAMBLING//KACHU/8/REH/HARE//2*BCN/3/CROC_1/AE.SQUARROSA (213)//PGO/4/HUITES/5/T.DICOCCON PI94624/AE.SQUARROSA (409)//BCN/6/REH/HARE//2*BCN/3/CROC_1/AE.SQUARROSA (213)//PGO/4/HUITES/7/MUTUS*2/9/BORL14*2//KFA/2*KACHU | A:A | T:T |
| BW22GS018939 | TACUPETO F2001/BRAMBLING//KACHU/8/REH/HARE//2*BCN/3/CROC_1/AE.SQUARROSA (213)//PGO/4/HUITES/5/T.DICOCCON PI94624/AE.SQUARROSA (409)//BCN/6/REH/HARE//2*BCN/3/CROC_1/AE.SQUARROSA (213)//PGO/4/HUITES/7/MUTUS*2/9/BORL14*2//KFA/2*KACHU | A:A | T:T |
| BW22GS018940 | TACUPETO F2001/BRAMBLING//KACHU/8/REH/HARE//2*BCN/3/CROC_1/AE.SQUARROSA (213)//PGO/4/HUITES/5/T.DICOCCON PI94624/AE.SQUARROSA (409)//BCN/6/REH/HARE//2*BCN/3/CROC_1/AE.SQUARROSA (213)//PGO/4/HUITES/7/MUTUS*2/9/BORL14*2//KFA/2*KACHU | A:A | T:T |
| BW22GS018941 | TACUPETO F2001/BRAMBLING//KACHU/8/REH/HARE//2*BCN/3/CROC_1/AE.SQUARROSA (213)//PGO/4/HUITES/5/T.DICOCCON PI94624/AE.SQUARROSA (409)//BCN/6/REH/HARE//2*BCN/3/CROC_1/AE.SQUARROSA (213)//PGO/4/HUITES/7/MUTUS*2/9/SUP152//WBLL1*2/BRAMBLING*2/3/KSW/SAUAL// | A:A | T:T |
| BW22GS018942 | PBW343*2/KUKUNA*2//FRTL/PIFED/3/ABLEU*2/4/WBLL1*2/KUKUNA//KIRITATI/2*TRCH/3/BAJ #1/AKURI | A:A | T:T |
| BW22GS018943 | FRANCOLIN #1//WBLL1*2/KURUKU/3/WBLL1*2/BRAMBLING//CHYAK*2/4/SUP152//WBLL1*2/BRAMBLING*2/3/KSW/SAUAL//SAUAL | A:A | T:T |
| BW22GS018944 | BORL14*2/MUNAL #1/4/SHORTENED SR26 TRANSLOCATION//2*WBLL1*2/KKTS/3/BECARD/5/BORL14*2/MUNAL #1 | A:A | T:T |
| BW22GS018945 | CIANO M2018/4/KACHU*2/3/ND643//2*PRL/2*PASTOR/5/BORL14*2//KFA/2*KACHU | A:A | T:T |
| BW22GS018946 | BLOUK #1/MUNAL/3/WBLL1*2/SHAMA//BAJ #1/4/SUP152/BAJ #1/5/KASUKO/6/KASUKO | A:A | T:T |
| BW22GS018947 | PRL/2*PASTOR*2//SKAUZ/BAV92/3/2*BECARD//ND643/2*WBLL1*2/4/KUTZ*2//KFA/2*KACHU | A:A | T:T |

| | | | |
|---|---|---|---|
| BW22GS018948 | SUP152/BLOUK #1/3/PRL/2*PASTOR*2//VORB/4/SUP152/BLOUK #1*2/5/BORL14*2//KFA/2*KACHU | A:A | C:T |
| BW22GS018949 | BECARD/AKURI/3/KACHU//WBLL1*2/BRAMBLING/4/MUTUS/AKURI*2/5/BORL14*2//KFA/2*KACHU | A:A | T:T |
| BW22GS018950 | BECARD/AKURI/4/WBLL1*2/BRAMBLING//JUCHI/3/WBLL1*2/BRAMBLING/5/BOKOTA/6/KASUKO/7/KASUKO | A:A | T:T |
| BW22GS018951 | KACHU/BECARD//WBLL1*2/BRAMBLING/4/FRET2/TUKURU//FRET2/3/MUNAL #1*2/6/KSW/SAUAL//SAUAL/3/TRCH/HUIRIVIS #1/5/UP2338*2/SHAMA/3/MILAN/KAUZ//CHIL/CHUM18/4/UP2338*2/SHAMA | A:A | T:T |
| BW22GS018952 | KACHU/BECARD//WBLL1*2/BRAMBLING/4/FRET2/TUKURU//FRET2/3/MUNAL #1*2/6/KSW/SAUAL//SAUAL/3/TRCH/HUIRIVIS #1/5/UP2338*2/SHAMA/3/MILAN/KAUZ//CHIL/CHUM18/4/UP2338*2/SHAMA | A:A | T:T |
| BW22GS018953 | ONIX/KBIRD//BORL14/3/ONIX/KBIRD/4/KUTZ//KFA/2*KACHU/5/KUTZ//KFA/2*KACHU | A:A | T:T |
| BW22GS018954 | KACHU #1/3/T.DICOCCON PI94624/AE.SQUARROSA (409)//BCN/4/2*KACHU/5/MUTUS*2/TECUE #1/6/MUTUS*2/TECUE #1*2/7/MOKUE #1 | A:A | T:T |
| BW22GS018955 | MOKUE*2/7/KACHU #1/3/T.DICOCCON PI94624/AE.SQUARROSA (409)//BCN/4/2*KACHU/5/MUTUS*2/TECUE #1/6/MUTUS*2/TECUE #1 | A:A | T:T |
| BW22GS018956 | SUP152/QUAIU #2//BECARD/QUAIU #1/3/KUTZ*2//KFA/2*KACHU/4/KUTZ//KFA/2*KACHU | A:A | T:T |
| BW22GS018957 | SUP152/QUAIU #2//BECARD/QUAIU #1/7/ATTILA/3*BCN//BAV92/3/PASTOR/4/TACUPETO F2001*2/BRAMBLING/5/PAURAQ/6/KFA/2*KACHU/8/KFA/2*KACHU*2//MISR 1 | A:A | T:T |
| BW22GS018958 | SUP152//WBLL1*2/BRAMBLING*2/3/KSW/SAUAL//SAUAL/4/BORL14*2//KFA/2*KACHU/5/BORL14*2//KFA/2*KAC HU | A:A | T:T |
| BW22GS018959 | SUP152//WBLL1*2/BRAMBLING*2/3/KSW/SAUAL//SAUAL*2/8/ATTILA/3*BCN//BAV92/3/TILHI/4/SUP152/5/SUP1 52/6/KFA/2*KACHU/7/ATTILA/3*BCN//BAV92/3/PASTOR/4/TACUPETO F2001*2/BRAMBLING/5/PAURAQ | A:A | T:T |
| BW22GS018960 | BECARD//ND643/2*WBLL1*2/3/KSW/SAUAL//SAUAL*2/4/KFA/2*KACHU*2//SUP152 | A:A | T:T |
| BW22GS018961 | MACE*2/3/KFA/2*KACHU*2//SUP152 | A:A | T:T |
| BW22GS018962 | SI-K56/9/BECARD #1/8/BOW/VEE/5/ND/VG9144//KAL/BB/3/YACO/4/CHIL/6/CASKOR/3/CROC_1/AE.SQUARROSA (224)//OPATA/7/PASTOR//MILAN/KAUZ/3/BAV92/10/ATTILA/3*BCN//BAV92/3/TILHI/4/SUP152/5/SUP152/6/KFA /2*KACHU/7/ATTILA/3*BCN//BAV92/3/PASTOR/4/TACUPETO F2001*2/B | A:A | T:T |
| BW22GS018963 | BORL14*2//BECARD/QUAIU #1/3/MOKUE #1 | A:A | T:T |
| BW22GS018964 | KAKURU #1 | A:A | T:T |
| BW22GS019306 | BONSU | A:A | C:C |
| BW22GS019307 | BECARD//ND643/2*WBLL1*2/3/BORL14 | A:A | T:T |

| | | | |
|---|---|---|---|
| BW22GS019308 | BORL14*2/BLANCA GRANDE 515 | A:A | T:T |
| BW22GS019309 | MUNAL #1/CHIPAK//KASUKO | A:A | T:T |
| BW22GS019310 | KAKURU/KASUKO//KASUKO | A:A | T:T |
| BW22GS019311 | KACHU*2/3/ND643//2*PRL/2*PASTOR*2/4/PRL/2*PASTOR*2//VORB | A:A | T:T |
| BW22GS019312 | KACHU*2/3/ND643//2*PRL/2*PASTOR*2/4/PRL/2*PASTOR*2//VORB | A:A | T:T |
| BW22GS019313 | PBW343*2/KUKUNA//PBW343*2/KUKUNA/3/WBLL1*2/SHAMA//KACHU/4/KASUKO/5/KASUKO | A:A | T:T |
| BW22GS019314 | SAUAL*2/6/CNDO/R143//ENTE/MEXI_2/3/AEGILOPS SQUARROSA (TAUS)/4/WEAVER/5/2*PASTOR/7/PBW343*2/KUKUNA*2//FRTL/PIFED/8/BORL14/9/KASUKO | A:A | T:T |
| BW22GS019315 | SAUAL*2/6/CNDO/R143//ENTE/MEXI_2/3/AEGILOPS SQUARROSA (TAUS)/4/WEAVER/5/2*PASTOR/7/PBW343*2/KUKUNA*2//FRTL/PIFED/8/BORL14/9/KASUKO | A:A | T:T |
| BW22GS019316 | FRANCOLIN #1/3/PBW343*2/KUKUNA*2//YANAC/4/KINGBIRD #1//INQALAB 91*2/TUKURU*2/5/PRL/2*PASTOR*2//VORB | A:A | T:T |
| BW22GS019317 | SAUAL/MUTUS/4/KACHU #1//WBLL1*2/KUKUNA/3/BRBT1*2/KIRITATI/5/2*KACHU/SAUAL*2//COPIO | A:A | T:T |
| BW22GS019318 | SAUAL/MUTUS/4/KACHU #1//WBLL1*2/KUKUNA/3/BRBT1*2/KIRITATI/5/2*KACHU/SAUAL*2//COPIO | A:A | T:T |
| BW22GS019319 | KASUKO/NADI#2//KASUKO | A:A | NA |
| BW22GS019320 | KASUKO/NADI#2//KASUKO | A:A | T:T |
| BW22GS019321 | KASUKO*2/5/MUTUS/DANPHE #1/4/C80.1/3*BATAVIA//2*WBLL1/3/C80.1/3*QT4522//2*PASTOR | A:A | T:T |
| BW22GS019322 | KASUKO/4/CIRO16*2/3/MUU #1/SAUAL//MUU/5/KASUKO | A:A | T:T |
| BW22GS019323 | KASUKO/4/CIRO16*2/3/MUU #1/SAUAL//MUU/5/KASUKO | A:A | T:T |
| BW22GS019324 | BORL14/KASUKO | A:A | T:T |
| BW22GS019325 | CHIPAK/4/SUP152//WBLL1*2/BRAMBLING*2/3/KSW/SAUAL//SAUAL | A:A | T:T |
| BW22GS019326 | KACHU//KIRITATI/2*TRCH/3/KASUKO | A:A | T:T |
| BW22GS019327 | KACHU//KIRITATI/2*TRCH/3/KASUKO | A:A | T:T |
| BW22GS019328 | BABAX/LR42//BABAX*2/3/SHAMA/4/WAXWING*2/KRONSTAD F2004/5/KASUKO | A:A | T:T |
| BW22GS019329 | KACHU*2/3/ND643//2*PRL/2*PASTOR/4/KASUKO | A:A | T:T |
| BW22GS019330 | PRL/2*PASTOR//KACHU/4/KACHU*2/3/ND643//2*PRL/2*PASTOR | A:A | T:T |
| BW22GS019331 | PRL/2*PASTOR//KACHU/3/KASUKO | A:A | T:T |
| BW22GS019332 | PRL/2*PASTOR//KACHU/3/KASUKO | A:A | T:T |
| BW22GS019333 | SUP152/CIRO16//KASUKO | A:A | T:T |
| BW22GS019334 | NELOKI/5/FRET2/KUKUNA//FRET2/3/TNMU/4/FRET2*2/SHAMA/6/KINGBIRD #1//INQALAB 91*2/TUKURU/7/KASUKO | A:A | T:T |

| | | | |
|---|---|---|---|
| BW22GS019335 | PARUS/FRANCOLIN #1/4/MUU #1//PBW343*2/KUKUNA/3/MUU/5/KASUKO | A:A | T:T |
| BW22GS019336 | BECARD/AKURI*2/3/PBW343*2/KUKUNA*2//FRTL/PIFED/4/SUP152//WBLL1*2/BRAMBLING*2/3/KSW/SAUAL//SAUAL | A:A | T:T |
| BW22GS019337 | BECARD/AKURI*2/3/PBW343*2/KUKUNA*2//FRTL/PIFED/4/SUP152//WBLL1*2/BRAMBLING*2/3/KSW/SAUAL//SAUAL | A:A | T:T |
| BW22GS019338 | CTRIGO/5/KAUZ//ALTAR 84/AOS/3/KAUZ/4/SW94.15464/6/2*UP2338*2/SHAMA/3/MILAN/KAUZ//CHIL/CHUM18/4/UP2338*2/SHAMA/7/KASUKO | A:A | T:T |
| BW22GS019339 | UP2338*2/SHAMA/3/MILAN/KAUZ//CHIL/CHUM18/4/UP2338*2/SHAMA/5/UP2338*2/VIVITSI/3/FRET2/TUKURU//FRET2/4/MISR 1/6/KASUKO | A:A | T:T |
| BW22GS019340 | KACHU/KIRITATI//BORL14/3/KASUKO | A:A | T:T |
| BW22GS019341 | TC870344/GUI//TEMPORALERA M 87/AGR/3/2*WBLL1/8/BOW/VEE/5/ND/VG9144//KAL/BB/3/YACO/4/CHIL/6/CASKOR/3/CROC_1/AE.SQUARROSA (224)//OPATA/7/PASTOR//MILAN/KAUZ/3/BAV92/9/KASUKO | A:A | T:T |
| BW22GS019342 | TC870344/GUI//TEMPORALERA M 87/AGR/3/2*WBLL1/8/BOW/VEE/5/ND/VG9144//KAL/BB/3/YACO/4/CHIL/6/CASKOR/3/CROC_1/AE.SQUARROSA (224)//OPATA/7/PASTOR//MILAN/KAUZ/3/BAV92/9/KASUKO | A:A | T:T |
| BW22GS019343 | KFA/2*KACHU/4/WBLL1*2/KURUKU//KRONSTAD F2004/3/WBLL1*2/BRAMBLING/5/KASUKO | A:A | T:T |
| BW22GS019344 | KFA/2*KACHU/4/WBLL1*2/KURUKU//KRONSTAD F2004/3/WBLL1*2/BRAMBLING/5/SUP152//WBLL1*2/BRAMBLING*2/3/KSW/SAUAL//SAUAL | A:A | T:T |
| BW22GS019345 | KFA/2*KACHU/4/WBLL1*2/KURUKU//KRONSTAD F2004/3/WBLL1*2/BRAMBLING/5/KUTZ//KFA/2*KACHU | A:A | T:T |
| BW22GS019346 | TACUPETO F2001/BRAMBLING//KIRITATI/3/FRANCOLIN #1/BLOUK #1/4/FRANCOLIN #1/BLOUK #1/5/SHORTENED SR26 TRANSLOCATION//2*WBLL1*2/KKTS/3/BECARD | A:A | T:T |
| BW22GS019347 | TACUPETO F2001/6/CNDO/R143//ENTE/MEXI_2/3/AEGILOPS SQUARROSA (TAUS)/4/WEAVER/5/PASTOR/7/ROLF07*2/8/SAUAL/YANAC//SAUAL/9/KASUKO | A:A | T:T |
| BW22GS019348 | TACUPETO F2001/6/CNDO/R143//ENTE/MEXI_2/3/AEGILOPS SQUARROSA (TAUS)/4/WEAVER/5/PASTOR/7/ROLF07*2/8/SAUAL/YANAC//SAUAL/9/KASUKO | A:A | T:T |
| BW22GS019349 | TACUPETO F2001/6/CNDO/R143//ENTE/MEXI_2/3/AEGILOPS SQUARROSA (TAUS)/4/WEAVER/5/PASTOR/7/ROLF07*2/8/SAUAL/YANAC//SAUAL/9/SUP152//WBLL1*2/BRAMBLING*2/3/KSW/SAUAL//SAUAL | A:A | T:T |
| BW22GS019350 | TACUPETO F2001/6/CNDO/R143//ENTE/MEXI_2/3/AEGILOPS SQUARROSA (TAUS)/4/WEAVER/5/PASTOR/7/ROLF07*2/8/SAUAL/YANAC//SAUAL/9/SUP152//WBLL1*2/BRAMBLING*2/3/KSW/SAUAL//SAUAL | A:A | T:T |

| | | | |
|---|---|---|---|
| BW22GS019351 | TACUPETO F2001/6/CNDO/R143//ENTE/MEXI_2/3/AEGILOPS SQUARROSA (TAUS)/4/WEAVER/5/PASTOR/7/ROLF07*2/8/SAUAL/YANAC//SAUAL/9/SUP152//WBLL1*2/BRAMBLING*2/3/KSW/SAUAL//SAUAL | A:A | T:T |
| BW22GS019352 | SAUAL/MUTUS/4/KACHU #1//WBLL1*2/KUKUNA/3/BRBT1*2/KIRITATI/5/BORL14//KFA/2*KACHU | A:A | T:T |
| BW22GS019353 | SAUAL/MUTUS/4/KACHU #1//WBLL1*2/KUKUNA/3/BRBT1*2/KIRITATI/5/KUTZ//KFA/2*KACHU | A:A | T:T |
| BW22GS019354 | SAUAL/MUTUS/4/KACHU #1//WBLL1*2/KUKUNA/3/BRBT1*2/KIRITATI/5/KUTZ//KFA/2*KACHU | A:A | T:T |
| BW22GS019355 | SAUAL/MUTUS/4/KACHU #1//WBLL1*2/KUKUNA/3/BRBT1*2/KIRITATI/5/KUTZ//KFA/2*KACHU | A:A | T:T |
| BW22GS019356 | KASUKO/MOKUE #1 | A:A | T:T |
| BW22GS019357 | ND643//2*ATTILA*2/PASTOR/3/WBLL1*2/KURUKU/4/WBLL1*2/BRAMBLING/6/BABAX/LR42//BABAX*2/3/KUKUNA/4/CROSBILL #1/5/BECARD/7/KASUKO | A:A | T:T |
| BW22GS019358 | ND643//2*ATTILA*2/PASTOR/3/WBLL1*2/KURUKU/4/WBLL1*2/BRAMBLING/6/BABAX/LR42//BABAX*2/3/KUKUNA/4/CROSBILL #1/5/BECARD/7/KASUKO | A:A | T:T |
| BW22GS019359 | WBLL1*2/KUKUNA//KIRITATI/3/WBLL1*2/KUKUNA/4/KINGBIRD #1//INQALAB 91*2/TUKURU/5/WBLL1*2/BRAMBLING//KACHU/6/KASUKO | A:A | T:T |
| BW22GS019360 | WBLL1*2/BRAMBLING//JUCHI/3/KINGBIRD #1//INQALAB 91*2/TUKURU/4/WBLL1*2/BRAMBLING//KACHU/5/KASUKO | A:A | T:T |
| BW22GS019361 | FRANCOLIN #1//WBLL1*2/KURUKU/3/WBLL1*2/BRAMBLING//CHYAK/4/SUP152//WBLL1*2/BRAMBLING*2/3/KSW/SAUAL//SAUAL | A:A | C:C |
| BW22GS019362 | FRANCOLIN #1//WBLL1*2/KURUKU/3/WBLL1*2/BRAMBLING//CHYAK/4/SUP152//WBLL1*2/BRAMBLING*2/3/KSW/SAUAL//SAUAL | A:A | C:C |
| BW22GS019363 | SUP152/BAJ #1//KFA/2*KACHU/10/KACHU #1/3/C80.1/3*BATAVIA//2*WBLL1/4/KACHU/8/TACUPETO F2001/6/CNDO/R143//ENTE/MEXI_2/3/AEGILOPS SQUARROSA (TAUS)/4/WEAVER/5/PASTOR/7/ROLF07/9/KFA/2*KACHU | A:A | T:T |
| BW22GS019364 | SUP152/BAJ #1//KFA/2*KACHU/10/KACHU #1/3/C80.1/3*BATAVIA//2*WBLL1/4/KACHU/8/TACUPETO F2001/6/CNDO/R143//ENTE/MEXI_2/3/AEGILOPS SQUARROSA (TAUS)/4/WEAVER/5/PASTOR/7/ROLF07/9/KFA/2*KACHU | A:A | T:T |

| | | | |
|---|---|---|---|
| BW22GS019365 | SUP152/BAJ #1//KFA/2*KACHU/10/KACHU #1/3/C80.1/3*BATAVIA//2*WBLL1/4/KACHU/8/TACUPETO F2001/6/CNDO/R143//ENTE/MEXI_2/3/AEGILOPS SQUARROSA (TAUS)/4/WEAVER/5/PASTOR/7/ROLF07/9/KFA/2*KACHU | A:A | T:T |
| BW22GS019366 | KACHU/BECARD//WBLL1*2/BRAMBLING/3/KACHU/KINDE/4/KASUKO | A:A | T:T |
| BW22GS019367 | BORL14*2/FITIS//KASUKO | A:A | T:T |
| BW22GS019368 | BORL14*2/FITIS//KASUKO | A:A | T:T |
| BW22GS019369 | GRACK/CHYAK/6/ROLF07*2/5/FCT/3/GOV/AZ//MUS/4/DOVE/BUC/7/SUP152//WBLL1*2/BRAMBLING*2/3/KSW/SAUAL//SAUAL | A:A | T:T |
| BW22GS019370 | MUTUS//WBLL1*2/BRAMBLING/3/WBLL1*2/BRAMBLING/4/KFA/2*KACHU/5/KASUKO | A:A | T:T |
| BW22GS019371 | KACHU #1/KIRITATI//KACHU*2/3/GRACK/CHYAK/4/KASUKO | A:A | T:T |
| BW22GS019372 | KUTZ//KFA/2*KACHU/3/KASUKO | A:A | T:T |
| BW22GS019373 | KUTZ//KFA/2*KACHU/4/SUP152//WBLL1*2/BRAMBLING*2/3/KSW/SAUAL//SAUAL | A:A | T:T |
| BW22GS019374 | MUNAL/WESTONIA//SUP152/BAJ #1/3/MOKUE #1 | A:A | T:T |
| BW22GS019375 | MUNAL/WESTONIA//SUP152/BAJ #1/4/SUP152//WBLL1*2/BRAMBLING*2/3/KSW/SAUAL//SAUAL | A:A | T:T |
| BW22GS019376 | MUTUS//WBLL1*2/BRAMBLING/3/WBLL1*2/BRAMBLING/4/CHYAK1/GRACK/5/KASUKO | A:A | T:T |
| BW22GS019377 | MUTUS//WBLL1*2/BRAMBLING/3/WBLL1*2/BRAMBLING*2/4/KACHU/KIRITATI/5/KUTZ//KFA/2*KACHU | A:A | T:T |
| BW22GS019378 | ROLF07/HUIRIVIS #1//TACUPETO F2001*2/KIRITATI/3/TRCH/HUIRIVIS #1/4/BORL14/5/MOKUE #1 | A:A | T:T |
| BW22GS019379 | Pavon 76 | A:A | T:T |
| BW22GS019380 | Pavon 76 | A:A | T:T |
| BW22GS019381 | KACHU//KIRITATI/2*TRCH/3/KASUKO/4/KASUKO | A:A | T:T |
| BW22GS019382 | BAVIS/NAVJ07/4/2*SUP152//WBLL1*2/BRAMBLING*2/3/KSW/SAUAL//SAUAL | A:A | T:T |
| BW22GS019383 | BONSU/4/2*SUP152//WBLL1*2/BRAMBLING*2/3/KSW/SAUAL//SAUAL | A:A | T:T |
| BW22GS019384 | MELON//FILIN/MILAN/3/FILIN/4/TRCH/SRTU//KACHU/5/KASUKO/6/KASUKO | A:A | T:T |
| BW22GS019385 | TILHI/SOKOLL*2//KINGBIRD #1/7/KFA/2*KACHU/5/WBLL1*2/4/BABAX/LR42//BABAX/3/BABAX/LR42//BABAX/6/KFA/2*KACHU/8/BORL14*2//MUNAL #1/FRANCOLIN #1 | A:A | T:T |
| BW22GS019386 | MUTUS//ND643/2*WBLL1*2/3/ONIX/KBIRD*2//KFA/2*KACHU | A:A | T:T |
| BW22GS019387 | BAACH/KASUKO//KASUKO | A:A | T:T |
| BW22GS019388 | FRANCOLIN #1/3/PBW343*2/KUKUNA*2//YANAC/4/KINGBIRD #1//INQALAB 91*2/TUKURU/5/2*SUP152//WBLL1*2/BRAMBLING*2/3/KSW/SAUAL//SAUAL | A:A | T:T |
| BW22GS019389 | TACUPETO F2001/BRAMBLING/5/NAC/TH.AC//3*PVN/3/MIRLO/BUC/4/2*PASTOR*2/6/WAXWING/SRTU//WAXWING/KIRITATI/7/2*KASUKO | A:A | T:T |

| | | | |
|---|---|---|---|
| BW22GS019390 | TACUPETO F2001/BRAMBLING/5/NAC/TH.AC//3*PVN/3/MIRLO/BUC/4/2*PASTOR*2/6/WAXWING/SRTU//WAXWING/KIRITATI/7/2*KASUKO | A:A | T:T |
| BW22GS019391 | TACUPETO F2001/BRAMBLING/5/NAC/TH.AC//3*PVN/3/MIRLO/BUC/4/2*PASTOR*2/6/WAXWING/SRTU//WAXWING/KIRITATI/7/2*KASUKO | A:A | T:T |
| BW22GS019392 | KAUZ//ALTAR 84/AOS/3/MILAN/KAUZ/4/SAUAL/5/TRCH/SRTU//KACHU/6/KACHU/SAUAL/7/2*SUP152//WBLL1*2/BRAMBLING*2/3/KSW/SAUAL//SAUAL | A:A | T:T |
| BW22GS019393 | SAUAL/MUTUS//KINGBIRD #1/3/SAUAL/MUTUS/4/2*KASUKO | A:A | T:T |
| BW22GS019394 | NELOKI/4/ATTILA*2/PBW65//PIHA/3/ATTILA/2*PASTOR/8/TACUPETO F2001/6/CNDO/R143//ENTE/MEXI_2/3/AEGILOPS SQUARROSA (TAUS)/4/WEAVER/5/PASTOR/7/ROLF07/9/2*KASUKO | A:A | T:T |
| BW22GS019395 | C80.1/3*BATAVIA//2*WBLL1/5/REH/HARE//2*BCN/3/CROC_1/AE.SQUARROSA (213)//PGO/4/HUITES/6/PBW343*2/KUKUNA*2//FRTL/PIFED/7/C80.1/3*BATAVIA//2*WBLL1/5/REH/HARE//2*BCN/3/CROC_1/AE.SQUARROSA (213)//PGO/4/HUITES/8/2*SUP152//WBLL1*2/BRAMBLING*2/3/KSW/SAUAL//S | A:A | T:T |
| BW22GS019396 | TACUPETO F2001/6/CNDO/R143//ENTE/MEXI_2/3/AEGILOPS SQUARROSA (TAUS)/4/WEAVER/5/PASTOR/7/ROLF07*2/8/SAUAL/YANAC//SAUAL/9/MOKUE #1/10/TACUPETO F2001/6/CNDO/R143//ENTE/MEXI_2/3/AEGILOPS SQUARROSA (TAUS)/4/WEAVER/5/PASTOR/7/ROLF07*2/8/SAUAL/YANAC//SAUAL | A:A | T:T |
| BW22GS019397 | TACUPETO F2001/6/CNDO/R143//ENTE/MEXI_2/3/AEGILOPS SQUARROSA (TAUS)/4/WEAVER/5/PASTOR/7/ROLF07*2/8/SAUAL/YANAC//SAUAL/9/MOKUE #1/10/TACUPETO F2001/6/CNDO/R143//ENTE/MEXI_2/3/AEGILOPS SQUARROSA (TAUS)/4/WEAVER/5/PASTOR/7/ROLF07*2/8/SAUAL/YANAC//SAUAL | A:A | T:T |
| BW22GS019398 | TACUPETO F2001/6/CNDO/R143//ENTE/MEXI_2/3/AEGILOPS SQUARROSA (TAUS)/4/WEAVER/5/PASTOR/7/ROLF07*2/8/SAUAL/YANAC//SAUAL*2/9/KASUKO | A:A | T:T |
| BW22GS019399 | TACUPETO F2001/6/CNDO/R143//ENTE/MEXI_2/3/AEGILOPS SQUARROSA (TAUS)/4/WEAVER/5/PASTOR/7/ROLF07*2/8/SAUAL/YANAC//SAUAL*2/9/KASUKO | A:A | T:T |
| BW22GS019400 | KASUKO*2/MOKUE #1 | A:A | T:T |
| BW22GS019401 | KASUKO*2/3/SUP152/QUAIU #2//BECARD/QUAIU #1 | A:A | T:T |
| BW22GS019402 | TACUPETO F2001/BRAMBLING//KACHU/8/REH/HARE//2*BCN/3/CROC_1/AE.SQUARROSA (213)//PGO/4/HUITES/5/T.DICOCCON PI94624/AE.SQUARROSA (409)//BCN/6/REH/HARE//2*BCN/3/CROC_1/AE.SQUARROSA (213)//PGO/4/HUITES/7/MUTUS*2/9/BORL14*2//KFA/2*KACHU | A:A | T:T |

| | | | |
|---|---|---|---|
| BW22GS019403 | KIRITATI/WBLL1//2*BLOUK #1*2/3/BECARD/QUAIU #1/4/SUP152//WBLL1*2/BRAMBLING*2/3/KSW/SAUAL//SAUAL/5/BECARD//ND643/2*WBLL1/3/KSW/SAUAL//SAUAL | A:A | C:C |
| BW22GS019404 | BLOUK #1/MUNAL/3/WBLL1*2/SHAMA//BAJ #1/4/SUP152/BAJ #1/5/2*SUP152//WBLL1*2/BRAMBLING*2/3/KSW/SAUAL//SAUAL | A:A | T:T |
| BW22GS019405 | BECARD/AKURI/4/WBLL1*2/BRAMBLING//JUCHI/3/WBLL1*2/BRAMBLING/5/BOKOTA/6/KASUKO/7/KASUKO | A:A | T:T |
| BW22GS019406 | KUTZ//KFA/2*KACHU/4/2*SUP152//WBLL1*2/BRAMBLING*2/3/KSW/SAUAL//SAUAL | A:A | T:T |
| BW22GS019407 | SUP152//WBLL1*2/BRAMBLING*2/3/KSW/SAUAL//SAUAL*2/4/NELOKI*2//KACHU/KIRITATI | A:A | T:T |
| BW22GS019408 | SUP152//WBLL1*2/BRAMBLING*2/3/KSW/SAUAL//SAUAL/4/BORL14*2//KFA/2*KACHU/5/BORL14*2//KFA/2*KACHU | A:A | T:T |
| BW22GS019409 | SUP152//WBLL1*2/BRAMBLING*2/3/KSW/SAUAL//SAUAL*2/4/PRL/2*PASTOR//KACHU | A:A | T:T |
| BW22GS019410 | SUP152//WBLL1*2/BRAMBLING*2/3/KSW/SAUAL//SAUAL*2/4/CIRO16/2*BORL14 | A:A | T:T |
| BW22GS019411 | SUP152//WBLL1*2/BRAMBLING*2/3/KSW/SAUAL//SAUAL*2/4/CIRO16/2*BORL14 | A:A | T:T |
| BW22GS019412 | BECARD//ND643/2*WBLL1*2/3/KSW/SAUAL//SAUAL*2/4/MOKUE #1 | A:A | T:T |
| BW22GS019413 | KINDE*2/SOLALA/3/UP2338*2/KKTS*2//YANAC/4/UP2338*2/SHAMA//2*BAJ #1*2/5/FRANCOLIN #1/3/PBW343*2/KUKUNA*2//YANAC/4/KINGBIRD #1//INQALAB 91*2/TUKURU | A:A | T:T |
| BW22GS019414 | T.DICOCCUM ABD/AE.SQUARROSA (895)/7/TRAP#1/BOW/3/VEE/PJN//2*TUI/4/BAV92/RAYON/5/KACHU #1/6/TOBA97/PASTOR/3/T.DICOCCON PI94624/AE.SQUARROSA (409)//BCN/4/BL 1496/MILAN/3/CROC_1/AE.SQUARROSA (205)//KAUZ/9/REH/HARE//2*BCN/3/CROC_1/AE.SQUARROSA (213)//PGO | A:A | T:T |
| BW22GS019415 | EARLY SPELT/AE.SQUARROSA (895)//MANKU/3/MOKUE | A:A | T:T |
| BW22GS019416 | CCB09H083/2*MOKUE #1 | A:A | T:T |
| BW22GS019417 | BERGAMO/4/WHEAR/VIVITSI//WHEAR/3/KIRITATI/2*TRCH/5/KACHU//WBLL1*2/BRAMBLING/3/KACHU/KIRITATI | A:A | T:T |
| BW22GS019418 | NADI#2/KASUKO | A:A | T:T |
| BW22GS019419 | KACHU//KIRITATI/2*TRCH/3/KASUKO | A:A | T:T |
| BW22GS019420 | CIRO16/3/TRCH/SRTU//KACHU/4/KASUKO | A:A | T:T |
| BW22GS019421 | PARUS/FRANCOLIN #1/4/MUU #1//PBW343*2/KUKUNA/3/MUU/5/KACHU*2/3/ND643//2*PRL/2*PASTOR | A:A | T:T |
| BW22GS019422 | BABAX/LR42//BABAX/3/ER2000/5/W15.92/4/PASTOR//HXL7573/2*BAU/3/WBLL1/6/SUP152//WBLL1*2/BRAMBLING*2/3/KSW/SAUAL//SAUAL | A:A | T:T |
| BW22GS019423 | SUP152//WBLL1*2/BRAMBLING*2/3/KSW/SAUAL//SAUAL/4/BORL14*2//KFA/2*KACHU | A:A | T:T |
| BW22GS019424 | MACE*2/4/KACHU*2/3/ND643//2*PRL/2*PASTOR | A:A | T:T |

| | | | |
|---|---|---|---|
| BW22GS019425 | SOKOLL/3/PASTOR//HXL7573/2*BAU*2/6/OASIS/5*BORL95/5/CNDO/R143//ENTE/MEXI75/3/AE.SQ/4/2*OCI | A:A | C:C |
| BW22GS019426 | NAINA #1 | A:A | T:T |
| BW22GS019427 | BAVIS #1//ND643/2*WBLL1/3/BORL14 | A:A | T:T |
| BW22GS019428 | AMUR*2/CHIPAK | A:A | NA |
| BW22GS019429 | MUCUY/5/PBW65/2*PASTOR/3/KIRITATI//PBW65/2*SERI.1B/4/DANPHE #1/6/MOKUE #1 | A:A | T:T |
| BW22GS019430 | MUCUY/3/KACHU//KIRITATI/2*TRCH/4/MOKUE #1 | A:A | T:T |
| BW22GS019431 | MUCUY/3/KACHU//KIRITATI/2*TRCH/4/MOKUE #1 | A:A | T:T |
| BW22GS019432 | MUCUY/3/KACHU//KIRITATI/2*TRCH/4/MOKUE #1 | NA | T:T |
| BW22GS019433 | KACHU*2/3/ND643//2*PRL/2*PASTOR*2/4/CHIPAK | A:A | T:T |
| BW22GS019434 | MELON//FILIN/MILAN/3/FILIN/4/TRCH/SRTU//KACHU/5/BORL14/6/BORL14//KFA/2*KACHU | A:A | T:T |
| BW22GS019435 | SAUAL/YANAC//SAUAL/5/UP2338*2/SHAMA/3/MILAN/KAUZ//CHIL/CHUM18/4/UP2338*2/SHAMA/6/BORL14/7/ BABAX/LR42//BABAX/3/ER2000/5/W15.92/4/PASTOR//HXL7573/2*BAU/3/WBLL1 | A:A | T:T |
| BW22GS019436 | FRANCOLIN #1/3/PBW343*2/KUKUNA*2//YANAC/4/KINGBIRD #1//INQALAB 91*2/TUKURU*2/5/BORL14 | A:A | T:T |
| BW22GS019437 | FRANCOLIN #1/3/PBW343*2/KUKUNA*2//YANAC/4/KINGBIRD #1//INQALAB 91*2/TUKURU*2/5/BORL14 | A:A | T:T |
| BW22GS019438 | FRANCOLIN #1/3/PBW343*2/KUKUNA*2//YANAC/4/KINGBIRD #1//INQALAB 91*2/TUKURU*2/5/BORL14 | A:A | T:T |
| BW22GS019439 | FRANCOLIN #1/3/PBW343*2/KUKUNA*2//YANAC/4/KINGBIRD #1//INQALAB 91*2/TUKURU*2/5/MUNAL #1 | A:A | T:T |
| BW22GS019440 | FRANCOLIN #1/3/PBW343*2/KUKUNA*2//YANAC/4/KINGBIRD #1//INQALAB 91*2/TUKURU*2/5/KINGBIRD #1//INQALAB 91*2/TUKURU | A:A | T:T |
| BW22GS019441 | FRANCOLIN #1/3/PBW343*2/KUKUNA*2//YANAC/4/KINGBIRD #1//INQALAB 91*2/TUKURU*2/5/KINGBIRD #1//INQALAB 91*2/TUKURU | A:A | T:T |
| BW22GS019442 | FRANCOLIN #1/3/PBW343*2/KUKUNA*2//YANAC/4/KINGBIRD #1//INQALAB 91*2/TUKURU*2/5/NADI#2 | A:A | C:T |
| BW22GS019443 | FRANCOLIN #1/3/PBW343*2/KUKUNA*2//YANAC/4/KINGBIRD #1//INQALAB 91*2/TUKURU*2/5/MUCUY | A:A | T:T |
| BW22GS019444 | FRANCOLIN #1/3/PBW343*2/KUKUNA*2//YANAC/4/KINGBIRD #1//INQALAB 91*2/TUKURU*2/5/BECARD/CHYAK | A:A | T:T |
| BW22GS019445 | KACHU/SAUAL/4/VARIS/MISR 2/3/FRET2/KUKUNA//FRET2/5/KACHU/SAUAL/6/BORL14/7/KASUKO | A:A | T:T |
| BW22GS019446 | PASTOR//HXL7573/2*BAU/3/SOKOLL/WBLL1/6/2*OASIS/5*BORL95/5/CNDO/R143//ENTE/MEXI75/3/AE.SQ/4/2* OCI*2/7/NADI#2 | A:A | T:T |

| | | | |
|---|---|---|---|
| BW22GS019447 | PASTOR//HXL7573/2*BAU/3/SOKOLL/WBLL1/6/2*OASIS/5*BORL95/5/CNDO/R143//ENTE/MEXI75/3/AE.SQ/4/2*OCI*2/7/NADI#2 | A:A | T:T |
| BW22GS019448 | CHIPAK/MOKUE #1 | A:A | T:T |
| BW22GS019449 | MUCUY/3/KFA/2*KACHU*2//SUP152 | A:A | T:T |
| BW22GS019450 | BLOUK #1/4/WHEAR/KUKUNA/3/C80.1/3*BATAVIA//2*WBLL1/5/MUNAL #1/6/KASUKO | A:A | T:T |
| BW22GS019451 | BLOUK #1/4/WHEAR/KUKUNA/3/C80.1/3*BATAVIA//2*WBLL1/5/MUNAL #1/6/KASUKO | A:A | T:T |
| BW22GS019452 | SUP152/FRNCLN/5/FRANCOLIN #1/3/PBW343*2/KUKUNA*2//YANAC/4/KINGBIRD #1//INQALAB 91*2/TUKURU | A:A | T:T |
| BW22GS019453 | SUP152/FRNCLN//KASUKO | A:A | T:T |
| BW22GS019454 | MUNAL*2/WESTONIA//KASUKO | A:A | T:T |
| BW22GS019455 | MUNAL*2/WESTONIA/3/KUTZ//KFA/2*KACHU | A:A | T:T |
| BW22GS019456 | MUNAL*2/WESTONIA/3/KUTZ//KFA/2*KACHU | A:A | C:C |
| BW22GS019457 | SWSR22T.B./KACHU//2*KACHU/8/ATTILA/3*BCN//BAV92/3/TILHI/4/SUP152/5/SUP152/6/KFA/2*KACHU/7/ATTILA/3*BCN//BAV92/3/PASTOR/4/TACUPETO F2001*2/BRAMBLING/5/PAURAQ | A:A | T:T |
| BW22GS019458 | SHORTENED SR26 TRANSLOCATION//2*WBLL1*2/KKTS/3/BECARD/4/SWSR22T.B.//TACUPETO F2001*2/BRAMBLING/3/2*TACUPETO F2001*2/BRAMBLING | A:A | T:T |
| BW22GS019459 | SHORTENED SR26 TRANSLOCATION//2*WBLL1*2/KKTS/3/BECARD/4/SWSR22T.B.//TACUPETO F2001*2/BRAMBLING/3/2*TACUPETO F2001*2/BRAMBLING | A:A | T:T |
| BW22GS019460 | SHORTENED SR26 TRANSLOCATION//2*WBLL1*2/KKTS/3/BECARD/4/KASUKO | A:A | T:T |
| BW22GS019461 | SWSR22T.B.//TACUPETO F2001*2/BRAMBLING/3/2*TACUPETO F2001*2/BRAMBLING/4/KASUKO | A:A | T:T |
| BW22GS019462 | BAJ #1*2/KISKADEE #1//KASUKO | A:A | T:T |
| BW22GS019463 | BAVIS//ATTILA*2/PBW65/3/KASUKO | A:A | T:T |
| BW22GS019464 | BAVIS//ATTILA*2/PBW65/3/KASUKO | A:A | T:T |
| BW22GS019465 | GLADIUS/3/2*KA/NAC//TRCH/4/KUTZ//KFA/2*KACHU | A:A | T:T |
| BW22GS019466 | GLADIUS/3/2*KA/NAC//TRCH/4/KUTZ//KFA/2*KACHU | A:A | T:T |
| BW22GS019467 | VENDA/4/SUP152//WBLL1*2/BRAMBLING*2/3/KSW/SAUAL//SAUAL | A:A | C:T |
| BW22GS019468 | BECARD//ND643/2*WBLL1/3/SWSR22T.B./2*BLOUK #1//WBLL1*2/KURUKU | A:A | T:T |
| BW22GS019469 | BECARD//ND643/2*WBLL1/3/SWSR22T.B./2*BLOUK #1//WBLL1*2/KURUKU | A:A | T:T |
| BW22GS019470 | KACHU*2/3/ND643//2*PRL/2*PASTOR/4/KASUKO | A:A | T:T |
| BW22GS019471 | KACHU*2/3/ND643//2*PRL/2*PASTOR/4/MOKUE #1 | A:A | T:T |
| BW22GS019472 | KACHU*2/3/ND643//2*PRL/2*PASTOR/4/MOKUE #1 | A:A | T:T |
| BW22GS019473 | KACHU*2/3/ND643//2*PRL/2*PASTOR/4/MOKUE #1 | A:A | T:T |
| BW22GS019474 | KACHU*2/3/ND643//2*PRL/2*PASTOR/4/MOKUE #1 | A:A | T:T |

| | | | |
|---|---|---|---|
| BW22GS019475 | KACHU*2/3/ND643//2*PRL/2*PASTOR/4/MOKUE #1 | A:A | T:T |
| BW22GS019476 | KACHU*2/3/ND643//2*PRL/2*PASTOR/4/MOKUE #1 | A:A | T:T |
| BW22GS019477 | KACHU*2/3/ND643//2*PRL/2*PASTOR/4/MOKUE #1 | A:A | T:T |
| BW22GS019478 | KACHU*2/3/ND643//2*PRL/2*PASTOR/4/MOKUE #1 | A:A | T:T |
| BW22GS019479 | PRL/2*PASTOR//KACHU/4/KACHU*2/3/ND643//2*PRL/2*PASTOR | A:A | T:T |
| BW22GS019480 | PRL/2*PASTOR//KACHU/3/MOKUE #1 | A:A | T:T |
| BW22GS019481 | PRL/2*PASTOR//KACHU/3/MOKUE #1 | A:A | T:T |
| BW22GS019482 | PRL/2*PASTOR//KACHU/3/MOKUE #1 | A:A | T:T |
| BW22GS019483 | PRL/2*PASTOR//KACHU/3/KASUKO | A:A | T:T |
| BW22GS019484 | PRL/2*PASTOR//KACHU/3/BORL14*2//KFA/2*KACHU | A:A | T:T |
| BW22GS019485 | FRANCOLIN #1/BAJ #1//MOKUE #1 | A:A | T:T |
| BW22GS019486 | SUP152/CIRO16//KASUKO | A:A | T:T |
| BW22GS019487 | COPIO/5/UP2338*2/SHAMA/3/MILAN/KAUZ//CHIL/CHUM18/4/UP2338*2/SHAMA/6/KSW/SAUAL//SAUAL/3/TRCH/HUIRIVIS #1/5/UP2338*2/SHAMA/3/MILAN/KAUZ//CHIL/CHUM18/4/UP2338*2/SHAMA | A:A | T:T |
| BW22GS019488 | SHA7//PRL/VEE#6/3/FASAN/4/HAAS8446/2*FASAN/5/CBRD/KAUZ/6/MILAN/AMSEL/7/FRET2*2/KUKUNA/8/TRAP #1/BOW/3/VEE/PJN//2*TUI/4/BAV92/RAYON/5/KACHU #1/9/COPIO/10/SHORTENED SR26 TRANSLOCATION//2*WBLL1*2/KKTS/3/BECARD | A:A | T:T |
| BW22GS019489 | WBLL1*2/4/YACO/PBW65/3/KAUZ*2/TRAP//KAUZ/5/KACHU #1*2/6/FRET2/KUKUNA//FRET2/3/TNMU/4/FRET2*2/SHAMA/7/KASUKO | A:A | T:T |
| BW22GS019490 | WBLL1*2/4/YACO/PBW65/3/KAUZ*2/TRAP//KAUZ/5/KACHU #1*2/6/FRET2/KUKUNA//FRET2/3/TNMU/4/FRET2*2/SHAMA/8/CNO79//PF70354/MUS/3/PASTOR/4/BAV92*2/5/HAR311/6/PBW343*2/KUKUNA*2//FRTL/PIFED/7/CNO79//PF70354/MUS/3/PASTOR/4/BAV92*2/5/HAR311 | A:A | T:T |
| BW22GS019491 | WBLL1*2/4/YACO/PBW65/3/KAUZ*2/TRAP//KAUZ/5/KACHU #1*2/6/FRET2/KUKUNA//FRET2/3/TNMU/4/FRET2*2/SHAMA/8/CNO79//PF70354/MUS/3/PASTOR/4/BAV92*2/5/HAR311/6/PBW343*2/KUKUNA*2//FRTL/PIFED/7/CNO79//PF70354/MUS/3/PASTOR/4/BAV92*2/5/HAR311 | A:A | T:T |
| BW22GS019492 | WBLL1*2/4/YACO/PBW65/3/KAUZ*2/TRAP//KAUZ/5/KACHU #1*2/6/FRET2/KUKUNA//FRET2/3/TNMU/4/FRET2*2/SHAMA/8/CNO79//PF70354/MUS/3/PASTOR/4/BAV92*2/5/HAR311/6/PBW343*2/KUKUNA*2//FRTL/PIFED/7/CNO79//PF70354/MUS/3/PASTOR/4/BAV92*2/5/HAR311 | A:A | T:T |
| BW22GS019493 | TACUPETO F2001/BRAMBLING/5/NAC/TH.AC//3*PVN/3/MIRLO/BUC/4/2*PASTOR*2/6/TRCH/SRTU//KACHU/7/PRL/2*PASTOR//KACHU | A:A | T:T |

| | | | |
|---|---|---|---|
| BW22GS019494 | TACUPETO F2001/BRAMBLING/5/NAC/TH.AC//3*PVN/3/MIRLO/BUC/4/2*PASTOR*2/6/WAXWING/SRTU//WAXWING/KIRITATI/7/KUTZ//KFA/2*KACHU | A:A | T:T |
| BW22GS019495 | KACHU/SAUAL*2/3/TACUPETO F2001/BRAMBLING//KIRITATI/4/KACHU*2/3/ND643//2*PRL/2*PASTOR | A:A | T:T |
| BW22GS019496 | KACHU/SAUAL*2/3/TACUPETO F2001/BRAMBLING//KIRITATI/4/BORL14*2//KFA/2*KACHU | A:A | T:T |
| BW22GS019497 | KACHU/SAUAL*2/4/ATTILA*2/PBW65//PIHA/3/ATTILA/2*PASTOR/5/BORL14*2//KFA/2*KACHU | A:A | T:T |
| BW22GS019498 | SAUAL/MUTUS//KINGBIRD #1/3/SAUAL/MUTUS/4/KASUKO | A:A | T:T |
| BW22GS019499 | SAUAL/MUTUS//KINGBIRD #1/3/SAUAL/MUTUS/4/KASUKO | A:A | T:T |
| BW22GS019500 | SAUAL/MUTUS//KINGBIRD #1/3/SAUAL/MUTUS/4/BORL14*2//KFA/2*KACHU | A:A | T:T |
| BW22GS019501 | FRET2/KUKUNA//FRET2/3/PARUS/4/FRET2*2/SHAMA*2/5/WBLL1/KUKUNA//TACUPETO F2001/3/UP2338*2/VIVITSI/6/KASUKO | A:A | T:T |
| BW22GS019502 | FRET2/KUKUNA//FRET2/3/PARUS/4/FRET2*2/SHAMA*2/5/WBLL1/KUKUNA//TACUPETO F2001/3/UP2338*2/VIVITSI/6/KASUKO | A:A | T:T |
| BW22GS019503 | FRET2/KUKUNA//FRET2/3/PARUS/4/FRET2*2/SHAMA*2/5/WBLL1/KUKUNA//TACUPETO F2001/3/UP2338*2/VIVITSI/6/KASUKO | A:A | T:T |
| BW22GS019504 | PBW343*2/KUKUNA*2//KITE/3/ATTILA*2/PBW65*2//YANAC/4/SAUAL/YANAC//SAUAL/6/WAXWING/KIRITATI*2/3/C80.1/3*BATAVIA//2*WBLL1/4/COPIO/5/ND643//2*ATTILA*2/PASTOR/3/WBLL1*2/KURUKU/4/WBLL1*2/BRAMBLING | A:A | T:T |
| BW22GS019505 | PARUS/FRANCOLIN #1/4/MUU #1//PBW343*2/KUKUNA/3/MUU/5/KASUKO | A:A | T:T |
| BW22GS019506 | WBLL1*2/KKTS//PASTOR/KUKUNA/3/KINGBIRD #1//INQALAB 91*2/TUKURU/5/KAUZ//ALTAR 84/AOS/3/MILAN/KAUZ/4/SAUAL/6/MOKUE #1 | A:A | T:T |
| BW22GS019507 | AMUR*2/CIRO16//KASUKO | A:A | T:T |
| BW22GS019508 | ROLF07/YANAC//TACUPETO F2001/BRAMBLING*2/5/UP2338*2/SHAMA/3/MILAN/KAUZ//CHIL/CHUM18/4/UP2338*2/SHAMA/6/BORL14 | A:A | T:T |
| BW22GS019509 | FRET2/KUKUNA//FRET2/3/YANAC/4/FRET2/KIRITATI/5/2*UP2338*2/SHAMA/3/MILAN/KAUZ//CHIL/CHUM18/4/UP2338*2/SHAMA/6/BORL14*2//KFA/2*KACHU | A:A | T:T |
| BW22GS019510 | WBLL1/3/STAR//KAUZ/STAR/4/BAV92/RAYON/5/TRAP#1/BOW/3/VEE/PJN//2*TUI/4/BAV92/RAYON*2/8/TACUPETO F2001/6/CNDO/R143//ENTE/MEXI_2/3/AEGILOPS SQUARROSA (TAUS)/4/WEAVER/5/PASTOR/7/ROLF07/9/SUP152//WBLL1*2/BRAMBLING*2/3/KSW/SAUAL//SAUAL | A:A | T:T |
| BW22GS019511 | BECARD/AKURI*2/3/PBW343*2/KUKUNA*2//FRTL/PIFED/4/SUP152//WBLL1*2/BRAMBLING*2/3/KSW/SAUAL//SAUAL | A:A | T:T |
| BW22GS019512 | TACUPETO F2001/6/CNDO/R143//ENTE/MEXI_2/3/AEGILOPS SQUARROSA (TAUS)/4/WEAVER/5/PASTOR/7/ROLF07*2/8/SAUAL/YANAC//SAUAL/9/KASUKO | A:A | T:T |

| | | | |
|---|---|---|---|
| BW22GS019513 | WADER/4/KACHU//WBLL1*2/BRAMBLING*2/3/KACHU/KIRITATI | A:A | T:T |
| BW22GS019514 | WADER/4/KACHU//WBLL1*2/BRAMBLING*2/3/KACHU/KIRITATI | A:A | T:T |
| BW22GS019515 | BECARD/AKURI/4/WBLL1*2/BRAMBLING//JUCHI/3/WBLL1*2/BRAMBLING/5/KUTZ//KFA/2*KACHU | A:A | C:C |
| BW22GS019516 | MOKUE #1/3/BORL14*2//KFA/2*KACHU | A:A | T:T |
| BW22GS019517 | MOKUE #1/3/BORL14*2//KFA/2*KACHU | A:A | T:T |
| BW22GS019518 | MOKUE #1/3/BORL14*2//KFA/2*KACHU | A:A | C:C |
| BW22GS019519 | MOKUE #1/3/KACHU/BECARD//WBLL1*2/BRAMBLING | A:A | C:T |
| BW22GS019520 | SITE/MO//PASTOR/3/TILHI/4/MUNAL #1/5/MUNAL/6/MUCUY/7/MOKUE #1 | A:A | T:T |
| BW22GS019521 | QUAIU #1/5/KIRITATI/4/2*BAV92//IRENA/KAUZ/3/HUITES/6/BECARD/QUAIU #1/7/SUP152//WBLL1*2/BRAMBLING*2/3/KSW/SAUAL//SAUAL | A:A | C:T |
| BW22GS019522 | NADI#1/3/PBW343*2/KUKUNA*2//FRTL/PIFED/4/NADI#2/5/FRANCOLIN #1/3/PBW343*2/KUKUNA*2//YANAC/4/KINGBIRD #1//INQALAB 91*2/TUKURU | A:A | T:T |
| BW22GS019523 | NAINA #2/3/PRL/2*PASTOR//KACHU | A:A | T:T |
| BW22GS019524 | KFA/2*KACHU/4/WBLL1*2/KURUKU//KRONSTAD F2004/3/WBLL1*2/BRAMBLING/5/KUTZ//KFA/2*KACHU | A:A | NA |
| BW22GS019525 | TACUPETO F2001/BRAMBLING//KIRITATI/3/FRANCOLIN #1/BLOUK #1/4/FRANCOLIN #1/BLOUK #1/5/SHORTENED SR26 TRANSLOCATION//2*WBLL1*2/KKTS/3/BECARD | A:A | T:T |
| BW22GS019526 | KASUKO/3/SUP152/QUAIU #2//BECARD/QUAIU #1 | A:A | T:T |
| BW22GS019527 | KASUKO/3/SUP152/QUAIU #2//BECARD/QUAIU #1 | A:A | T:T |
| BW22GS019528 | BOKOTA/5/UP2338*2/VIVITSI/3/FRET2/TUKURU//FRET2/4/MISR 1/6/BABAX/LR42//BABAX*2/3/KUKUNA/4/CROSBILL #1/5/BECARD/7/BORL14//KFA/2*KACHU | A:A | T:T |
| BW22GS019529 | WBLL1*2/BRAMBLING//JUCHI/3/KINGBIRD #1//INQALAB 91*2/TUKURU/4/WBLL1*2/BRAMBLING//KACHU/5/KASUKO | A:A | T:T |
| BW22GS019530 | BABAX/LR42//BABAX/3/ER2000/5/BABAX/LR39//BABAX*2/4/KABY/BAV92/3/CROC_1/AE.SQUARROSA (224)//OPATA/6/SUP152//WBLL1*2/BRAMBLING*2/3/KSW/SAUAL//SAUAL | A:A | T:T |
| BW22GS019531 | TACUPETO F2001/BRAMBLING//KACHU/8/REH/HARE//2*BCN/3/CROC_1/AE.SQUARROSA (213)//PGO/4/HUITES/5/T.DICOCCON PI94624/AE.SQUARROSA (409)//BCN/6/REH/HARE//2*BCN/3/CROC_1/AE.SQUARROSA (213)//PGO/4/HUITES/7/MUTUS/9/BORL14*2//KFA/2*KACHU | A:A | T:T |
| BW22GS019532 | FRANCOLIN #1//WBLL1*2/KURUKU/3/WBLL1*2/BRAMBLING//CHYAK/4/SUP152//WBLL1*2/BRAMBLING*2/3/KSW/SAUAL//SAUAL | A:A | C:C |
| BW22GS019533 | SUP152/BAJ #1/3/KINGBIRD #1//INQALAB 91*2/TUKURU/8/ATTILA/3*BCN//BAV92/3/TILHI/4/SUP152/5/SUP152/6/KFA/2*KACHU/7/ATTILA/3*BCN//BAV92/3/PASTOR/4/TACUPETO F2001*2/BRAMBLING/5/PAURAQ | A:A | T:T |

| | | | |
|---|---|---|---|
| BW22GS019534 | SUP152/BAJ #1/3/KINGBIRD #1//INQALAB 91*2/TUKURU/8/ATTILA/3*BCN//BAV92/3/TILHI/4/SUP152/5/SUP152/6/KFA/2*KACHU/7/ATTILA/3*BCN//BAV92 /3/PASTOR/4/TACUPETO F2001*2/BRAMBLING/5/PAURAQ | A:A | T:T |
| BW22GS019535 | KACHU/BECARD//WBLL1*2/BRAMBLING/3/KACHU/KINDE/4/KASUKO | A:A | T:T |
| BW22GS019536 | PREMIO/4/CROC_1/AE.SQUARROSA (205)//KAUZ/3/PIFED/5/VORB/FISCAL//KACHU/3/WBLL1*2/BRAMBLING/6/KASUKO | NA | T:T |
| BW22GS019537 | PREMIO/4/CROC_1/AE.SQUARROSA (205)//KAUZ/3/PIFED/5/VORB/FISCAL//KACHU/3/WBLL1*2/BRAMBLING/6/KASUKO | A:A | T:T |
| BW22GS019538 | PREMIO/4/CROC_1/AE.SQUARROSA (205)//KAUZ/3/PIFED/5/VORB/FISCAL//KACHU/3/WBLL1*2/BRAMBLING/6/KASUKO | A:A | T:T |
| BW22GS019539 | BORL14*2//MUNAL #1/FRANCOLIN #1/4/KACHU*2/3/ND643//2*PRL/2*PASTOR | A:A | T:T |
| BW22GS019540 | KACHU//WBLL1*2/BRAMBLING*2/6/ROLF07*2/5/REH/HARE//2*BCN/3/CROC_1/AE.SQUARROSA (213)//PGO/4/HUITES/7/KUTZ*2//KFA/2*KACHU | A:A | T:T |
| BW22GS019541 | CIANO M2018/4/KACHU//WBLL1*2/BRAMBLING*2/3/KACHU/KIRITATI | A:A | T:T |
| BW22GS019542 | BLOUK #1/MUNAL/3/WBLL1*2/SHAMA//BAJ #1/4/SUP152/BAJ #1/5/MOKUE #1 | A:A | T:T |
| BW22GS019543 | BECARD/AKURI/3/KACHU//WBLL1*2/BRAMBLING/4/MUTUS/AKURI/5/MOKUE #1 | A:A | T:T |
| BW22GS019544 | BECARD/AKURI/3/KACHU//WBLL1*2/BRAMBLING/4/MUTUS/AKURI/5/MOKUE #1 | A:A | T:T |
| BW22GS019545 | BECARD/AKURI/3/KACHU//WBLL1*2/BRAMBLING/4/MUTUS/AKURI/5/MOKUE #1 | A:A | T:T |
| BW22GS019546 | ATTILA/3*BCN//BAV92/3/PASTOR/4/MUNAL #1/5/MUNAL/6/2*BECARD/QUAIU #1/7/KFA/2*KACHU*2//SUP152 | A:A | T:T |
| BW22GS019547 | SUP152/BAJ #1/3/KACHU//WBLL1*2/BRAMBLING/6/KSW/SAUAL//SAUAL/3/TRCH/HUIRIVIS #1/5/UP2338*2/SHAMA/3/MILAN/KAUZ//CHIL/CHUM18/4/UP2338*2/SHAMA | A:A | T:T |
| BW22GS019548 | CHIPAK*2//KFA/2*KACHU/3/KUTZ//KFA/2*KACHU | A:A | T:T |
| BW22GS019549 | CHIPAK*2//KFA/2*KACHU/3/KUTZ//KFA/2*KACHU | A:A | T:T |
| BW22GS019550 | CHIPAK*2//KFA/2*KACHU/7/ATTILA/3*BCN//BAV92/3/PASTOR/4/TACUPETO F2001*2/BRAMBLING/5/PAURAQ/6/KFA/2*KACHU | A:A | T:T |
| BW22GS019551 | TUKURU//BAV92/RAYON/3/MUNAL #1/4/2*KFA/2*KACHU/5/BORL14*2//KFA/2*KACHU | A:A | T:T |
| BW22GS019552 | KUTZ//KFA/2*KACHU/3/KASUKO | A:A | T:T |
| BW22GS019553 | BORL14*2//KFA/2*KACHU/3/KASUKO | A:A | T:T |
| BW22GS019554 | ROLF07*2/DIAMONDBIRD//TRCH/HUIRIVIS #1/3/BORL14/4/NELOKI*2//KACHU/KIRITATI | A:A | T:T |
| BW22GS019555 | ROLF07*2/DIAMONDBIRD//TRCH/HUIRIVIS #1/3/BORL14/4/NELOKI*2//KACHU/KIRITATI | A:A | T:T |
| BW22GS019556 | FRANCOLIN #1*2/HAWFINCH #1//2*MUCUY/4/MUTUS*2/KINGBIRD #1/3/KSW/SAUAL//SAUAL | A:A | T:T |
| BW22GS019557 | FRANCOLIN #1*2/HAWFINCH #1//2*MUCUY/4/MUTUS*2/KINGBIRD #1/3/KSW/SAUAL//SAUAL | A:A | T:T |
| BW22GS019558 | FRANCOLIN #1*2/HAWFINCH #1//2*MUCUY/4/MUTUS*2/KINGBIRD #1/3/KSW/SAUAL//SAUAL | A:A | T:T |

| | | | |
|---|---|---|---|
| BW22GS019559 | TACUPETO F2001/6/CNDO/R143//ENTE/MEXI_2/3/AEGILOPS SQUARROSA (TAUS)/4/WEAVER/5/PASTOR/7/ROLF07/8/PBW343*2/KUKUNA*2//FRTL/PIFED/9/KUTZ*2//KFA/2*KACHU | A:A | T:T |
| BW22GS019560 | ND643/2*TRCH//BECARD/3/BECARD/4/SUP152*2/TECUE #1/5/KUTZ*2//KFA/2*KACHU | A:A | T:T |
| BW22GS019561 | MUNAL/WESTONIA//SUP152/BAJ #1/3/MOKUE #1 | A:A | T:T |
| BW22GS019562 | BORL14/MOKUE #1//KASUKO | A:A | C:T |
| BW22GS019563 | MUCUY/4/SUP152//WBLL1*2/BRAMBLING*2/3/KSW/SAUAL//SAUAL/5/MOKUE #1 | A:A | T:T |
| BW22GS019564 | QUAIU #1/SUP152/3/BAJ #1/TECUE #1//MUTUS*2/TECUE #1/4/CHIPAK*2/3/KSW/SAUAL//SAUAL | A:A | T:T |
| BW22GS019565 | PRL/2*PASTOR//PBW343*2/KUKUNA/3/ROLF07/4/BERKUT//PBW343*2/KUKUNA/5/KASUKO/6/BORL14//KFA/2*KACHU | A:A | T:T |
| BW22GS019566 | MUTUS//ND643/2*WBLL1/4/SHORTENED SR26 TRANSLOCATION//2*WBLL1*2/KKTS/3/BECARD/5/MOKUE #1 | A:A | T:T |
| BW22GS019567 | CHIBIA//PRLII/CM65531/3/MISR 2*2/4/HUW234+LR34/PRINIA//PBW343*2/KUKUNA/3/ROLF07/5/BORL14*2//KFA/2*KACHU/6/BORL14*2//KFA/2*KACHU | A:A | T:T |
| BW22GS019568 | MELON//FILIN/MILAN/3/FILIN/4/TRCH/SRTU//KACHU/5/2*KACHU//WBLL1*2/BRAMBLING/3/KACHU/KIRITATI | A:A | T:T |
| BW22GS019569 | MELON//FILIN/MILAN/3/FILIN/4/TRCH/SRTU//KACHU/5/2*KACHU//WBLL1*2/BRAMBLING/3/KACHU/KIRITATI | A:A | T:T |
| BW22GS019570 | KACHU/3/WHEAR//2*PRL/2*PASTOR/4/KASUKO/5/PRL/2*PASTOR//KACHU | A:A | T:T |
| BW22GS019571 | KACHU/3/WHEAR//2*PRL/2*PASTOR/4/KASUKO/5/PRL/2*PASTOR//KACHU | A:A | T:T |
| BW22GS019572 | BAV92//IRENA/KAUZ/3/HUITES/4/PVN/5/CIRO16/6/2*MOKUE #1 | A:A | T:T |
| BW22GS019573 | CIRO16/3/TRCH/SRTU//KACHU/4/KUTZ//KFA/2*KACHU/5/KUTZ//KFA/2*KACHU | A:A | T:T |
| BW22GS019574 | KACHU/3/WHEAR//2*PRL/2*PASTOR/4/BOKOTA*2/5/MOKUE #1 | A:A | T:T |
| BW22GS019575 | FRANCOLIN #1/3/PBW343*2/KUKUNA*2//YANAC/4/KINGBIRD #1//INQALAB 91*2/TUKURU*2/5/BORL14 | A:A | T:T |
| BW22GS019576 | TACUPETO F2001/BRAMBLING/5/NAC/TH.AC//3*PVN/3/MIRLO/BUC/4/2*PASTOR*2/6/WAXWING/SRTU//WAXWING/KIRITATI/7/KASUKO/8/ROLF07*2/DIAMONDBIRD//TRCH/HUIRIVIS #1/3/BORL14 | A:A | T:T |
| BW22GS019577 | SAUAL/MUTUS//KINGBIRD #1/3/SAUAL/MUTUS/4/BORL14*2//KFA/2*KACHU/5/BORL14*2//KFA/2*KACHU | A:A | T:T |
| BW22GS019578 | SAUAL/MUTUS//KINGBIRD #1/3/SAUAL/MUTUS/4/KUTZ//KFA/2*KACHU/5/KUTZ*2//KFA/2*KACHU | A:A | T:T |
| BW22GS019579 | SAUAL/MUTUS*2//CIRO16*2/3/MOKUE #1 | A:A | T:T |
| BW22GS019580 | SAUAL/MUTUS*2//CIRO16*2/3/MOKUE #1 | A:A | T:T |

| | | | |
|---|---|---|---|
| BW22GS019581 | SAUAL/MUTUS*2//CIRO16*2/3/MOKUE #1 | A:A | T:T |
| BW22GS019582 | FRET2/KUKUNA//FRET2/3/PARUS/4/FRET2*2/SHAMA*2/5/WBLL1/KUKUNA//TACUPETO F2001/3/UP2338*2/VIVITSI/6/PRL/2*PASTOR//KACHU/7/KFA/2*KACHU*2//SUP152 | A:A | T:T |
| BW22GS019583 | CROC_1/AE.SQUARROSA (205)//BORL95/3/PRL/SARA//TSI/VEE#5/4/FRET2/6/MTRWA92.161/PRINIA/5/SERI*3//RL6010/4*YR/3/PASTOR/4/BAV92/7/BORL14//KFA/2*KACHU/8/BORL14*2//KFA/2*KACHU | A:A | T:T |
| BW22GS019584 | CROC_1/AE.SQUARROSA (205)//BORL95/3/PRL/SARA//TSI/VEE#5/4/FRET2/6/MTRWA92.161/PRINIA/5/SERI*3//RL6010/4*YR/3/PASTOR/4/BAV92/7/2*KACHU//WBLL1*2/BRAMBLING/3/KACHU/KIRITATI | A:A | T:T |
| BW22GS019585 | QUAIU #2/BAVIS #1//KAKURU/4/SHORTENED SR26 TRANSLOCATION//2*WBLL1*2/KKTS/3/BECARD | A:A | T:T |
| BW22GS019586 | FRET2*2/SHAMA//PARUS/3/FRET2*2/KUKUNA/4/WBLL1/KUKUNA//TACUPETO F2001/3/UP2338*2/VIVITSI/5/PRL/2*PASTOR//KACHU/6/KACHU//WBLL1*2/BRAMBLING*2/3/KACHU/KIRITATI | A:A | T:T |
| BW22GS019587 | FRET2*2/SHAMA//PARUS/3/FRET2*2/KUKUNA/4/WBLL1/KUKUNA//TACUPETO F2001/3/UP2338*2/VIVITSI/5/PRL/2*PASTOR//KACHU/6/KACHU//WBLL1*2/BRAMBLING*2/3/KACHU/KIRITATI | A:A | T:T |
| BW22GS019588 | AMUR*2/CIRO16*2/3/KFA/2*KACHU*2//SUP152 | A:A | T:T |
| BW22GS019589 | WADER #2/4/2*SHORTENED SR26 TRANSLOCATION//2*WBLL1*2/KKTS/3/BECARD | A:A | T:T |
| BW22GS019590 | WADER #2/4/2*SHORTENED SR26 TRANSLOCATION//2*WBLL1*2/KKTS/3/BECARD | A:A | T:T |
| BW22GS019591 | KUTZ//KFA/2*KACHU/3/CHIPAK/4/CHIPAK*2/3/KSW/SAUAL//SAUAL | A:A | T:T |
| BW22GS019592 | MOKUE #1/3/BORL14*2//KFA/2*KACHU/4/BORL14*2//KFA/2*KACHU | A:A | T:T |
| BW22GS019593 | MOKUE #1/3/BORL14*2//KFA/2*KACHU/4/BORL14*2//KFA/2*KACHU | A:A | T:T |
| BW22GS019594 | MOKUE #1*2/NADI | A:A | T:T |
| BW22GS019595 | MOKUE #1*2/NADI | A:A | T:T |
| BW22GS019596 | MOKUE #1*2/NADI | A:A | T:T |
| BW22GS019597 | MOKUE #1/6/WBLL1*2/KKTS//PASTOR/KUKUNA/3/KINGBIRD #1//INQALAB 91*2/TUKURU/5/KAUZ//ALTAR 84/AOS/3/MILAN/KAUZ/4/SAUAL/7/BORL14*2//BECARD/QUAIU #1 | A:A | T:T |
| BW22GS019598 | NADI#1/3/PBW343*2/KUKUNA*2//FRTL/PIFED/4/NADI#2*2/5/KASUKO | A:A | T:T |
| BW22GS019599 | NAINA #3/3/NELOKI*2//KACHU/KIRITATI/4/NAINA #2 | A:A | T:T |
| BW22GS019600 | WAXWING/4/BL 1496/MILAN/3/CROC_1/AE.SQUARROSA (205)//KAUZ/5/FRNCLN/6/KINGBIRD #1//INQALAB 91*2/TUKURU/7/BECARD/QUAIU #1/8/2*BORL14*2//KFA/2*KACHU | A:A | T:T |
| BW22GS019601 | WAXWING/4/BL 1496/MILAN/3/CROC_1/AE.SQUARROSA (205)//KAUZ/5/FRNCLN/6/KINGBIRD #1//INQALAB 91*2/TUKURU/7/BECARD/QUAIU #1/8/2*KACHU//WBLL1*2/BRAMBLING*2/3/KACHU/KIRITATI | A:A | T:T |

| | | | |
|---|---|---|---|
| BW22GS019602 | W15.92/4/PASTOR//HXL7573/2*BAU/3/WBLL1/7/CNO79//PF70354/MUS/3/PASTOR/4/BAV92/5/FRET2/KUKUNA//FRET2/6/MILAN/KAUZ//PRINIA/3/BAV92*2/8/MOKUE #1 | A:A | T:T |
| BW22GS019603 | KFA/2*KACHU/4/WBLL1*2/KURUKU//KRONSTAD F2004/3/WBLL1*2/BRAMBLING/5/KUTZ//KFA/2*KACHU/6/KUTZ//KFA/2*KACHU | A:A | T:T |
| BW22GS019604 | BAJ #1*2/PREMIO//2*MOKUE #1 | A:A | T:T |
| BW22GS019605 | PBW343*2/KUKUNA*2//FRTL/PIFED/3/ABLEU*2/4/MOKUE #1 | A:A | T:T |
| BW22GS019606 | PBW343*2/KUKUNA*2//FRTL/PIFED/3/ABLEU*2/4/MOKUE #1 | A:A | T:T |
| BW22GS019607 | SUP152/BAJ #1/3/KINGBIRD #1//INQALAB 91*2/TUKURU*2/4/MOKUE #1 | A:A | T:T |
| BW22GS019608 | SUP152/BAJ #1/3/KINGBIRD #1//INQALAB 91*2/TUKURU*2/4/MOKUE #1 | A:A | T:T |
| BW22GS019609 | SUP152/BAJ #1/3/KINGBIRD #1//INQALAB 91*2/TUKURU*2/4/MOKUE #1 | A:A | T:T |
| BW22GS019610 | SUP152/BAJ #1/3/KINGBIRD #1//INQALAB 91*2/TUKURU*2/4/MOKUE #1 | A:A | T:T |
| BW22GS019611 | CIRO16/2*BORL14//MOKUE #1/3/BORL14*2//KFA/2*KACHU | A:A | T:T |
| BW22GS019612 | BORL14*2/MUNAL #1*2/3/SWSR22T.B./2*BLOUK #1//WBLL1*2/KURUKU | A:A | T:T |
| BW22GS019613 | BORL14*2/MUNAL #1/4/SHORTENED SR26 TRANSLOCATION//2*WBLL1*2/KKTS/3/BECARD/5/BORL14*2/MUNAL #1 | A:A | T:T |
| BW22GS019614 | BORL14*2/3/KBIRD//WBLL1*2/KURUKU/4/SUP152//WBLL1*2/BRAMBLING*2/3/KSW/SAUAL//SAUAL/5/ROLF07*2/DIAMONDBIRD//TRCH/HUIRIVIS #1/3/BORL14 | A:A | T:T |
| BW22GS019615 | BORL14*2//KFA/2*KACHU/3/MOKUE #1/4/BORL14*2//KFA/2*KACHU | A:A | T:T |
| BW22GS019616 | BORL14*2//BECARD/QUAIU #1/3/MOKUE #1/4/BORL14*2//KFA/2*KACHU | A:A | T:T |
| BW22GS019617 | CNO79//PF70354/MUS/3/PASTOR/4/BAV92*2/5/HAR311/6/BECARD/QUAIU #1/7/BECARD/QUAIU #1/8/BORL14//KFA/2*KACHU/9/ROLF07*2/DIAMONDBIRD//TRCH/HUIRIVIS #1/3/BORL14 | A:A | T:T |
| BW22GS019618 | BAJ #1/AKURI*2//HUIRIVIS #1/KBIRD*2/3/KUTZ*2//KFA/2*KACHU | A:A | T:T |
| BW22GS019619 | NADI#1*2/3/ATTILA*2/PBW65*2//MURGA*2/4/BORL14*2//KFA/2*KACHU | A:A | T:T |
| BW22GS019620 | NADI#1*2/3/ATTILA*2/PBW65*2//MURGA*2/4/BORL14*2//KFA/2*KACHU | A:A | T:T |
| BW22GS019621 | NADI#1*2/3/ATTILA*2/PBW65*2//MURGA*2/4/BORL14*2//KFA/2*KACHU | A:A | T:T |
| BW22GS019622 | KACHU//WBLL1*2/BRAMBLING*2/6/ROLF07*2/5/REH/HARE//2*BCN/3/CROC_1/AE.SQUARROSA (213)//PGO/4/HUITES*2/7/KUTZ//KFA/2*KACHU | A:A | T:T |
| BW22GS019623 | KACHU//WBLL1*2/BRAMBLING*2/6/ROLF07*2/5/REH/HARE//2*BCN/3/CROC_1/AE.SQUARROSA (213)//PGO/4/HUITES*2/7/KUTZ//KFA/2*KACHU | A:A | T:T |
| BW22GS019624 | KACHU//WBLL1*2/BRAMBLING*2/6/ROLF07*2/5/REH/HARE//2*BCN/3/CROC_1/AE.SQUARROSA (213)//PGO/4/HUITES*2/7/KUTZ//KFA/2*KACHU | A:A | T:T |
| BW22GS019625 | KACHU//WBLL1*2/BRAMBLING*2/6/ROLF07*2/5/REH/HARE//2*BCN/3/CROC_1/AE.SQUARROSA (213)//PGO/4/HUITES*2/7/KUTZ//KFA/2*KACHU | A:A | T:T |
| BW22GS019626 | KACHU//WBLL1*2/BRAMBLING*2/3/KACHU/KIRITATI/4/CIRO16/2*BORL14/5/KACHU//WBLL1*2/BRAMBLING*2/3/KACHU/KIRITATI | A:A | T:T |

| | | | |
|---|---|---|---|
| BW22GS019627 | CHIBIA//PRLII/CM65531/3/MISR 2*2/4/QUAIU/5/PBW343*2/KUKUNA*2//FRTL/PIFED/6/CHIBIA//PRLII/CM65531/3/FISCAL/4/SUP152/7/2*MOKUE #1 | A:A | T:T |
| BW22GS019628 | BLOUK #1/MUNAL/3/WBLL1*2/SHAMA//BAJ #1/4/SUP152/BAJ #1/5/MOKUE #1/6/KACHU//WBLL1*2/BRAMBLING*2/3/KACHU/KIRITATI | A:A | T:T |
| BW22GS019629 | BLOUK #1/MUNAL/3/WBLL1*2/SHAMA//BAJ #1/4/SUP152/BAJ #1/5/MOKUE #1/6/KACHU//WBLL1*2/BRAMBLING*2/3/KACHU/KIRITATI | A:A | T:T |
| BW22GS019630 | BLOUK #1/MUNAL/3/WBLL1*2/SHAMA//BAJ #1/4/SUP152/BAJ #1/5/MOKUE #1/6/KACHU//WBLL1*2/BRAMBLING*2/3/KACHU/KIRITATI | A:A | T:T |
| BW22GS019631 | SUP152/BLOUK #1/3/PRL/2*PASTOR*2//VORB/4/SUP152/BLOUK #1*2/5/BORL14*2//KFA/2*KACHU | A:A | T:T |
| BW22GS019632 | SUP152/BLOUK #1/3/PRL/2*PASTOR*2//VORB/4/SUP152/BLOUK #1*2/5/BORL14*2//KFA/2*KACHU | A:A | C:T |
| BW22GS019633 | BECARD/AKURI/3/KACHU//WBLL1*2/BRAMBLING/4/MUTUS/AKURI*2/5/MOKUE #1 | A:A | T:T |
| BW22GS019634 | KACHU//WBLL1*2/BRAMBLING/3/KACHU/KIRITATI/4/MOKUE #1/5/BORL14*2//MUNAL #1/FRANCOLIN #1 | A:A | T:T |
| BW22GS019635 | KACHU #1/3/T.DICOCCON PI94624/AE.SQUARROSA (409)//BCN/4/2*KACHU/5/MUTUS*2/TECUE #1/6/MUTUS*2/TECUE #1*2/7/MOKUE #1 | A:A | T:T |
| BW22GS019636 | KACHU #1/3/T.DICOCCON PI94624/AE.SQUARROSA (409)//BCN/4/2*KACHU/5/MUTUS*2/TECUE #1/6/MUTUS*2/TECUE #1*2/7/NELOKI*2//KACHU/KIRITATI | A:A | T:T |
| BW22GS019637 | KACHU #1/3/T.DICOCCON PI94624/AE.SQUARROSA (409)//BCN/4/2*KACHU/5/MUTUS*2/TECUE #1/6/MUTUS*2/TECUE #1*2/7/NELOKI*2//KACHU/KIRITATI | A:A | T:T |
| BW22GS019638 | KACHU #1/3/T.DICOCCON PI94624/AE.SQUARROSA (409)//BCN/4/2*KACHU/5/MUTUS*2/TECUE #1/6/MUTUS*2/TECUE #1*2/7/NELOKI*2//KACHU/KIRITATI | A:A | T:T |
| BW22GS019639 | KACHU #1/3/T.DICOCCON PI94624/AE.SQUARROSA (409)//BCN/4/2*KACHU/5/MUTUS*2/TECUE #1/6/MUTUS*2/TECUE #1*2/7/NELOKI*2//KACHU/KIRITATI | A:A | T:T |
| BW22GS019640 | SUP152/QUAIU #2//BECARD/QUAIU #1/7/ATTILA/3*BCN//BAV92/3/PASTOR/4/TACUPETO F2001*2/BRAMBLING/5/PAURAQ/6/KFA/2*KACHU/8/KFA/2*KACHU*2//MISR 1 | A:A | T:T |
| BW22GS019641 | KACHU #1/3/T.DICOCCON PI94624/AE.SQUARROSA (409)//BCN/4/2*KACHU/5/MUTUS*2/TECUE #1*2/7/ATTILA/3*BCN//BAV92/3/PASTOR/4/TACUPETO F2001*2/BRAMBLING/5/PAURAQ/6/KFA/2*KACHU | A:A | T:T |

| | | | |
|---|---|---|---|
| BW22GS019642 | ATTILA*2/PBW65*2//KACHU/3/FRNCLN*2/TECUE #1/4/KUTZ*2//KFA/2*KACHU/5/BORL14*2//MUNAL #1/FRANCOLIN #1 | A:A | T:T |
| BW22GS019643 | ATTILA*2/PBW65*2//KACHU/3/FRNCLN*2/TECUE #1*2/6/WBLL1*2/KKTS//PASTOR/KUKUNA/3/KINGBIRD #1//INQALAB 91*2/TUKURU/5/KAUZ//ALTAR 84/AOS/3/MILAN/KAUZ/4/SAUAL | A:A | T:T |
| BW22GS019644 | SUP152//WBLL1*2/BRAMBLING*2/3/KSW/SAUAL//SAUAL/4/BORL14*2//KFA/2*KACHU/5/BORL14*2//KFA/2*KACHU | A:A | T:T |
| BW22GS019645 | SUP152//WBLL1*2/BRAMBLING*2/3/KSW/SAUAL//SAUAL*2/4/PRL/2*PASTOR//KACHU | A:A | T:T |
| BW22GS019646 | SUP152//WBLL1*2/BRAMBLING*2/3/KSW/SAUAL//SAUAL*2/4/CIRO16/2*BORL14 | NA | T:T |
| BW22GS019647 | BECARD//ND643/2*WBLL1*2/3/KSW/SAUAL//SAUAL*2/4/KFA/2*KACHU*2//SUP152 | NA | T:T |
